# A proteogenomic tool uncovers protein markers for human microglial states

**DOI:** 10.1101/2025.03.31.646212

**Authors:** Verena Haage, Alex R. Bautista, John Tuddenham, Victoria Marshe, Rebecca Chiu, YunDuo Liu, Tsering Lama, Shane S. Kelly, Neelang A. Parghi, Junho Park, Alice Buonfiglioli, Julia L. Furnari, Imdadul Haq, Richard Pearse, Ronak Patel, Gizem Terzioglu, Hanane Touil, Lu Zeng, James Noble, Rani A. Sarkis, Neil A. Shneider, Lotje D. de Witte, Julie Schneider, Andrew Teich, Tracy L. Young-Pearse, Claire Riley, David A. Bennett, Peter Canoll, Jeffrey N. Bruce, Andrew J.M. Howden, Amy F. Lloyd, Bart de Strooper, Falak Sher, Andrew A. Sproul, Marta Olah, Mariko Taga, Ya Zhang, Lu Caisheng, Masashi Fujita, Vladislav A. Petyuk, Philip L. De Jager

## Abstract

Human microglial heterogeneity has been largely described using transcriptomic data. Here, we introduce a microglial proteomic data resource and a Cellular Indexing of Transcriptomes and Epitopes by Sequencing panel enhanced with antibodies targeting 17 microglial cell surface proteins (mCITE-Seq). We evaluated mCITE-Seq on HMC3 microglia-like cells, induced-pluripotent stem cell-derived microglia (iMG), and freshly isolated primary human microglia. We identified novel protein microglial markers such as CD51 and relate expression of 101 cell surface proteins to transcriptional programs. This results in the identification and validation of three protein marker combinations with which to purify microglia enriched with each of 23 transcriptional programs; for example, CD49D, HLA-DR and CD32 enrich for GPNMB^high^ (“disease associated”) microglia. Further, we identify and validate proteins - SIRPA, PDPN and CD162 – that differentiate microglia from infiltrating macrophages. The mCITE-Seq panel enables the transition from RNA-based classification and facilitates the functional characterization and harmonization of model systems.

## Introduction

Given the diverse functions of microglia in contributing to the maintenance of central nervous system (CNS) homeostasis, there has been significant interest in characterizing specialized subsets of human microglia in different brain regions, physiological or pathophysiological contexts over the past decade ^1,2^. Leveraging single-cell RNA sequencing (scRNAseq), multiple datasets that delve into the characteristics of human microglia ^1,3–11^ have played a pivotal role in reshaping our understanding of microglial heterogeneity.

The need for a shared nomenclature of human microglia is evident^12^, however current definitions are primarily based on transcriptomic data. While a few approaches to characterize human microglia at the proteomic level have been attempted and enriched our understanding ^6,13–15^, those studies were only able to provide bulk proteomic data as single-cell proteomic technologies for human microglia continue to be optimized. Such data would provide the best resource to classify functional subtypes or programs.

In the interim, we attempted to provide a new **Resource** by developing a proteogenomic tool with enhanced microglial content, enabling the simultaneous measurement of single human microglia on the transcriptomic and proteomic level. Specifically, we generated and leveraged a new human microglial proteome dataset and the existing Cellular Indexing of Transcriptomes and Epitopes by Sequencing (CITE-Seq)^16^ approach to quantitatively assess the expression of 164 cell surface proteins while capturing the transcriptome of each cell in parallel. The target proteins include 147 proteins that are present in a commercial product used in leukocyte profiling as well as 17 additional microglial cell surface proteins identified from our bulk proteomic data generated from freshly isolated human microglia.

Following the *in silico* analysis of our human microglial proteome data, candidate antibodies were selected, titrated, and then added to the commercially available TotalSeq™-A Human Universal Cocktail, V1.0 to generate a CITE-Seq cocktail enriched with antibodies targeting human microglial cell surface proteins (mCITE-Seq panel). We tested the mCITE-Seq panel on different preparations of human microglia and microglia-like cells: the HMC3 cell line, different preparations of induced-pluripotent stem cell-derived microglia (iMGs)^17–19^, peripheral blood mononuclear cells (PBMC), as well as human microglia freshly isolated from brain tissue derived from donors with Alzheimer’s disease (AD), Multiple Sclerosis (MS) or glioblastoma multiforme (GBM) (**Figure 1A**). We contrast the information in the RNA and proteomic data, and we propose a set of cell-surface protein markers that capture major transcriptional programs of human microglia that may help to define microglial states, discriminate microglia from infiltrating macrophage, and enable purification of selected microglia to facilitate functional studies.

**Figure 1.**
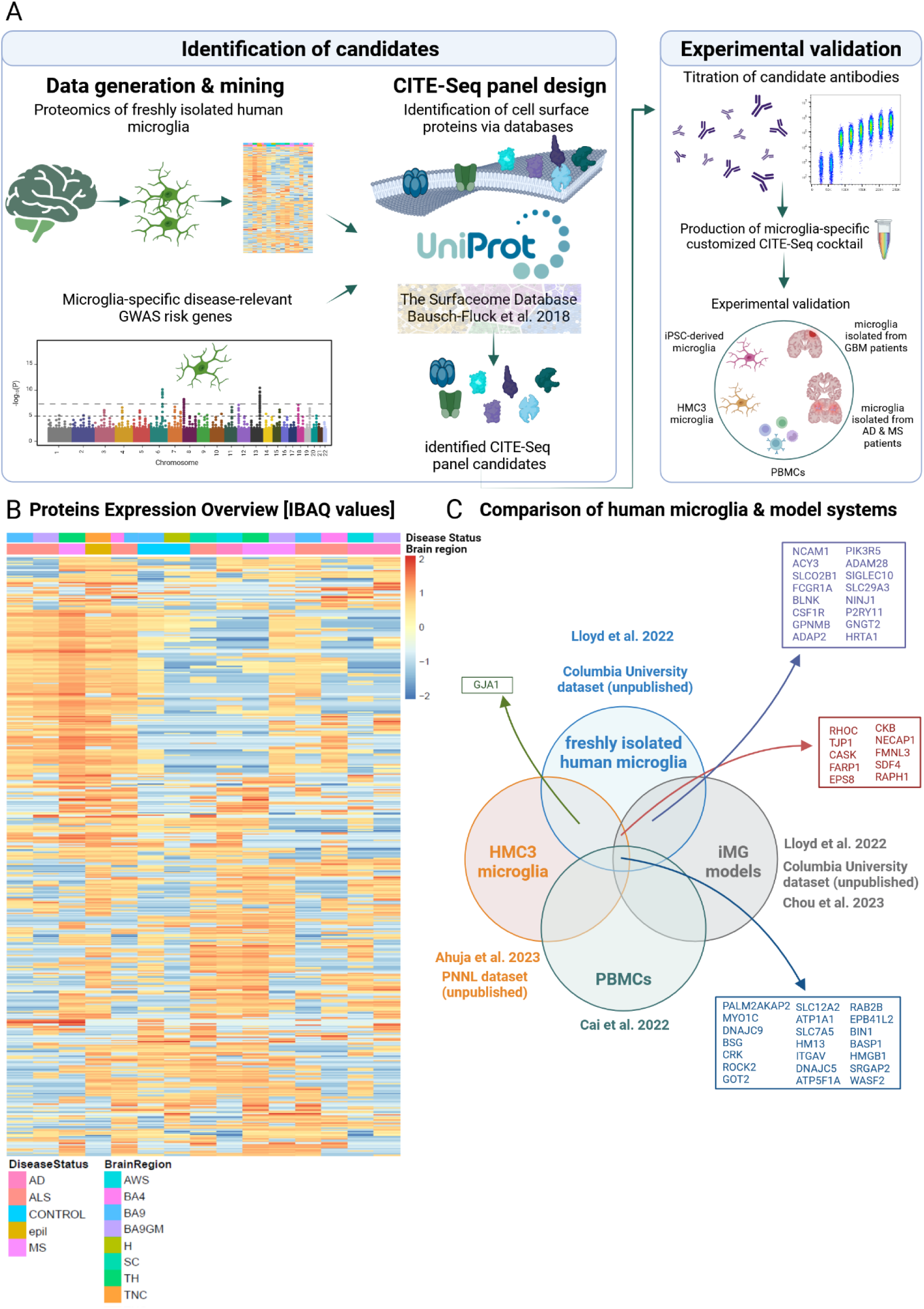
**A. Overview of the experimental approach for the design of a microglia-specific CITE-Seq panel.** The study was divided into two major parts. The first part - the identification of candidates encompassed the generation of a proteomic dataset from freshly isolated human microglia as well as a GWAS study identifying disease-relevant microglial risk genes. Using the proteomics dataset as well as the GWAS data, microglia-specific cell surface and cell membrane proteins were subsequently *in silico* identified using the databases Uniprot (https://www.uniprot.org/) and the surfaceome^25^. Double-scored candidates from both analyses were subsequently researched for antibody availability, titrated using flow cytometry and the optimal concentrations for each selected candidate were determined for the production of the customized microglia-specific CITE-Seq panel in the Total-Seq A format by BioLegend (Experimental Validation). Subsequently, the produced microglia-specific CITE-Seq cocktail was tested on different preparations of microglia, including HMC3 microglia, iPSC-derived human microglia, human microglia freshly isolated from patients with diagnosis of either glioblastoma multiforme (GBM), Alzheimer’s disease (AD), Multiple sclerosis (MS), THP-1 monocytes and PBMCs. **B. Heatmap depicting proteomics results of freshly isolated human microglia.** The heatmap depicts the proteins sorted by expression values represented in IBAQ values (top500). Each row represents a protein, each column represents one sample. Expression levels are indicated by coloring with red showing high and blue showing low expression based on IBAQ values. On top of each column, the disease status of each analyzed samples is indicated (AD (Alzheimer’s disease)= pink; ALS (Amyotrophic Lateral Sclerosis)= red; Epilepsy = brown; MS (Multiple Sclerosis)= purple) as well as the brain region the sample was collected from (AWS = Anterior watershed, turquoise; BA4 = pink; BA9 = blue; BA9_GM (Grey Matter) = lila; H (Hippocampus) = olive; SC (Superior Colliculus) = teal; TH (Thalamus) = green; TNC (trigeminal nuclear complex) = orange). **C. Venn diagram summarizing the results of the comparison of datasets of human microglia and their model systems.** For the comparison, proteins from different datasets – freshly isolated human microglia (blue; Lloyd et al. 2024^14^); HMC3 microglia (yellow; Ahuja et al. 2023^21^ and an unpublished dataset generated by our collaborators from the Pacific Northwest National Laboratory (PNNL)); PBMCs (green; Cai et al. 2022^22^)) and iPSC-derived human microglia (iMGs) (grey; Lloyd et al. 2024^14^; Chou et al. 2023^17^; an unpublished dataset generated by us and collaborators at Columbia University) - were retained if they were observed in all studies incorporated into the analysis for one model system and unobserved in all other models and their selected studies. Next, fold difference between model systems was calculated. The top protein candidates shared between and exclusively expressed by different combinations of model systems are depicted in the Venn diagram.

## Results

### Generation of a proteomic dataset derived from freshly isolated human microglia

Proteomic datasets derived from freshly isolated human microglia are still limited given the difficulty in accessing human brain tissue^6,14^. As this study aimed to develop a proteogenomic tool enriched for microglial targets to allow in-depth characterization of human microglia at the proteomic and transcriptomic level, we initially generated shotgun proteomic profiles of purified, living human microglia to create a **Resource** and identify candidate microglial proteins to incorporate into our panel (**Figure 1A**). Human microglia were isolated from postmortem tissue samples derived from 11 individuals (n= 7 women; n= 4 men) with different neurological diagnoses including Alzheimer’s disease (AD), Amyotrophic Lateral Sclerosis (ALS), Epilepsy (Epil), Multiple Sclerosis (MS) and one control individual with no neuropathologic diagnosis. Brain tissue samples were collected from different brain regions including BA4 and BA9 (cortex), AWS (Anterior Watershed area of the white matter), TH (Thalamus), SC (*superior colliculus*), H (Hippocampus) and TNC (trigeminal nucleus complex). Brain tissue was homogenized using mechanical tissue dissociation on ice, which was followed by myelin removal using Magnetic-Activated-Cell-Sorting (MACS) columns and anti-myelin magnetic beads, and finally microglia were purified from the resulting cell suspension with Fluorescence-Activated-Cell-Sorting (FACS) gating on CD11b+CD45+ live cells (7-Aminoactinomycin D negative events (7-AAD)). 5000 isolated cells/sample were subsequently snap-frozen until further processing for LC-MS/MS proteomic data capture. The demographic and clinicopathologic characteristics of the donors are presented in **Table S1**, the proteomic raw data are presented in **Table S2A**. In single-cell RNA sequencing of cells purified using the same experimental pipeline, microglia represented, on average, >95% of these cells ^20^. Thus, while our samples contain some non-microglial immune cells found in brain tissue (macrophages and other leukocytes from blood or specialized central nervous system niches), such cells represent a very small fraction of the profiled cells; their impact on the proteomic profile is likely very small but should be kept in mind as the data are evaluated.

### General analysis of microglial proteome

We identified a total of 3,658 proteins across the 15 different tissue samples derived from the 11 donors (**Figure S1A**). For downstream analyses, we used Intensity-Based Absolute Quantification (iBAQ) values for the top 500 proteins in terms of abundance, as depicted in the heatmap in **Figure 1B**. Interestingly, while the two control tissue samples show a rather low expression across the top 500 proteins, we see a quite strong expression of a module of co-expressed proteins in a subset of samples from ALS, MS or epilepsy donors, and a second module of co-expressed proteins was shared among a different subset of ALS, AD and MS samples. There is thus some clear heterogeneity among our samples, but our moderate sample size is not designed to identify associations of proteomic features with disease diagnosis. We sampled a diverse source of microglia to capture as great a diversity of the microglial proteome as possible. Plotting samples using principle components (PC) derived from the proteomic data highlight the top two dimensions of variation (**Figure S1B**). While we see some predilection of certain sample types for a region of the graph (such as ALS samples in the right lower quadrant), we will clearly need much larger studies to resolve significant differences in the microglial proteome that are related to diagnostic categories.

### Structured review of proteomic datasets from human microglia model systems

In order to identify protein markers common between different preparations of freshly isolated human microglia and human microglial model systems such as iMGs or the HMC3 cell line, we compared several proteomic datasets to our data. As input datasets, we selected several published as well as unpublished datasets (**Figure 1C**): for HMC3, we selected data derived from serum-free and serum-cultured (10% FCS) HMC3 cells^21^ and an HMC3 proteomic dataset generated for this study (**Table S2B**). For iMGs, we selected two different existing proteomic datasets^14,17^ and a dataset of iMG proteomic data generated for this study (**Table S2C**). For freshly isolated human microglia, we used a published proteomic dataset ^14^ and our generated human microglial proteomic dataset (n= 7 female donors, n= 4 male donors; **Table S2A**). As a comparator, we also included one dataset of peripheral blood mononuclear cells (PBMC)^22^, which include monocytes. Subsequently, protein intensities were converted to copy numbers by normalization to total histone intensity using “proteomic ruler”^23^. Datasets were harmonized by removing dataset-to-dataset bias calculated as the median protein-to-protein ratio to the reference dataset. To select putative cell type-specific surface protein markers we performed the following two selection steps. First, we retained only those proteins that are enriched, that is having fold difference > 1 vs the rest of the cell models. Second, only the cell membrane proteins, according to Uniprot^24^- keyword (KW-1003) annotation were retained. **Figure 1C** illustrates the extent to which the proteomic profiles overlap.

A number of proteins were shared by all datatypes, but we also identified 10 proteins exclusively shared between freshly isolated human microglia, HMC3 microglia and iMG models, including proteins such as RHOC, TJP1, CASK, FARP1 and EPS8 among others. While we only identified one protein, GJA1 (Connexin 43), that is shared between HMC3 microglia and freshly isolated human microglia, we identified a range of proteins (n=16) shared between freshly isolated microglia and the iMG models, including common microglial markers (at the RNA level) such as CSF1R, GPNMB, SIGLEC10 as well as less common markers including ADAM28, GNGT2, SLC29A3, PIK3R5 and NINJ1 (**Figure 1C**). A detailed summary of all results is presented in **Figure S1.**

### Prioritization of human microglial cell surface proteins as candidates for the development of a microglia-enriched CITE-Seq panel

In order to identify cell-surface microglial proteins, we applied the following strategy to two sets of microglial proteins: (1) those proteins measured in our primary human microglia proteome and (2) 72 proteins encoded by genes that are implicated in Alzheimer’s disease (AD) in genome-wide association studies (GWAS) and are expressed in microglia (see Methods). To identify cell surface proteins, we first queried our microglial proteome list using two publicly available databases, Uniprot^24^ annotations and the Surfaceome^25^. We ran three different search approaches with Uniprot^24^: (1) proteins under the Gene Ontology [GO] term cell surface - resulting in 9986 proteins, (2) proteins with manual assertion of being localized at the cell membrane - resulting in 733 proteins, and (3) proteins listed under subcellular location as being cell surface proteins. These 3 searches yielded a total of 156 proteins. As a second approach, cell surface proteins were identified using the Surfaceome database^25^, a public biomedical resource which can be used to filter multi-omics data to uncover cellular phenotypes and new Surfaceome markers^25^. To do so, the publicly available table_S3_surfaceome, a detailed table for the full human membranome, was downloaded and data filtered for the Surfaceome label “surface” only, yielding 2063 predicted cell surface proteins. We then identified the subset of proteins in each of the two lists of surface proteins that are present in our microglial protein list. 57 proteins were scored as membrane- or cell surface-specific in both approaches (**Figure 2A**). We repeated this process with the set of GWAS-derived proteins, which yielded 28 additional candidates annotated as cell surface proteins. We additionally added TREM1, TREM2, TSPO, SIRPB1, PLCG2 and CR1 as cell surface proteins of interest. We then combined the selected candidates which resulted in a final number of 91 distinct candidate microglial cell surface proteins (see **Table 2**; **Figure 2A**). In **Figure S2**, we illustrate the RNA expression patterns of the genes encoding these 91 proteins across the major cortical cell types based on single nucleus RNA sequence (RNAseq) data^26^ and in different microglial subtypes based on single cell RNAseq data^20^.

**Figure 2.**
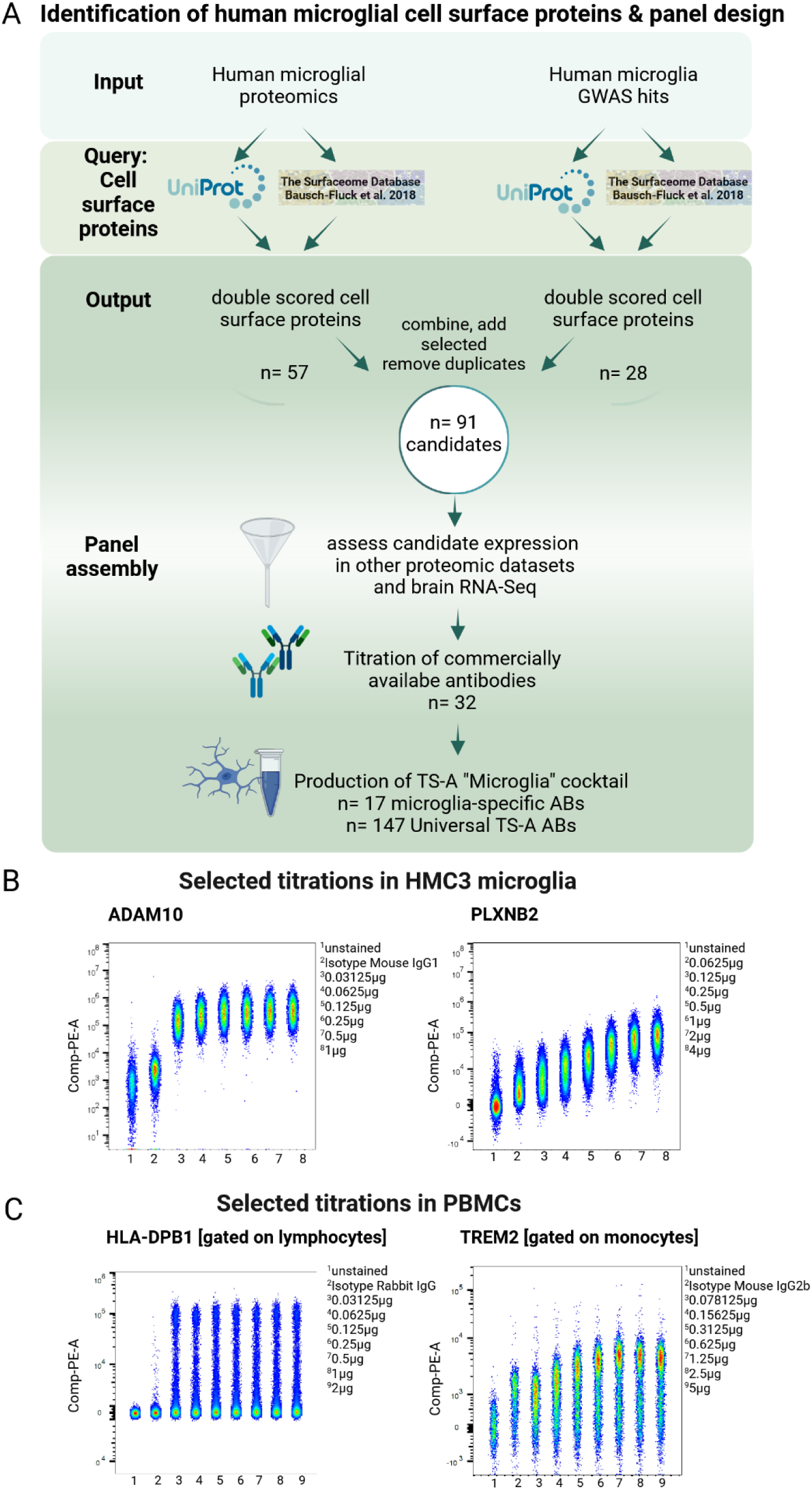
Panel building and candidate antibody titrations. **A.** Identification of human microglial cell surface proteins and panel design. For each of the datasets (generated proteomics dataset from freshly isolated human microglia, microglia-specific disease-relevant GWAS hits, *Input*), cell surface and cell membrane proteins were identified using the databases Uniprot (https://www.uniprot.org/) and the Surfaceome^25^ (*Query*). Proteins/candidates that scored in both approaches - the Uniprot and Surfaceome analysis - for the proteomics as well as the GWAS hits, were subsequently compiled into a list of n=91 initial candidates (*Output*). Candidate expression was then assessed in other available proteomics datasets of HCM3 microglia and iPSC-derived microglia as well as brain RNA-Seq datasets (n=87 candidates), commercially available antibodies were then titrated using flow cytometry (n=32) and selected candidates subsequently added to the production of the customized microglia-specific CITE-Seq panel produced in Total-Seq A (TS-A) format by BioLegend (n=17 microglia-specific antibodies, n=150 Universal cocktail 1.0 TS-A antibodies)(*Panel assembly*). **B.** Examples for titration results for HMC3 microglia. To evaluate all titration results for the antibody candidates titrated in HMC3 microglia, the PE-positive population of all samples (unstained^1^, Isotype control^2^, 1:32^3^, 1:16^4^, 1:8^5^, 1:4^6^, 1:2^7^, recommended concentration^8^) were concatenated and plotted based on their Comp-PE-A values. For visualization, ADAM10 and PLXNB2 are shown here. **C.** Examples for titration results for PBMCs. Examples for titration results for PBMCs. To evaluate all titration results for the antibody candidates titrated in PBMCs, the PE-positive population of all samples (unstained^1^, Isotype control^2^, 1:32^3^, 1:16^4^, 1:8^5^, 1:4^6^, 1:2^7^, recommended concentration^8^, doubled recommended concentration (2R)^9^) gated on lymphocytes and monocytes were concatenated and plotted based on their Comp-PE-A values. For visualization, ADAM10 and PLXNB2 are shown here.

**Table 1.** Overview of references. used for identification of microglia-enriched GWAS risk genes for Alzheimer’s disease.

| References |  |
| --- | --- |
| 1 | Beecham, G.W. *et al.* *PLoS Genet* **10**, e1004606 (2014). |
| 2 | Chung submitted |
| 3 | Corder, E.H. *et al.* *Science* **261**, 921-923 (1993). |
| 4 | Cruchaga, C. *et al.* *Nature* **505**, 550-554 (2014). |
| 5 | Cruchaga, C. *et al.* *Neuron* **78**, 256-68 (2013) |
| 6 | Cruchaga, C. *et al.* *Neuron* **78**, 256-68 (2013); Lambert, J.C. *et al.* *Nat Genet* **45**, 1452-8 (2013). |
| 7 | Deming *et al.*, *Acta Neuropath.* **133**, 839-856 |
| 8 | Deming *et al.*, *Acta Neuropath.* **133**, 839-856 (2017) |
| 9 | Escott-Price, V. *et al.* *PLoS One* **9**, e94661 (2014). |
| 10 | Sims *et al* (in press, *Nature Genetics*) |
| 11 | Goate, A. *et al.* *Nature* **349**, 704-706 (1991). |
| 12 | in preparation |
| 13 | Jakobsdottir *et al.* *PLOS Genetics* **12**(10) e1006327 (2016) |
| 14 | Jun *et al* (in press) |
| 15 | Jun, G. *et al.* *Ann Neurol* **76**, 379-92 (2014). |
| 16 | Jun, G. *et al.* *Molec Psychiatry* (2015) |
| 17 | Kohli, M.A. *et al.* *Neurology Genetics* **2**(1):e41 (2016). |
| 18 | Lambert, J.C. *et al.* *Mol Psychiatry* **18**, 461-70 (2013). |
| 19 | Lambert, J.C. *et al.* *Nat Genet* **45**, 1452-8 (2013). |
| 20 | Lambert, J.C. *et al.* *Nat Genet* **45**, 1452-8 (2013); Steinberg, S. *et al.* *Nat Genet* **47**, 445-7 (2015). |
| 21 | Levy-Lahad, E. *et al.* *Science* **269**, 970-973 (1995). |
| 22 | Logue, M.W. *et al.* *Alzheimers Dement* **10**, 609-618 e11 (2014). |
| 23 | Mez *et al.* *Alzheimer's & Dementia* **13**, 119-129 |
| 24 | Herold *et al.*, *Mol Psychiatry* **21**, 1608-1612 (2016). |
| 25 | Ruiz, A. *et al.* *Transl Psychiatry* **4**, e358 (2014) |
| 26 | Sherrington, R. *et al.* *Nature* **375**, 754-760 (1995). |
| 27 | Tosto, G. *et al.* *Ann Clin. Trans. Neurol.* (2015) |
| 28 | Wetzel-Smith, M.K. *et al.* *Nat. Med.* **20**, 1452-7 (2014). |

**Table 2.**
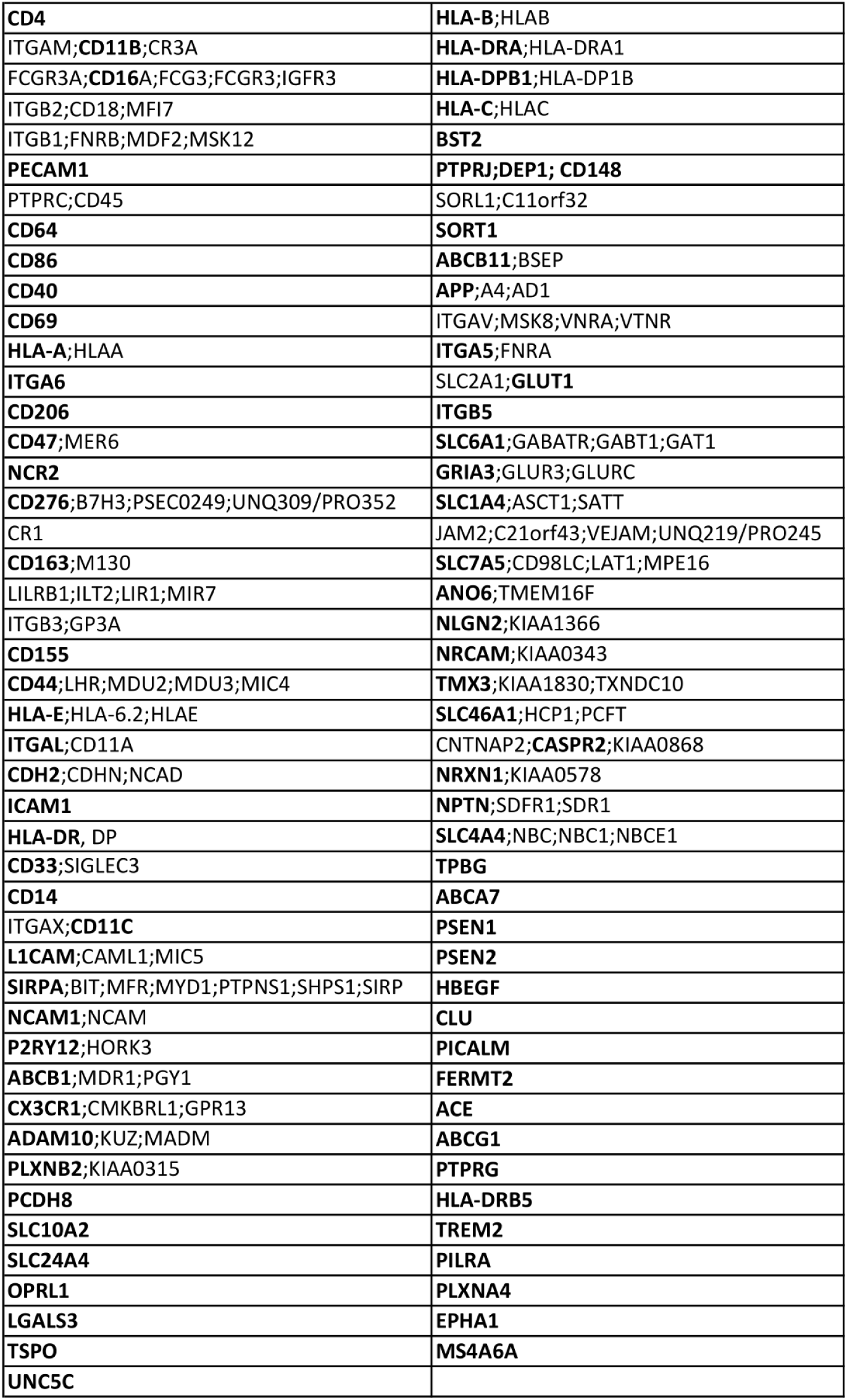
Gene names of CITE-Seq candidates.

Of the 91 candidate surface proteins, 37 are targeted by antibodies included in the Total-Seq A reagent produced by BioLegend (TotalSeqA Human Universal cocktail V.1.0). Of the remaining 54 proteins, 17 were excluded based on a lack of available antibodies. This left 37 candidate proteins that were testable (**Figure 2A**; **Table 3**); to allow for comparison of titrations across all antibodies to be tested, all antibodies were ordered in either a PE-conjugated format or, if not available, unconjugated. The latter were detected using PE-conjugated secondary antibodies.

**Table 3.**
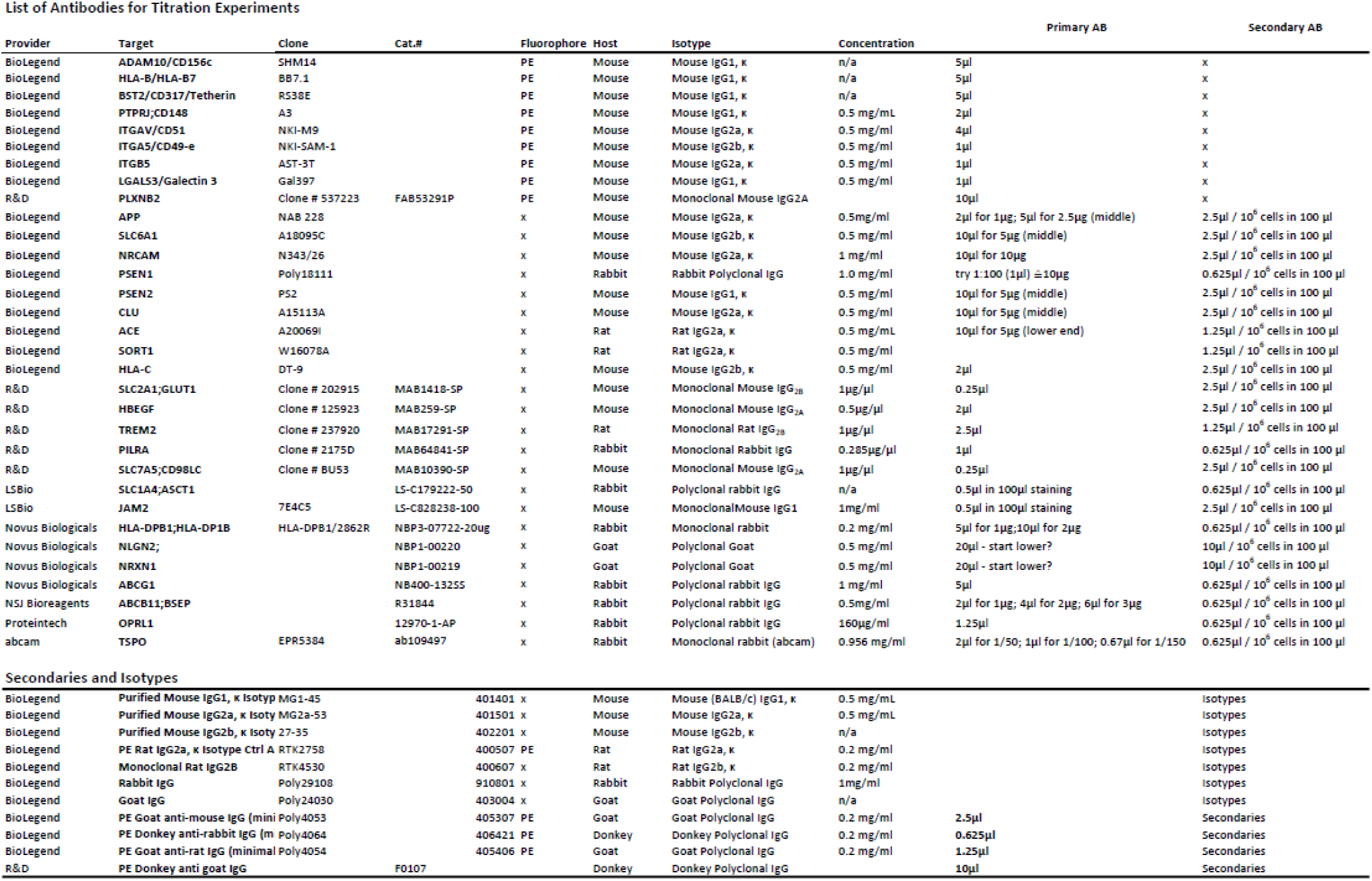
List of antibodies for titration experiments.

**Table 4.** Overview of datasets used for meta-analysis of different human microglial preparations and model systems.

| Cell type | Details | Unpublished/published | Raw data processing |
| --- | --- | --- | --- |
| HMC3 | serum-free and serum-cultured (10% FCS) HMC3 cells | Ahuja et al. 2023<sup>21</sup> | Standard FragPipe DDA-LFQ settings |
| HMC3 | untreated serum-cultured HMC3 | unpublished, generated for this study, Columbia University | Not used |
| HMC3 | Cultured HMC3 cells | unpublished, generated for this study, PNNL | Standard FragPipe TMT-16 settings |
| iMG | untreated iMGs at Day >40 of differentiation, based on protocols Abud et al.2017<sup>21</sup>; McQuade et al.2018<sup>22</sup> | Chou et al. 2023<sup>17</sup> | Standard FragPipe TMT-16 settings |
| iMG | untreated iMGs, differentiated based on Abud et al. 2017<sup>21</sup> and Bohlen et al. 2017 protocols; maintained in TIC media<sup>23</sup> | Llyod et al.2024<sup>14</sup> | Protein copy numbers used directly (Raw data unavailable) |
| iMG | untreated iMGs at Day 28-30 of differentiation, based on protocols Abud et al.2017<sup>21</sup>; McQuade et al.2018<sup>22</sup> | unpublished, generated for this study, Columbia University | Standard DIA-NN library-free settings |
| freshly isolated human microglia | Neural Tissue Dissociation Kit (P) followed by FACS- | Llyod et al.2024<sup>14</sup> | Protein copy numbers used directly (Raw |
|  | based isolation using CD11b and CD45 |  | data unavailable) |
| freshly isolated human microglia | Manual dissociation on ice followed by Percoll for myelin removal, isolated with CD11b+-MACS beads | unpublished, generated for this study, Columbia University | Standard FragPipe DDA-LFQ settings |
| Monocytes | Monocytes were isolated from blood collected from total of 76 unrelated female Caucasians (Age: $57.8 \pm 9.0$) using density gradient centrifugation. | Zeng et al. 2017<sup>61</sup> | Not used |
| Monocytes | PBMCs from healthy UK-based blood donors via NHS blood and transplant (NHSBT, Cambridge, UK) were isolated using Ficoll and monocytes sorted based on CD14+- and CD16-based AB staining. | Ravenhill et al. 2020<sup>62</sup> | Not used |
| PBMCs | PBMC were collected from 15 patients with SLE (7 males, 8 females) and 15 age-matched healthy controls (7 males, 8 females) for proteomic analysis. | Cai et al. 2022<sup>22</sup> | Standard FragPipe DDA-LFQ settings |

### Titration and selection of candidate antibodies for the microglia-specific CITE-Seq panel

Next, we titrated all antibodies. While using primary human microglia would be of interest in these titrations, such purified cells are not available in large numbers and each preparation comes from a different brain, raising concerns about comparability of results across batches. Given these practical constraints, we conducted titrations in HMC3 microglia-like cells for antibodies targeting CD49E, ADAM10, CD51, PLXNB2, PTPRJ, HLA-C, ITGB5 (for details see **Table 5**). Titrations were performed in a dilution series starting from the concentration recommended by the manufacturer following 5 serial dilutions (1:2, 1:4, 1:8, 1:16, 1:32), and the best concentration for each antibody was chosen (for an example, see **Figure 2B**). Gating schemes are depicted in **Figure S2C-D**.

**Table 5.** Overview of primary antibodies used for titration experiments.

| **Target/**<br>**Antibody** | **Supplier** | **Cat#:** | **Clone** | **PE-**<br>**conj.** | **Host**<br>**species** | **Isotype** |
| --- | --- | --- | --- | --- | --- | --- |
| ADAM10/<br>CD156c | BioLegend | 352703 | SHM14 | PE | Mouse | Mouse IgG1, κ |
| HLA-B/<br>HLA-B7 | BioLegend | 372403 | BB7.1 | PE | Mouse | Mouse IgG1, κ |
| BST2/<br>Tetherin | BioLegend | 348405 | RS38E | PE | Mouse | Mouse IgG1, κ |
| PTPRJ;<br>CD148 | BioLegend | 328708 | A3 | PE | Mouse | Mouse IgG1, κ |
| ITGAV/<br>CD51 | BioLegend | 327909 | NKI-M9 | PE | Mouse | Mouse IgG2a, κ |
| ITGA5/<br>CD49-e | BioLegend | 328009 | NKI-SAM-1 | PE | Mouse | Mouse IgG2b, κ |
| ITGB5 | BioLegend | 345203 | AST-3T | PE | Mouse | Mouse IgG2a, κ |
| LGALS3/<br>Galectin 3 | BioLegend | 126706 | Gal397 | PE | Mouse | Mouse IgG1, κ |
| PLXNB2 | Miltenyi | 130-095-212 | REA626 | PE | Human<br>recombinant | n/a |
| APP | BioLegend | 835104 | NAB 228 | n/a | Mouse | Mouse IgG2a, κ |
| SLC6A1 | BioLegend | 867401 | A18095C | n/a | Mouse | Mouse IgG2b, κ |
| NRCAM | BioLegend | 821602 | N343/26 | n/a | Mouse | Mouse IgG2a, κ |
| PSEN1 | BioLegend | 811101 | Poly18111 | n/a | Rabbit | Rabbit<br>Polyclonal IgG |
| PSEN2 | BioLegend | 814203 | PS2 | n/a | Mouse | Mouse IgG1, κ |
| CLU | BioLegend | 848701 | A15113A | n/a | Mouse | Mouse IgG2a, κ |
| ACE | BioLegend | 375801 | A20069I | n/a | Rat | Rat IgG2a, κ |
| SORT1 | BioLegend | 852101 | W16078A | n/a | Rat | Rat IgG2a, κ |
| HLA-C | BioLegend | 373302 | DT-9 | n/a | Mouse | Mouse IgG2b, κ |
| SLC2A1;<br>GLUT1 | R&D | MAB1418-SP | 202915 | n/a | Mouse | Monoclonal<br>Mouse IgG2B |
| HBEGF | R&D | MAB259-SP | 125923 | n/a | Mouse | Monoclonal<br>Mouse IgG2A |
| TREM2 | BioLegend | _6E9 | 824801 | n/a | Rat | Rat IgG2a, $\kappa$ Isotype/Monoclonal Rat IgG2B |
| TREM2 | R&D | MAB17291-SP | 237920 | n/a | Rat | Monoclonal Rat IgG2B |
| PILRA | R&D | MAB10390-SP | 2175D | n/a | Rabbit | Recombinant Monoclonal Rabbit IgG |
| SLC7A5; CD98LC | R&D | MAB10390-SP | BU53 | n/a | Mouse | Monoclonal Mouse IgG2A |
| SLC1A4; ASCT1 | LSBio | LS-C179222-50 | n/a | n/a | Rabbit | Polyclonal rabbit IgG |
| HLA-DPB1; HLA-DP1B | Novus Biologicals | NBP3-07722-20ug | HLA-DPB1/2862 R | n/a | Rabbit | Monoclonal rabbit |
| NLGN2 | Novus Biologicals | NBP1-00220 | n/a | n/a | Goat | Polyclonal Goat |
| ABCG1 | Novus Biologicals | NB400-132SS | n/a | n/a | Rabbit | Polyclonal rabbit IgG |
| ABCB11;BSEP | NSJ Bioreagents | R31844 | n/a | n/a | Rabbit | Polyclonal rabbit IgG |
| OPRL1 | Proteintech | 12970-1-AP | n/a | n/a | Rabbit | Polyclonal rabbit IgG |
| TSPO/PBR | Thermo Fisher | MA5-31966 | n/a | n/a | Rabbit | Monoclonal recombinant rabbit |
| NRXN1 | Novus Biologicals | NBP1-00219. C2 | n/a | n/a | Goat | Polyclonal Goat |
| JAM2 | LSBio | LS-C828238-100 | 7E4C5 | n/a | Mouse | Monoclonal Mouse IgG1 |

**Table 6.**
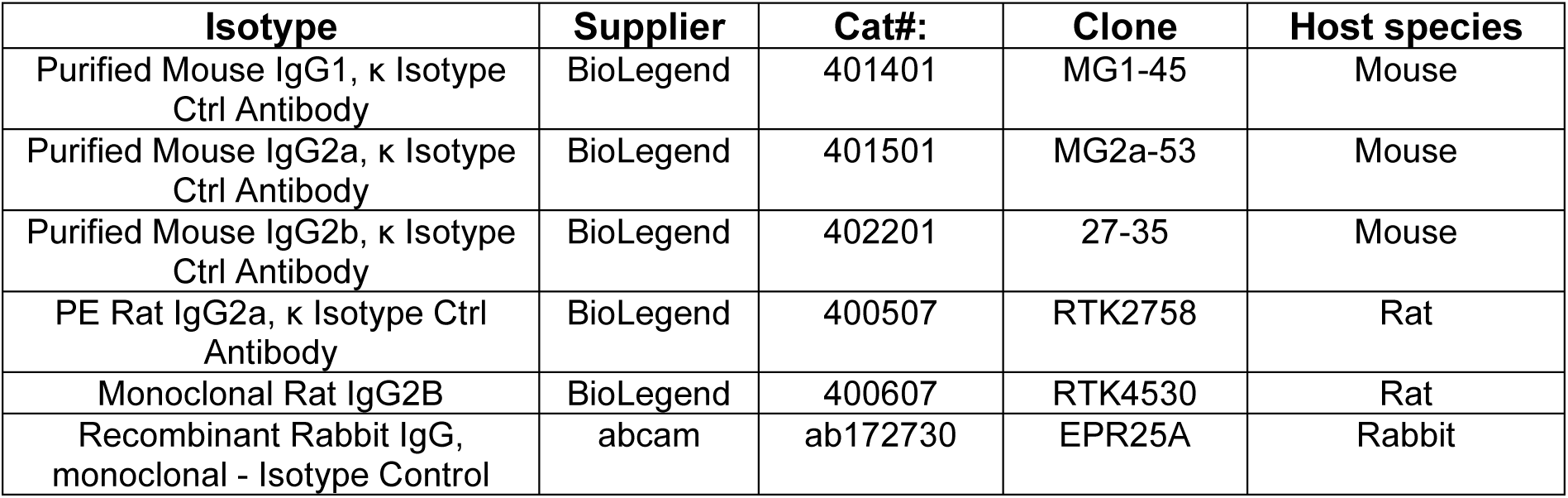
A. Overview of isotype antibodies used for titration experiments.

**Table 6.**
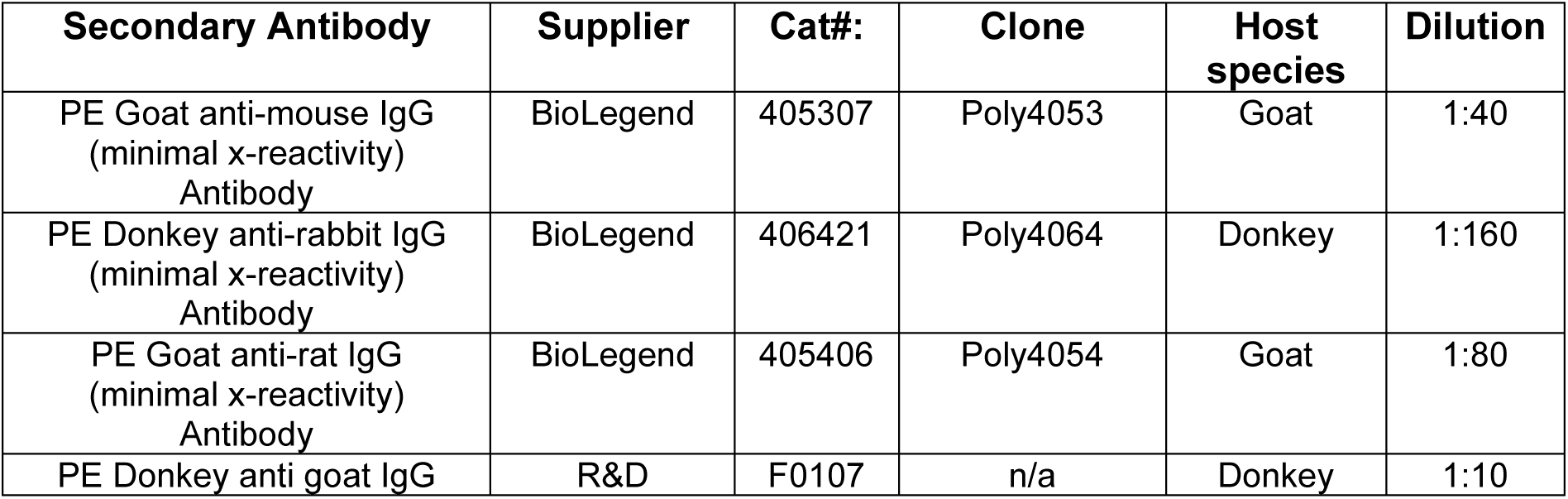
B. Overview of secondary antibodies used for titration experiments.

Certain target candidates were either not expressed on HMC3 microglia-like cells (**Figure S2A-B; Table S2B**) or not detected as a result of our initial titrations: ABCG1, HLA-B7, BST2 (see **Table 5**). Thus, we next performed titrations using PBMCs since they include monocytes, which share many expression patterns with microglia and have been used to model human microglia in functional genomic studies^27–29^. A detailed description of our stepwise evaluation of titrations is presented in the **Methods** section and results of all titrations are presented in **Table 7**.

**Table 7.** Summary table of all titration results.

| Target | Detected in HMC3 | Titred in HMC3 | Detected in PBMCs | Titred in PBMCs | Vendor | Cat# |
| --- | --- | --- | --- | --- | --- | --- |
| ABCB11 | x | x | ✓ | ✓ | NSJ Bioreagents | R31844 |
| ABCG1 | ✓ | x | ✓ | ✓ | Novus Biologicals | NB400-132SS |
| ACE2 | x | x | ✓ | ✓ | BioLegend | 375801 |
| ADAM10 | ✓ | ✓ | n/a | x | BioLegend | 352703 |
| APP | x | x | ≤1% | x | BioLegend | 835104 |
| BST2 | ✓ | x | ✓ | ✓ | BioLegend | 348405 |
| CD49e | ✓ | ✓ | n/a | x | BioLegend | 328009 |
| CD51 | ✓ | ✓ | n/a | x | BioLegend | 327909 |
| CLU | x | x | ✓ | ✓ | BioLegend | 848701 |
| GLUT1 | x | x | ✓ | ✓ | R&D | MAB1418-SP |
| HBEGF | x | x | ≤1% | x | R&D | MAB259-SP |
| HLA-B7 | ✓ | x | ≤1% | x | BioLegend | 372403 |
| HLA-C | ✓ | ✓ | n/a | x | BioLegend | 373302 |
| HLA-DPB1 | n/a | n/a | ✓ | ✓ | Novus Biologicals | NBP3-07722 |
| ITGB5 | ✓ | ✓ | n/a | x | BioLegend | 345203 |
| JAM2 | x | x | ≤1% | x | LSBio | LS-C828238-100 |
| LGALS3 | x | x | ≤1% | x | BioLegend | 126706 |
| NLGN2 | x | x | ✓ | ✓ | Novus Biologicals | NBP1-00220 |
| NRCAM | x | x | ✓ | ✓ | BioLegend | 821602 |
| NRXN1 | x | x | ≤1% | x | Novus Biologicals | NBP1-00219 |
| OPRL1 | x | x | ✓ | ✓ | Proteintech | 12970-1-AP |
| PILRA | x | x | ✓ | ✓ | R&D | MAB64841-SP |
| PLXNB2 | ✓ | ✓ | n/a | x | Miltenyi | 130-095-212 |
| PSEN1 | x | x | ≤1% | ✓ | BioLegend | 811101 |
| PSEN2 | x | x | ≤1% | ✓ | BioLegend | 814203 |
| PRPTJ | ✓ | ✓ | n/a | x | BioLegend | 328708 |
| SLC1A4 | x | x | ✓ | ✓ | LSBio | LS-C179222-50 |
| SLC6A1 | x | x | ≤1% | x | BioLegend | 867401 |
| SLC7A5 | x | x | ✓ | ✓ | R&D | MAB10390-SP |
| SORT1 | x | x | ✓ | ✓ | BioLegend | 852101 |
| TREM2 | x | x | ✓ | ✓ | R&D/BioLegend | MAB17291-SP/824801 |
| **TSPO** | x | x | ✓ | ✓ | Abcam/LSBio | ab109497/MA5-31966 |

As a result of these titration experiments, we selected 17 new antibodies that were incorporated into a customized microglia-targeting CITE-Seq antibody pool to target the following proteins: CD49E, ADAM10, CD51, PTPRJ, HLA-C, ITGB5, BST2, TREM2, CLU, NRCAM, PSEN1, PSEN2, ACE, SORT1, P2RY12 and PLXNB2 (**Table 8**). Several of these antibodies have not previously been tested for flow cytometry experiments and appear to work reliably for this application: CLU, NRCAM, PSEN1, ACE2, PSEN2, SORT1 (see **Tables 5&7**). We therefore expand the toolkit of antibody reagents for microglia with new tools that can be used for future flow cytometry applications, such as the development of a microglia-specific flow cytometry panel. For the following validation experiments, the custom panel of 17 antibodies was reconstituted and added to the TS-A Human Universal Cocktail V1.0 from BioLegend, resulting in the microglial-enhanced CITE-Seq (mCITE-Seq) panel, with the ability to detect expression of up to 164 cell surface proteins.

**Table 8.**
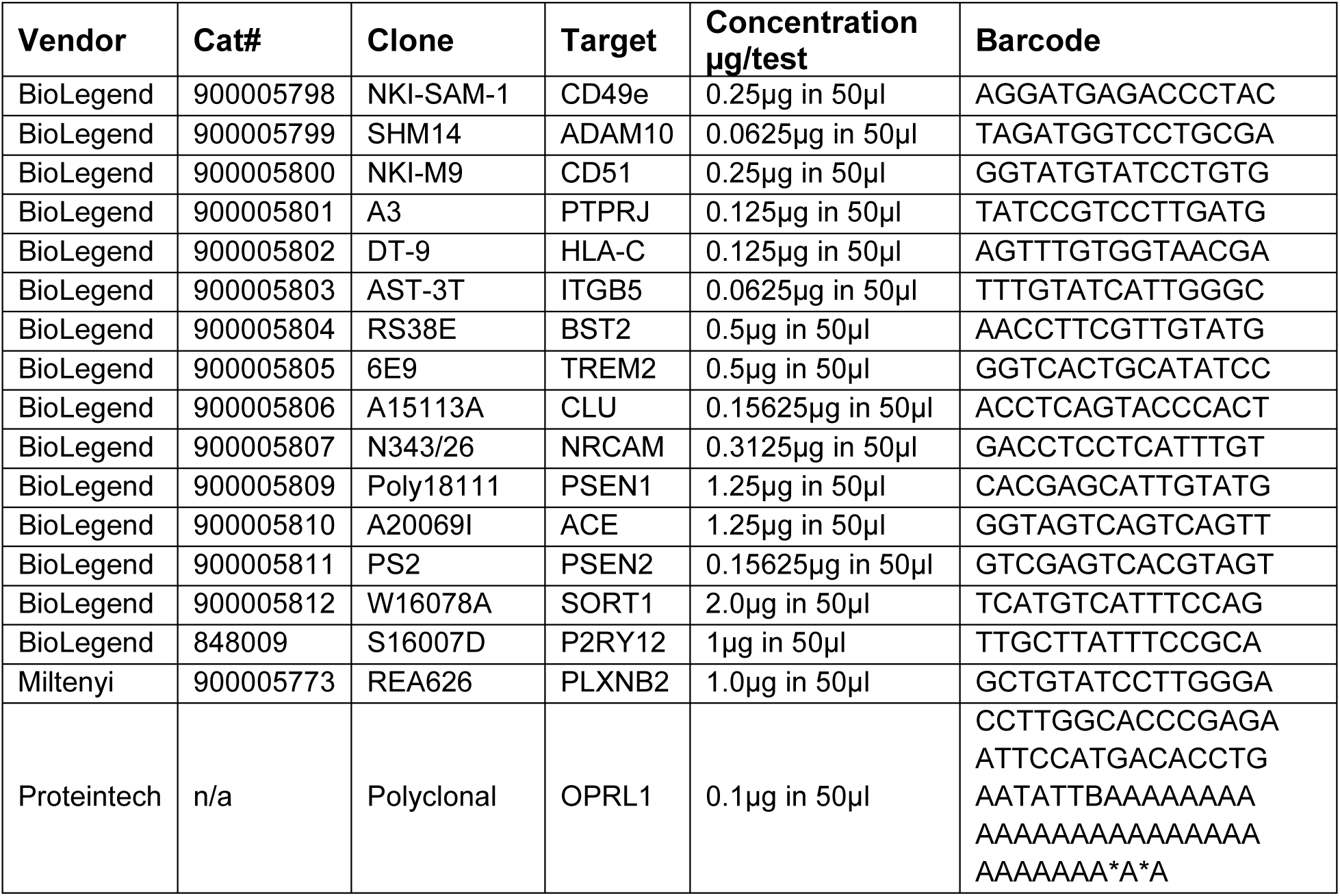
Microglia-specific markers selected for custom-conjugation in TotalSeq A format.

**Table 9.** Overview of cells/samples before and after QC.

| **Sample Name** | **n_pre_QC** | **n_post_QC** |
| --- | --- | --- |
| HMC3 | 12866 | 8921 |
| PBMCs | 15463 |  |
| iMG I | 14048 | 9125 |
| iMG II | 7689 | 6296 |
| iMG III | 6753 | 6285 |
| GBM I | 14336 | 10826 |
| GBM II | 5938 | 2482 |
| GBM III | 524 | 143 |
| MS I | 1647 | 862 |
| MS II | 966 | 28 |
| MS III | 575 | 354 |
| MS IV | 1557 | 1168 |
| AD | 1872 | 971 |

### Assessment of the microglial-enhanced CITE-Seq panel on different preparations of human microglia and microglia-like cells

We then tested our mCITE-Seq cocktail on different human microglia-like model systems: the HMC3 cell line (n= 8,921 cells) and three different preparations of iPSC-derived microglia-like (iMG) cells from different laboratories (iMG I: n= 9,125 cells; iMG II: n= 6,296 cells; iMG III: n=6,285 cells). We also evaluated freshly isolated myeloid cells from glioblastoma multiforme resections (GBM I: n= 10,826, GBM II: n= 2,482; GBM III: n= 143 cells) and freshly isolated microglia from postmortem tissue of donors diagnosed with Multiple Sclerosis (MS; MS I: n= 862 ; MS II: n= 28; MS III: n= 354;MS IV: n= 1,168) or Alzheimer’s disease (AD; n= 971 cells). We also profiled PBMCs as a control; we first clustered PBMCs (n= 15,463 cells) based on antibody expression levels, which resulted in 5 main clusters, including B cells, monocytes, Natural Killer cells, CD8+ and CD4 T cells (**Figure S3A-B**), suggesting that our novel mCITE-Seq cocktail is functioning as expected. For downstream analyses, we then computed a PBMC-based expression threshold and applied it to the microglia data to exclude proteins that were not expressed in microglia (see **Methods**). The 17 new antibodies in our microglia panel were retained in the downstream analysis irrespective of their expression levels.

We then performed clustering based on RNA expression levels (**Figure 3A**), resulting in distinct clusters for HMC3 cells and iMGs, while cells from the GBM I and II samples clustered together, distinct from GBM III, a sample with a low cell number and evidence of necrosis on pathologic examination. The postmortem MS and AD samples clustered together. We then separately performed hierarchical clustering based on the expression of the 101 target proteins or “Antibody-Derived Tags” (ADTs) left after our pre-processing and quality control pipeline (see **Methods**; **Figure 3B**). ADT-based clustering resulted in a refined version of the RNA-based clusters, characterized by more discrete and coherent clusters for HMC3 microglia-like cells, all of the iMG preparations and a clear separation of the GBM I and GBM II samples. MS and AD samples still clustered together; in this version, they clustered with the GBM III sample that has few cells. In order to see whether and to what extent RNA- vs. ADT-based clustering would distinguish the different microglia/myeloid cell preparations, we performed hierarchical clustering for HMC3 microglia-like cells, iMG I-III and freshly isolated human microglia/myeloid cell samples (AD, MS I-IV, GBM I-III) (**Figure 3C-D**). While RNA-based clustering yielded three major clusters, there was still some overlap between them (**Figure 3C**). ADT-based clustering in contrast, clearly separated HMC3 microglia-like cells from iMGs and freshly isolated human microglia/myeloid cells (**Figure 3D**). The iMG cluster contained a small fraction of freshly isolated human microglia/myeloid cells (1.89%), suggesting that the *in vitro* system may capture a specific subtype of microglia at baseline. The cluster of freshly isolated human microglia/myeloid cells also contained a small portion of iMG cells (6.56%) and HMC3 cells (0.58%), suggesting that a portion of iMGs and HMC3 microglia-like cells closely resembles freshly isolated human microglia/myeloid cells at the protein level (**Figure S3D-E**). While these subsets of iMG and HMC3 cells represent a small minority of the cell cultures, they suggest that the *in vitro* systems are capable of capturing the *ex vivo* states but are not yet optimal as most of the cells are in a distinct state that is uncommon in *ex vivo* samples.

**Figure 3.**
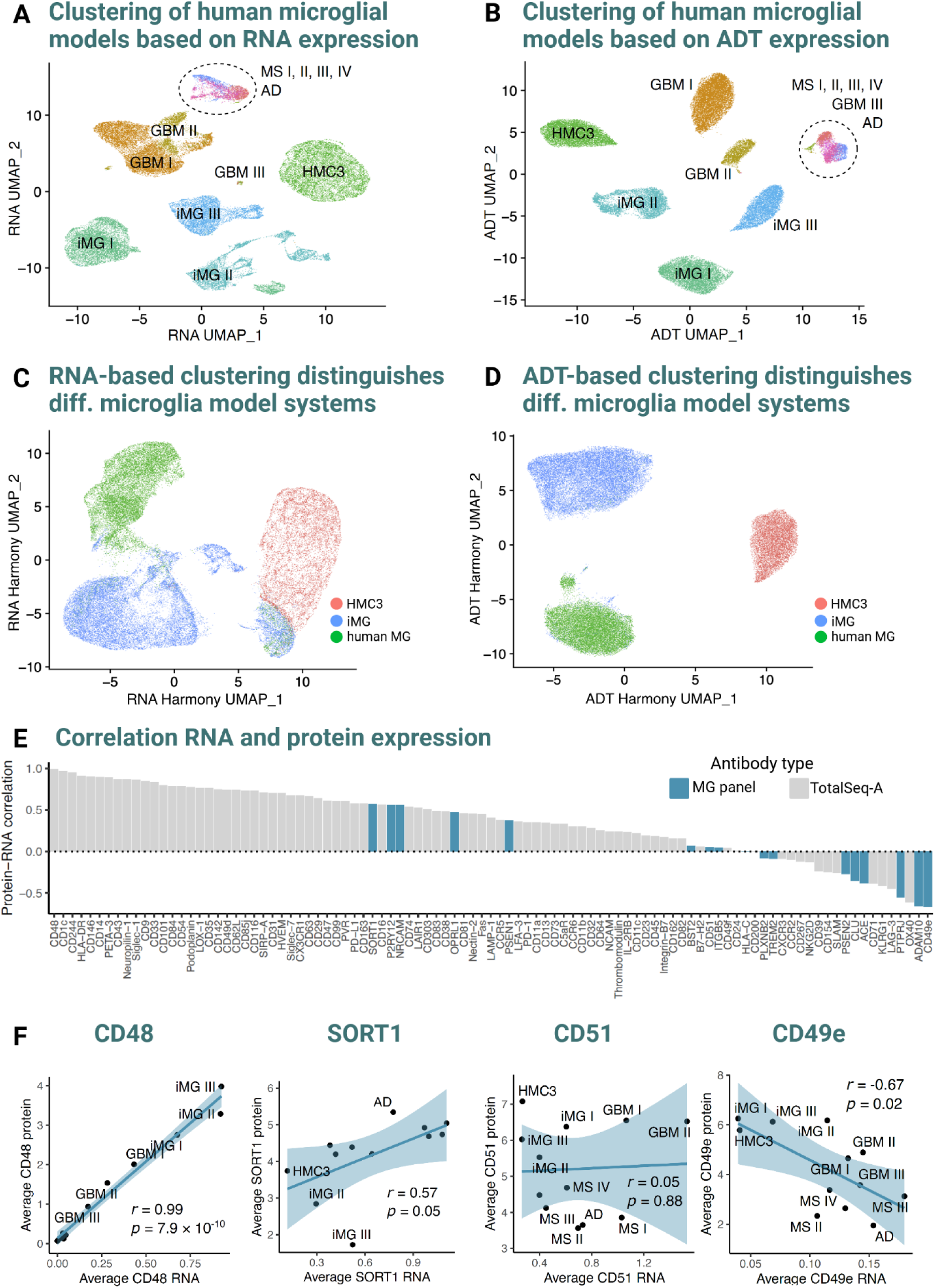
Microglia-specific CITE-Seq panel validation in human postmortem and surgical tissue, HMC3 microglia, iPSC-derived microglia and PBMCs. A.RNA-based UMAP of freshly isolated human microglia and their model systems. Following sc-RNA-seq, clustering of cells was performed based on RNA expression and plotted in a two-dimensional UMAP plot. iPSC-derived human microglia (iMG) I-III: green, teal, blue; HMC3 microglia: light green; microglia isolated from surgical tissue of GBM (Glioblastoma multiforme) patients I-III (dark green, brown, khaki); microglia freshly isolated from postmortem human brain tissue of AD (Alzheimer’s disease) and MS (Multiple Sclerosis) I-IV patients: blue, violet, pink. **B. ADT-based UMAP of freshly isolated human microglia and their model systems.** Following sc-RNA-seq, clustering of cells was performed based on ADT (Antibody-Derived Tag) expression and plotted in a two-dimensional UMAP plot. iPSC-derived human microglia (iMGs) I-III: green, teal, blue; HMC3 microglia: light green; microglia isolated from surgical tissue of GBM (Glioblastoma multiforme) patients I-III (dark green, brown, khaki); microglia freshly isolated from postmortem human brain tissue of AD (Alzheimer’s disease) and MS (Multiple Sclerosis) I-IV patients: blue, violet, pink. **C. RNA-based clustering distinguishes human microglia and their model systems.** Hierarchical clustering of cells based on their transcriptome results in three distinct clusters, namely HMC3 microglia (red), iMGs (iPSC-derived human microglia; blue) and freshly isolated human microglia from surgical and postmortem brain tissue samples (green). **D. ADT-based clustering distinguishes human microglia and their model systems.** Hierarchical clustering of cells based on their transcriptome results in three distinct clusters, namely HMC3 microglia (red), iMG (iPSC-derived human microglia; blue) and freshly isolated human microglia from surgical and postmortem brain tissue samples (green). **E. Correlation of RNA and protein expression across all samples analyzed.** Plot shows the correlation between the 101 ADTs integrated into the final analysis depicted on the x-axis and the Protein-RNA correlation coefficient ranging from -1 (negative correlation) to +1 (positive correlation) on the y-axis. Microglia-specific ADTs are marked in turquoise, Universal TS-A ADTs are marked in grey. **F. Correlation of RNA and protein expression for selected markers.** Correlation plots depict average RNA vs. average protein correlation for the selected markers CD48, SORT1, CD51 and CD49e. Black dots indicate correlation data for the samples included in the analysis (AD, MS I-IV, HMC3, PBMCs, iMG I-III), while r indicates the calculated correlation coefficient for each of the selected markers.

We then correlated RNA and protein expression for the 101 analyzed ADTs across all of the analyzed samples (**Figure 3E**). For most candidate proteins we found a positive correlation with RNA expression, including six of our custom microglial markers (P2RY12, SORT1, NRCAM, PSEN1, OPRL1, CD51). Correlations for CD48, which showed the highest correlation between RNA and protein expression (r= 0.99) as well as three selected microglial proteins are highlighted in **Figure 3F**: SORT1 (r= 0.57), CD51 (r= 0.51), and CLU (r= -0.37). The inverse correlation is intriguing and is also seen with a number of the antibodies in the standard TS-A Human Universal Cocktail; it may relate to the relative dynamics of transcription, translation and half-lives of the respective transcripts and proteins.

Next, we assessed the expression of the 17 target proteins in our custom pool in more detail. **Figure 4A** depicts a heatmap summarizing the expression of the 17 targeted microglia-enriched proteins across the different model systems and samples that we assessed. AD and MS samples seemed to specifically express high levels of CLU, PSEN1, PSEN2, ACE, BST2, OPRL1, P2RY12, SORT1, NRCAM and TREM2 (microglial cocktail) in addition to CD74, CD23, and CD303 (TS-A Human Universal cocktail V1.0) when assessing the 101 ADTs (**Figure S4A, Table S3**). Focusing on the GBM I and II samples which have a large number of sequenced cells, tumor-derived cells showed a high expression in PLXNB2, ITGB5 and CD51 (microglial cocktail) in addition to CD64, HLA-DR, CD9 and CD11c (from the TS-A Human Universal cocktail V1.0)(**Figure S4A, Table S3**). Consistent with principal component analysis, GBM III cells had a different expression pattern with regards to these microglial markers (PLXNB2, ITGB5 and CD51), clustering with the freshly isolated human microglia/myeloid cells isolated from postmortem brain tissue (AD, MS I-IV).

**Figure 4.**
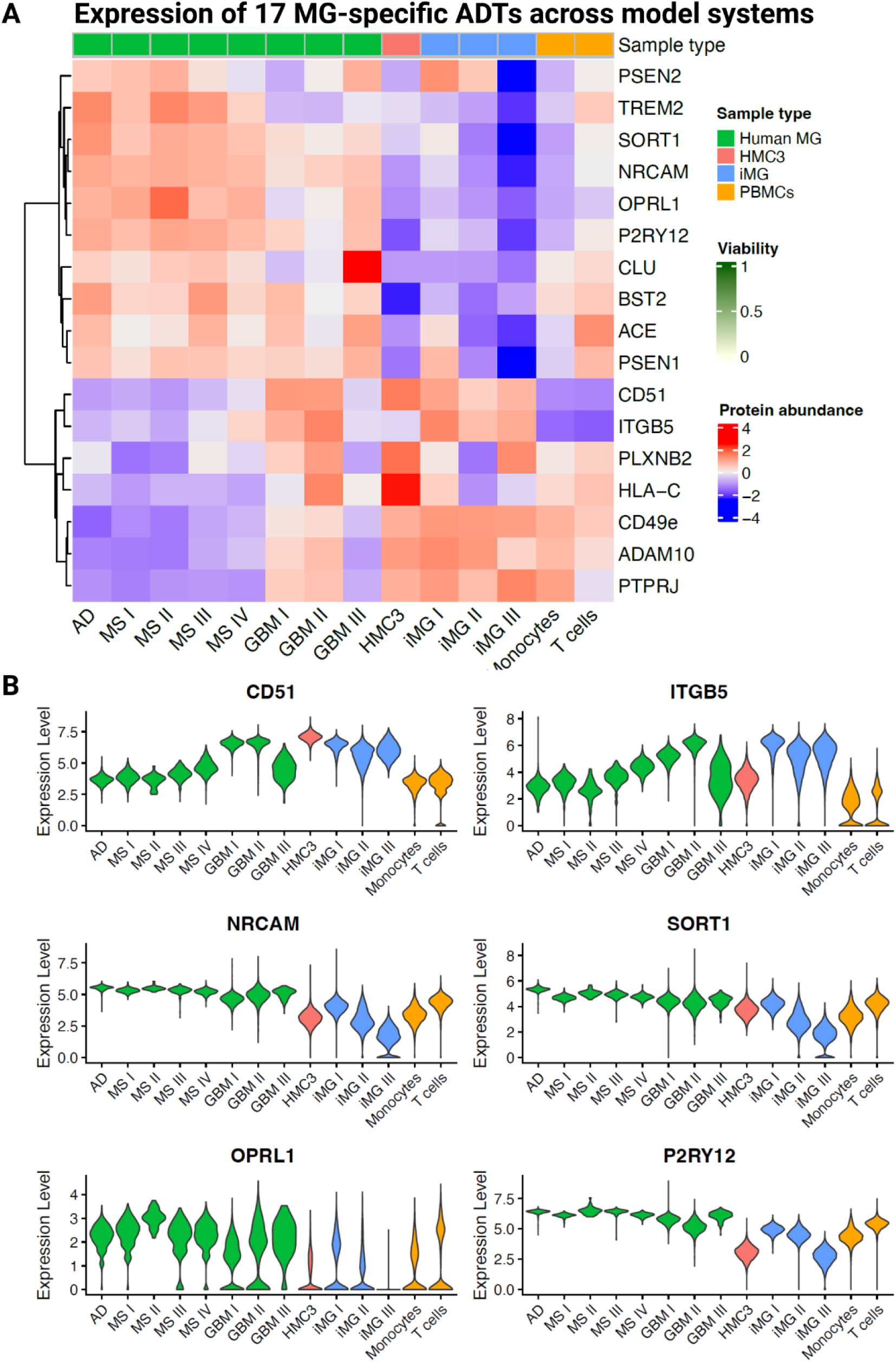
Expression of microglia-specific CITE-Seq candidates across different preparations of freshly isolated human microglia and microglia model systems. A. Heatmap depicting the expression of microglia-specific ADTs (Antibody-derived tags) across different samples of human microglia and model systems. Rows depict the different antibody candidates while columns depict the different samples included in the analysis, including freshly isolated microglia from postmortem samples of an AD (Alzheimer’s disease) patient, MS patients I-IV (Multiple sclerosis), microglia isolated from surgical tissue derived from three GBM (Glioblastoma multiforme) patients, HMC3 microglia, three different preparations of iPSC-derived human microglia (iMG) from different laboratories and PBMCs derived from 4 different individuals (here depicted are only data for monocytes and T cells). Cell viability in % ranging from 0% (white) to 100% (green) is indicated on top of each sample together with classification of the sample type (green: freshly isolated human microglia samples; red: HMC3 microglia; blue: iPSC-derived human microglia; yellow: PBMCs, extracted are data for monocytes and T cells). **B. Expression of selected microglia-specific CITE-Seq antibodies (ADTs) across sample types.** Violin plots depict the expression levels of selected microglia-specific marker proteins based on quantified ADT expression (CD51, NRCAM, OPRL1, ITGB5, SORT1, P2RY12) across the different sample types (green: freshly isolated human microglia samples – AD, MSI-IV; red: HMC3 microglia; blue: iPSC-derived human microglia, iMG I-III; yellow: PBMCs, extracted are data for monocytes and T cells).

To address the microglial specificity of our new markers, we evaluated the PBMC-derived cells: monocytes showed a low level of expression of PTPRJ, CD49E, ADAM10 , HLA-C, PLXNB2, while T cells had strong expression of ACE, followed by HLA-C, PSEN1 and CD49e and low expression of a few other markers. Overall, many of the 17 markers were expressed at low levels or were absent in monocytes and T-cells, strengthening their validity as microglia-enriched markers (**Figure 4A**, **Figure S4A**). As a result of these analyses, P2RY12, SORT1, ITGB5, CD51, OPRL1, NRCAM emerged as the strongest candidates for microglia-enriched markers (**Figure 4B; Figure S4)**. CD51 and ITGB5 are highly expressed by microglia/myeloid cells isolated from brain tumors, HMC3 and iMG microglia-like cells. While the expression of NRCAM, OPRL1, SORT1 and P2RY12 expression is clearly highest in freshly isolated human microglia samples (AD, MS, GBM), positioning them as interesting and strong protein candidates for future studies.

### Identification of HLA-DR and CD51 as protein markers for GPNMB^high^ and OxPhos-1-like human microglia

To assess whether we could identify protein markers for defined human microglial transcriptional programs, we projected the highest quality subset of the mCITEseq data (GBM I & II samples derived from surgical tissue) into two different models derived from single cell RNA sequencing (scRNAseq) data derived from purified living microglia^20,30^. First, we accessed a recently compiled model of microglial transcriptional programs built from a scRNAseq resource of 441,088 live microglia extracted from 13 different brain regions of 161 individuals with a variety of neurodegenerative disease diagnoses^30^. This report presents a model that decomposes the microglial transcriptome into 23 transcriptional programs using a dimensionality-reduction approach (single-cell Hierarchical Poisson Factorization, scHPF^31^), and the programs are referred to as “factors”, with each microglia having a value for each factor. Most of these programs can be clearly annotated functionally, with two of them capturing the disease associated microglia-like (DAM) signature originally identified in mice^32^. For the subsequent analyses, we focused on the GBM I and GBM II samples as they constituted the best primary microglia mCITE-Seq data. We projected 10,826 cells for GBM I and 2,482 cells for GBM II into the scHPF model (**Figure S5A**). Due to the nature of brain tumor (GBM) samples and CD11b-based magnetic-associated cell sorting (MACS) isolation of cells for subsequent CITE-Seq experiments, we expected a mixture of microglia and infiltrating macrophages. When assessing hierarchical clustering analysis of the joint cells from GBM I and GBM II, we identified two bigger clusters (0,1) and one smaller cluster (2) (**Figure 5A**). We assessed marker expression across the three identified clusters using two different marker sets previously used for distinguishing microglia from peripheral macrophages/infiltrated monocytes^33,10^ (**Figure 5B**), identifying cluster 1 as microglia, characterized by the high expression of *TREM2*, *GPR34*, *P2RY12*, *TMEM119*, *SALL1*, *SLC2A5*. Further, we identified cluster 0 as macrophages, characterized by the high expression of *OLFML3*, *HEXB*, *EMILIN2*, *MS4A7*, *LYZ*, *CD163*. Cluster 2 expressed high levels of *PCNA* and *MKI67*, indicating a highly proliferative state of myeloid cells. We therefore proceeded to use cluster 1, the microglial cluster, for downstream analyses. Evaluating the 23 transcriptional programs, we found that tumor-derived microglia have the highest expression of the OxPhos-1 (DAM1) and a GPNMB^high^ (DAM2) expression program (**Figure 5C**)^30^.

**Figure 5.**
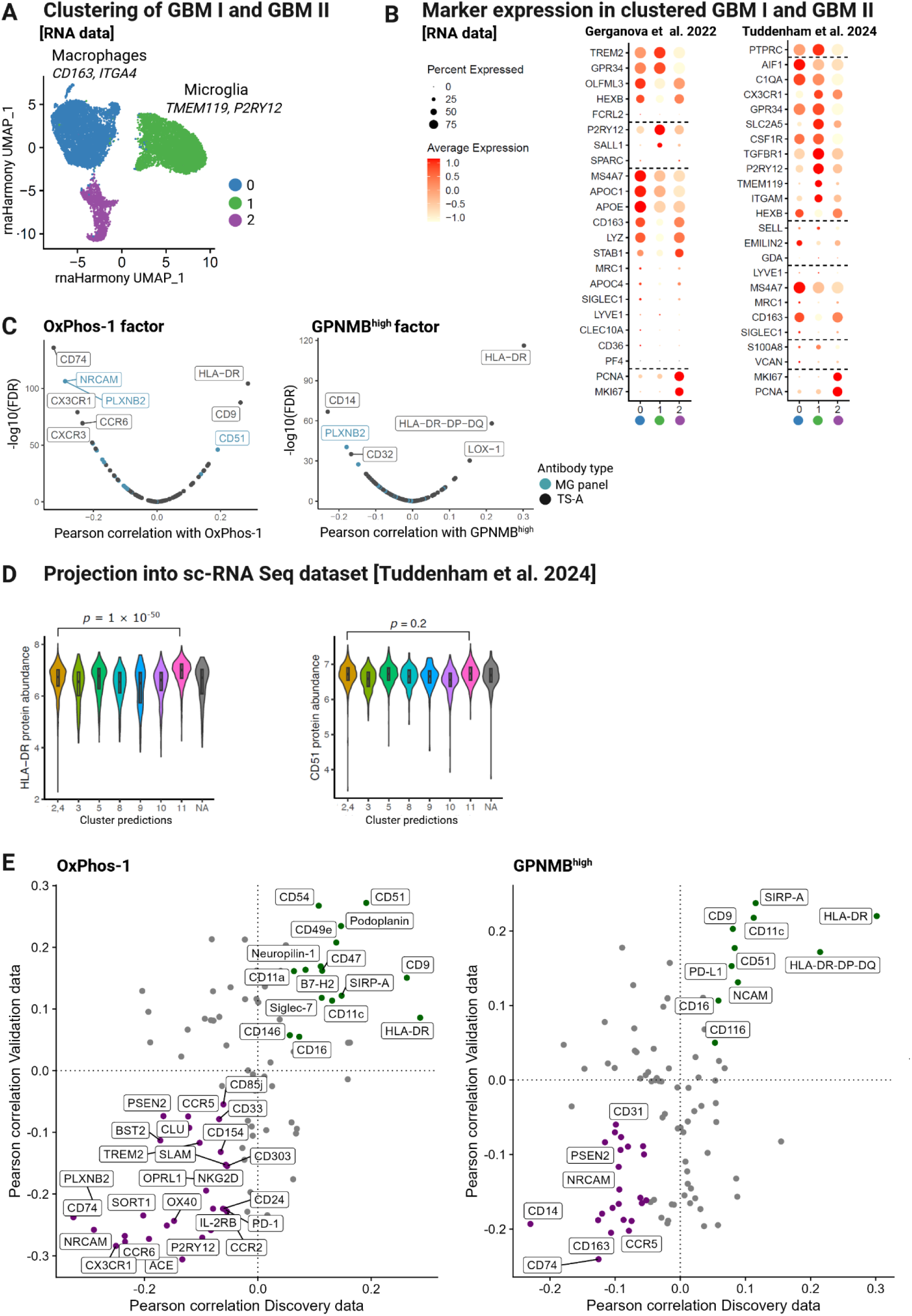
A. Hierarchical clustering of joint cells from GBM I and GBM II. **A.** Graph represents hierarchical clustering of joint cells derived from GBM I and GBM II samples. Clustering resulted in three clusters, termed 0 (blue), 1 (green) and 2 (purple). **B. Expression of defined microglia- and macrophage markers across the identified clusters derived from GBM I and GBM II.** Marker expression of microglia- and macrophage markers as previously used by Gerganova et. al. ^33^(left) and Tuddenham et al.^20^(right) across the clusters 0,1,2 derived from joint GBM I and GBM II samples. Plots show markers derived from both datasets and their expression across the different cluster 0,1,2. Average expression is indicated by color scheme with high expression in red and low expression in light yellow. Cluster 1 emerges as a microglia-specific cluster. **C. Projections of scHPF factors into isolated microglia from GBM I and II.** Using Cluster 1 identified as microglia, previously identified scHPF factors^30^ were projected into the microglia cluster. Plots show the Pearson correlation for each of the factors (OxPhos-1 - left, GBNMP^high^-right) for the CITE-Seq antibodies (microglia (MG) panel – turquoise; TS-A Human Universal Cocktail 1.0 - black) with each of the assessed factors. Correlation for all antibodies is shown in relation to -log10(FDR) values. Antibodies with lowest and highest correlation are labelled in different, colors depending on their origin - turquoise (microglia-specific panel) or black (TS-A Universal cocktail). **D. Projection into sc-RNA Seq dataset of human microglia (Tuddenham et al. 2024**^20^**) and expression of HLA-DR and CD51 across the sc-RNA Seq microglial clusters.** Violin plots show the protein abundance of selected proteins – HLA-DR (left) and CD51 (right) across the 11 different clusters. Internal box plot indicates the first, second, and third quartiles of protein abundance. For statistical analysis, the Wilcoxon Rank Sum test was performed. **E. Comparison of correlation analysis for factors OxPhos-1 and GPNMB^high^ between the discovery dataset and the validation dataset**. For each factor (OxPhos-1 – left; GPNMB^high^ – right;), the Pearson correlation results (FDR) between the ADTs (antibody-derived tags; Proteins) and the genes within each factor are plotted with the Pearson correlation (FDR) for the discovery data (GBM I-II) on the x-axis and the Pearson correlation (FDR) for the validation data (GBM IV-IVI) on the y-axis. Each dot indicates one protein. Green dots indicate proteins with positive correlation with FDR >0.05 (OxPhos-1; GPNMB^high^), purple dots indicate proteins with negative correlation with FDR >-0.05 (OxPhos-1; GPNMB^high^) including labelling.

We then systematically correlated ADT expression of the 101 antibodies with each of the 23 factors (**Table S4A**). Interestingly, for the OxPhos-1 transcriptional program, we observed the highest level of correlation with the HLA-DR, CD9 and CD51 proteins; for the GPNMB^high^ factor, we observed the highest correlation with HLA-DR protein levels followed by expression of an epitope shared by HLA-DR/-DP/-DQ (**Figure 5C**). When assessing the correlation between HLA-DR or CD51 with the OxPhos-1-like factor and the GPNMB^high^-like factor (**Figure S5F**), HLA-DR surface expression displayed a similar level of correlation with both the OxPhos-1- and the GPNMB^high^-like factors (OxPhos-1 r= 0.29; GPNMB^high^ r= 0.30), suggesting that HLA-DR could contribute as a general marker for both subtypes. CD51 showed a stronger correlation with the OxPhos-1-like factor (r= 0.19 vs. GPNMB^high^- r=0.08), suggesting that CD51 is more strongly linked to the OxPhos-1-like state.

To complement these analyses with a cell cluster-based approach, we projected our GBM-derived microglia into a second, cluster-based model of microglial subtypes derived from an earlier version of our scRNA-seq data^10^ of purified human microglia, focusing on 11 clusters that partitioned non-proliferating microglia (**Figure 5D**). Among the GBM I/II microglia, we detected cells corresponding to the signature of each of the defined subtypes (**Figure S5D**). Most cells corresponded to clusters 2 and 4, as well as cluster 11, a cluster that we previously defined as enriched for the DAM signature and high in GPNMB expression^10^. As we have previously done, we merged related clusters together for comparative analyses, grouping clusters 2 and 4 together (homeostatic microglia).^20^ We then analyzed the protein expression of HLA-DR and CD51, and we find that HLA-DR is significantly increased in the GPNMB^high^-expressing cluster 11 microglia relative to the homeostatic microglia of cluster 2,4; however, CD51 showed no significant difference between these two clusters, illustrating how individual protein markers could be helpful to distinguish transcriptomically-defined microglial subtypes.

In order to replicate the results of our association analyses, we generated a second independent dataset of mCITEseq data from GBM samples. Namely, we used the same experimental pipeline and isolated human microglia/macrophages from four additional glioblastoma patient samples (GBM IV-GBM VII; **Figure S5L-N**) and projected the cells from each sample into the scHPF model and its 23 factors^30^: GBM IV = 16853 cells; GBM V = 14183 cells; GBM VI = 11703 cells; and GBM VII = 14286 cells (**Figure S5L**). As previously, we identified a microglia-specific cluster and other cell clusters consisting of macrophages, proliferating myeloid cells and T cells (**Figure S5O-Q**^33,20^). We then assessed the correlation between ADT protein expression levels and the 23 scHPF factors in the microglial cluster only (**Figure S5S; Table S4B**). **Figure 5E** depicts the correlation coefficients for both of our datasets - the discovery (GBM I-II) vs. the validation (GBM IV-VII) datasets for the OxPhos-1- and GPNMB^high^ factors, thereby highlighting the proteins with the most positive and most negative correlation across both datasets for each of the factors (FDR threshold = 0.05).

Overall, the associations of individual proteins with the expression level of these two factors is fairly consistent across the two datasets. For example, CD51, CD54 and PDPN (Podoplanin) showed the highest positive correlation for OxPhos-1 in both datasets, and this transcriptional program is significantly less expressed in cells decorated with CX3CR1, P2RY12 and CCR6. This is consistent with the concept that the DAM1 signature (captured by the OxPhos-1 factor) identifies a subset of microglia that are distinct from homeostatic microglia, which can be defined, in part, by high P2RY12 expression (**Figure 5E**). For the GPNMB^high^ factor, HLA-DR, SIRPA, CD9, and an HLA-DR-DP-DQ epitope are among the most highly correlated markers, while CD74, CD14 and CD163 are depleted in microglia with high expression of the GPNMB^high^ factor.

### *In silico* sorting identifies CD49D, HLA-DR and CD32 as protein markers for human microglia enriched for the GPNMB^high^ factor

As a next step, we aimed to prioritize a small number of marker proteins that could be used to sort human microglia enriched for each of our 23 factors^30^. Identifying such markers would be helpful to develop sorting strategies using a platform such as flow-activated cell sorting (FACS) which typically utilizes a small number of protein markers to isolate target cells from a mixture of cells. We therefore undertook an effort to develop a marker panel *in silico* and operationalized this by deploying a decision tree algorithm that defines gating thresholds based on protein abundance to optimize the isolation of cells with high expression of a given scHPF factor using only 3 protein markers. We completed this selection process for each of the 23 factors and illustrate it using the solution for sorting cells expressing a high level of the GPNMB^high^ factor (**Figure 6A**). The other factors are shown in **Figure S5V** and **Table S4C**. We first deployed the strategy in the discovery dataset (GBM I-II), yielding, for example, a sequence involving gating by CD49D, HLA-DR and finally CD32 to isolate cells with a high level of the GPNMB^high^ factor (**Figure 6A)**. Specifically, the target cells are CD49D^low^, HLA- DR^high^ and CD32^low^; this gating strategy takes our set of mixed microglia found in the discovery set (which has 18.2% of GPNMB^high^ microglia) and yields a subset in which 53.6% of the microglia are GPNMB^high^ (an enrichmenrt of +35.4%; **Figure 6B**). When we deployed this GPNMB^high^ gating strategy derived from the discovery data into the validation data, we found that the proportion of GPNMB^high^ microglia is also enriched, in this case from 15.9% to 31.4% (+15.41%; **Figure 6C**).

**Figure 6.**
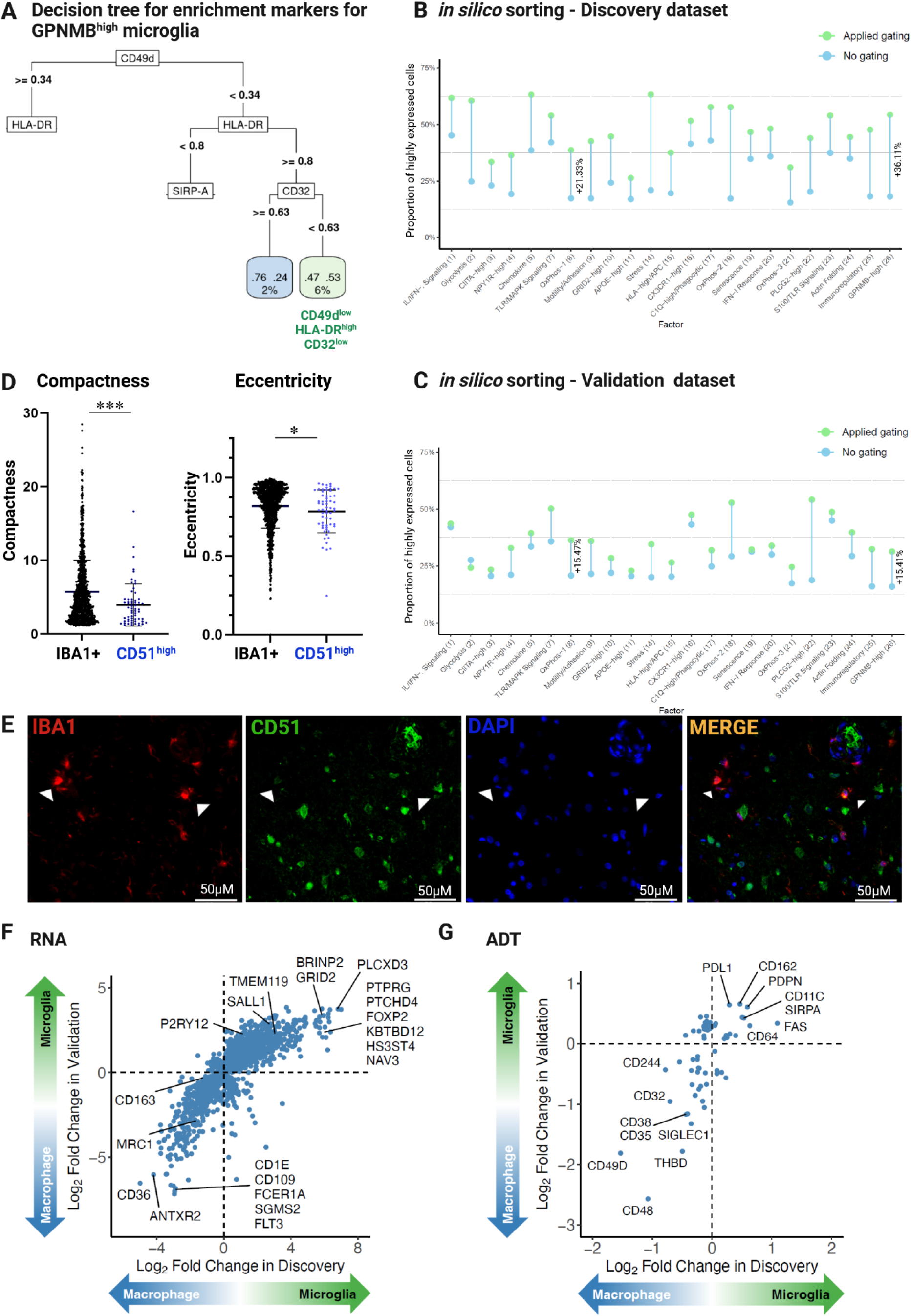
CD51+ microglia, in silico sorting strategy and glioma-associated microglia/macrophage marker identification. A. Decision Tree indicating the potential of selected marker proteins to enrich for. GPNMB^high^ microglia. Square boxes indicate marker proteins (CD49d, HLA-DR, CD32) selected by machine learning to maximize the enrichment of GPNMB^high^ microglia. Branches indicate decisions based on computed expression levels (arbitrary) for each marker. For lower expression levels branches turn left, for higher expression levels branches turn right. Round boxes indicate number of enriched cells in % starting from total cell input (100%) on the top. Green colored box indicates highest potential of marker combination to enrich for GPNMB^high^ microglia – resulting in 6% of total microglia when sorted based on CD49dl^ow^, HLA-DR^high^ and CD32^low^ expression. B-C. *In silico* sorting using the top 3 defined markers for each of the 26 defined scHPF factors^30^ using the discovery (**B**) or validation (**C**) dataset GBM IV-VII. X-axis depicts each of the assessed 26 scHPF factors/expression programs, y-axis indicates proportion of highly expressed cells for the assessed factors in percent (%). Blue dots indicate proportion of highly expressing cells for each of the scHPF factors without prior gating, while green dots indicate percentage of highly expressing cells for each of the factors following gating based on the previously defined top 3 protein markers. Numbers between no gating and applied gating datapoints indicate enrichment (%) for any given scHPF factor in the proportion of highly expressed cells. **D**. **Left**: Dot plot depicting measured compactness of all IBA1+ vs. IBA1+CD51+ cells. Each dot represents a single cell. For statistical analysis, Mann-Whitney test was performed. *p ≤ 0.05; **p ≤ 0.01; ***p ≤ 0.001. **Right:**.Dot plot depicting measured eccentricity of all IBA1+ vs. IBA1+ CD51+ cells. Each dot represents a single cell. For statistical analysis, Mann-Whitney test was performed. *p ≤ 0.05; **p ≤ 0.01; ***p ≤ 0.001. **E.** IHC (FFPE sections) of DLPFC region (BA9) from human AD brain tissue were co-stained with DAPI (mountant, 347 channel), CD51 (1:100, 488 channel, Invitrogen Cat#: PA5-80745), and IBA1 (1:100, 647 channel, Wako cat#01919741). Heat-mediated antigen retrieval was performed with Citrate buffer, pH 6. Images were taken at 40X. Individual panels show staining for microglia using IBA1 (red; upper left), CD51 (green; upper right), nuclei using DAPI (blue; lower left) and merged images (lower right). **F. Differentially expressed genes between tumor-associated microglia and tumor-associated macrophages isolated from human Glioblastoma multiforme (GBM).** x-axis represents log fold change of RNA expression in the discovery data; y-axis represents log fold change of RNA expression in the validation data; Positive log FC means upregulation in microglia (green arrow), whereas negative log FC represents upregulation in macrophages (blue arrow). Only genes that passed FDR < 0.05 both in discovery and in validation datasets are shown. **G. Differentially expressed proteins (ADTs) between tumor-associated microglia and tumor-associated macrophages isolated from human Glioblastoma multiforme (GBM).** x-axis represents log fold change of ADT/protein expression in the discovery data; y-axis represents log fold change of ADT/protein expression in the validation data; Positive log FC means upregulation in microglia (green arrow), whereas negative log FC represents upregulation in macrophages (blue arrow). Only proteins that passed FDR < 0.05 both in discovery and in validation datasets are shown.

Results for the other factors are shown in **Figure S5W-X, Table S4C**. One of the strongest enrichments is seen with PLCG2^high^ microglia: two markers - CX3CR1 and LAMP1 – yield an increase of 23.7% in the proportion of PLCG2^high^ cells in the discovery dataset and an increase of 35.3% in the validation dataset (**Figure S5W-X**). The degree of enrichment varies across factors, and all but one of the decision trees (for the Senescence factor) derived from the discovery set returns an enriched set of cells in the replication set (**Figure S5W-X**). This suggests that our marker selection strategy is robust. These marker combinations will need to be refined, but they outline an approach by which sorting of microglia with certain transcriptional features could be attained, enabling the functional characterization of these selected subsets. The last question, then, that remains to be answered is whether these markers identify microglial subsets in human tissue, prior to cell purification.

As our analyses identified CD51 as potential new markers for human microglia with high expression of the OxPhos-1 factor, we wanted to assess whether we could use this new microglial marker (one of our 17 new antibodies) to confirm the existence of CD51+ microglia in the human brain. This antibody was compatible with immunofluorescence staining of human brain tissue, so we used it to stain a sample of frontal cortex tissue derived from an individual diagnosed with AD (**Supplementary Table S1; Figure 6D-E**). Using an automated image segmentation pipeline based on CellProfiler^34^ optimized for microglia^35^, we segmented 1356 IBA+ cells and assessed their CD51 expression. Defining CD51+ cells as those cells with a CD51 mean fluorescence intensity ≥ 2 standard deviations from the median of CD51 expression (**Figure S5Y**), we found that 4.5% of all IBA1+ cells in this sample are CD51+ (**Figure S5Z**). To begin to explore the function of these cells, we assessed the morphological characteristics of CD51+ microglia: both the eccentricity (p= 0.0178) and compactness (p= 0.0013) of CD51+ microglia were significantly reduced in CD51+ relative to CD51-IBA1+ cells (**Figure 6D**), suggesting that CD51+ cells have a more circular shape than IBA1+CD51-cells.

### Differentiating microglia from infiltrating macrophages

The search for markers that distinguish glioma-associated macrophages/monocytes from glioma-associated microglia based on RNA and protein levels is contemporary and crucial for understanding the function of each population along with the potential for targeting each population separately; however, to date, no commonly accepted set of markers for each population has been defined by the field^36,37^.

Our discovery and replication mCITEseq datasets generated from a total of six human glioblastoma tissue samples (discovery: GBM I-II; validation: GBM IV-VII) provide an opportunity to identify RNA transcripts and proteins that are differentially expressed between these two glioma-associated populations. Namely, we prioritize the genes and proteins that were significantly different (FDR < 0.05) in both the Discovery and Replication datasets when comparing glioma-associated microglia to infiltrating macrophages: RNA in **Figure 6F** and ADT/protein in **Figure 6G** (see full results in **Table S5**). *TMEM119* RNA levels, as expected, are higher in microglia (**Figure 6F**), and several proteins are more highly expressed, such as PDL1, CD162, PDPN, SIRPA and FAS. Notably, *SALL1* (Spalt-Like Transcription Factor 1) RNA is more highly expressed in microglia; it is a gene associated with microglial activation that has been reported to suppress glioma cell proliferation and migration^38,39^. Similarly, signaling via CD162 has been shown to play a role in the regulation of glioma-associated microglia towards an immunosuppressive state ^40,41^. Interestingly, the genes and proteins more highly expressed in macrophages have a larger effect size, including CD48, CD49D and THBD at the protein level (**Figure 6G**). CD36, one of the significant transcripts (**Figure 6F**), has recently been described as a marker for tumor-associated macrophages promoting cancer progression^42^. Moreover, CD49D has previously been described as a macrophage marker in GBM tumors^36,43^, and a THBD+ macrophage subpopulations has been closely associated with hypoxia in glioma^44^. Thus, macrophage appear to have more distinguishing markers than microglia, including CD48 which has not been reported previously.

## Discussion

In the present study, we addressed the important challenge of translating insights into human microglial heterogeneity that have emerged from single cell transcriptomic studies to functional studies of this key resident cell type of the central nervous system. The dynamic and plastic nature of microglia makes them difficult to model as measuring function from a pool of microglia or microglia-like cells is vulnerable to the rapidly shifting frequencies of microglial states found in the pool: functional changes attributed to a perturbation deployed in an *in vitro* system could simply be due to changes in the composition of the pool, and lack of reproducibility of functional observations across or within laboratories may simply be due to compositional differences in the cellular substrate of the experiment. Transcriptomics has been critical to uncover microglial heterogeneity, but RNA-based measures are not optimal to help refine functional studies, particularly as they are an indirect measure of many cellular functions. This is illustrated by the high variability in the level of correlation between RNA and protein measures (**Figure 3E**). Proteins offer a more proximal target to measure cellular function, are a frequent therapeutic target, and have many tools that enable monitoring of individual cells in a cost-effective manner. Further, as seen with other immune cells ^16,45^, proteins offer better resolution of the architecture of microglial cellular populations (**Figure 3D**) and pave the way for sorting subsets of microglia with particular features to understand the relation of molecular features to function. Overall, proteomic characterization is not a panacea, but it represents an important step in the functional characterization and manipulation of microglia and other CNS myeloid cells.

We contribute to the efforts of the human microglial community by sharing a few different **Resources**: (1) additional proteomes from purified, living human microglia (**Figure 1B**), (2) additional antibodies targeting cell surface protein epitopes on human microglia that can be added to a commercially available antibody cocktail to form the mCITEseq panel (**Figure 2**), (3) prioritized proteins that can be explored further to resolve human CNS macrophage from microglia (**Figures 4-6**), and (4) a structured approach to sort human microglia enriched for different transcriptomic features (**Figure 6C-E**). An important ancillary observation is that, while current model systems (iMGs, HMC3 microglia) have distinct transcriptomic and proteomic profiles compared to primary human microglia, a small subset of iMGs and even of the microglia-like HMC3 cell line attain a proteomic profile that places them in the middle of the primary human microglia (**Figure 3D**). This suggests that these model systems have the capacity to improve and offer a more accurate representation of human microglia. With the antibodies that we have characterized in hand, we now have important new tools that can be used for this model system optimization.

While identifying markers for individual “microglial subtypes” is intuitively the clear question that the community is pursuing, defining such subtypes is not straightforward since, from a transcriptomic point of view, the boundaries between groups of microglia with similar profiles are fuzzy, with individual cells being distributed along different gradients^11,20,46^. Our proteomic data to date suggest that proteomic measures will probably not help to identify clear microglial subtypes. We therefore focused our analyses on a panel of 23 transcriptional programs (that we refer to as “factors”) that we have previously developed in a large set of 441,088 transcriptomes from individual human microglia^30^; these factors are more biologically interpretable than microglial clusters and several are associated with Alzheimer’s disease-related traits^30^. One of these is factor 26 (the GPNMB^high^ factor) which is enriched for the DAM2 signature and is associated with AD and AD-related traits; so, we used this factor to illustrate our results, which are available for all 23 factors (**Figure S5**). As a contrast, we used the related DAM1 signature, which is thought to precede the DAM2 state^32^ and is not associated with AD-related traits^30^; here, the OxPhos-1 factor captures the DAM1 signature. Noteworthy, one of the new antibodies that we have evaluated here, the one recognizing a CD51 epitope, appears to be one of the epitopes that is differentially abundant between the OxPhos-1 and GPNMB^high^ factors that are closely related and share, for example, HLA-DR as a protein marker.

Given that microglia do not form clear, discrete subtypes, it is not surprising that purifying microglia enriched for a given factor/transcriptional program is challenging. Nonetheless, our *in silico* sorting approach returned combinations of three proteins that can be used to gate cells that express a high level of each of our 23 factors; combinations that are robust since the purification process is replicable for all but one factor (**Figure 6D-E**). Much work is now needed to further validate this approach using alternative technologies such as flow cytometry. Further, these marker combinations could be used *in situ*, in human brain tissue, and *in vitro* where they could be used to monitor the proportion of cells that have attained a given state. We provide an example of that here as we profiled iMGs from three different laboratories (**Figure 3**).

While our work provides a CITE-Seq cocktail specifically engineered to assess human microglia, two studies using commercially available CITE-Seq technology on human brain have emerged over the past two years^47,48^. CITE-Seq and mass cytometry has been used to characterize subpopulations of CNS-associated macrophages (CAMs) to support the broader goal of generating a comprehensive molecular census of the immune compartment at the human CNS interfaces^48^. This study also profiled Glioblastoma Multiforme (n=21 tumors) using the commercial Universal cocktail of TS-B instead of TS-A technology^48^. With regards to their CITE-Seq analysis of GBM-isolated microglia, the authors report a downregulation of CXCR3 along with an upregulation of CD112, CD58, CD33 and NRP1 in tumor-associated microglia^48^. When comparing these data to our results, we also observed a downregulation of CX3CR1 along with an upregulation and of CD58 (LFA-3) in microglia, while CD112 (Nectin-2) and Neuropilin-1 (NRP1) showed an upregulation in the cluster (0) which we identified as glioma-associated macrophages in our study (**Figure S6C**). Conversly, we do not see replication of our most extreme surface markers for tumor-infiltrating macrophages (CD48 and CD49D) in the prior study^48^. The limited replicability of results across the two studies is not surprising given their small sample sizes and the heterogeneity found among the tumor samples. Larger, dedicated studies are needed and will benefit from further expansion of the mCITEseq cocktail. Interestingly, CD48 is more highly expressed in GBM with a high proportion of cell death, which may be related to poor prognosis with immune checkpoint blocker therapy^49^, and greater CD49D RNA expression in GBM is associated with shorter progression-free survival^50^.

As with any study, there were limitations to the work presented here. Antibody titrations would have ideally been performed on freshly isolated human microglia or iPSC-derived microglia; however, this was not practical given the challenges of each of those sample types and the variability amongst brain samples and among iMG preparations. Further, microglial specificity is operationalized here by significantly higher levels of expression in microglia as compared to PBMCs/peripheral monocytes rather than fully exclusive expression on human microglia. Markers that we identified to be highly enriched for human microglia include OPRL1, P2RY12, SORT1, NRCAM, ITGB5 and CD51. Resolving microglia from CNS macrophage is a more difficult challenge that needs to be addressed next; while we performed these analyses in our brain tumor samples and we have prioritized certain proteins, our study was not designed to explore this question. It will need much larger samples and a broader sampling of myeloid cells found in the CNS. In addition, we would like to highlight that our effort was limited by the availability of antibodies targeting our prioritized surface proteins; much of this is due to limited antibody identification, but there were also some antibodies from a subset of providers that were not considered given their cost or being encumbered by the vendor in terms of permissible uses. The next iteration of this project will therefore aim to take more of the identified candidates into account by not only including the possibility to convert identified antibody clones into recombinant antibodies suited for CITE-Seq but also extending our search to the discovery of novel antibodies targeting our identified microglia-specific candidates. We further aim to perform an extended validation in a bigger sample set of freshly isolated human microglia along with the development of a spectral flow cytometry panel based on our identified microglia-specific protein markers.

In conclusion, we introduce mCITE-Seq as a new proteogenomic tool for the neuroscience community, and we illustrate its utility in exploring the relation of protein markers to the well-described transcriptomics-based heterogeneity of human microglia. In particular, it is clear that proteomics is unlikely to resolve discrete microglia subtypes. Our factor-based approach better captures the transcriptional gradients that underlie this heterogeneity, and we see that selected groups of surface proteins can be used to purify cells with high expression of individual factors, such as the GPNMB^high^ factor that is associated with AD. Thus, we facilitate the transition to studying the function linked to these factors in enriched subsets of microglia. Given that we leveraged GBM samples as a source of myeloid cells, we also described limited protein differences between microglia and macrophage in this context, which suggests that the two cell types may be converging to similar states when present in a shared environment. Overall, these data have yielded a number of important insights – such as the facts that a minority of microglia-like cells from model systems adopt a fate that is proteomically similar to primary microglia – that need to be elaborated with a richer antibody panel and larger number of samples *ex vivo* and *in vitro* to yield a refined understanding of human microglia that is supported by model systems that are optimized with a new proteomic reference.

## Methods

### Source of central nervous system specimens

Details of the acquisition of autopsy samples from the Rush Alzheimer’s Disease Center (RADC)^51,52^ in Chicago, IL (Dr. Bennett) and Columbia University Medical Center/New York Brain Bank in New York, NY (Drs. Vonsattel and Teich)^53^, as well as surgically resected brain specimens from Brigham and Women’s Hospital in Boston, MA (Drs. Sarkis, Cosgrove, Helgager, Golden, and Pennell) were detailed in our prior publication^6^. In addition, samples were obtained from donation programs at Rocky Mountain Multiple Sclerosis Center, Denver, CO (Dr. John Corboy) and Alzheimer’s Disease Research Center/Precision Neuropathology Core of University of Washington School of Medicine (Dr. C. Dirk Keene). All brain specimens were obtained through informed consent and/or brain donation program at the respective organizations. All procedures and research protocols were approved by the corresponding ethical committees of our collaborator’s institutions as well as the Institutional Review Board (IRB) of Columbia University Medical Center (protocol AAAR4962). For a detailed description of the brain regions sampled, clinical diagnosis, age and sex of the donors, as well experimental use (proteomics *versus* CITE-Seq validation) see **Table S1**.

### Shipping of brain specimens

After weighing, the tissue was placed in ice-cold transportation medium (Hibernate-A medium (Gibco, A1247501) containing 1% B27 serum-free supplement (Gibco, Cat#: 17504044) and 1% GlutaMax (Gibco, Cat#: 35050061)) and shipped overnight at 4°C with priority shipping.

### Human microglia isolation for shotgun proteomics – Columbia University

Upon arrival of the autopsy and surgical resection brain samples, the cerebral cortex and the underlying white matter were dissected on ice. The dissected tissue was placed in ice-cold HBSS (Lonza, Cat#: 10-508F) and weighed. The tissue was then homogenized in a 15 ml glass homogenizer with a maximum amount of 0.5g tissue at the time. The resulting homogenate was subsequently filtered through a 70 μm filter and centrifuged at 300 g for 10 min. The pellet was resuspended in 2 ml staining buffer (PBS (Lonza, Cat#: 17-516F) containing 1% FBS) per 0.5 g of initial tissue and incubated with anti-myelin magnetic beads (Miltenyi, Cat#: 130-096-733) for 15 min according to the manufacturer’s instructions. The homogenate was than washed once with staining buffer and myelin was depleted using Miltenyi large separation (LS) columns. The flow through was collected and after centrifugation the pellet was then incubated with anti-CD11b AlexaFluor488 (BioLegend, Cat#: 301318, clone ICRF44) and anti-CD45 AlexaFluor647 (BioLegend, Cat#: 304018, clone HI30) antibodies as well as 7AAD (BD Pharmingen, 559925) for 20 min on ice. Subsequently the cell suspension was washed twice with staining buffer, filtered through a 70 μm filter and the CD11b+/CD45+/7AAD− cells were sorted on a BD Influx sorter. 5000 CD11b+/CD45+/7AAD-cells were sorted in A1 well of a 96 well plate containing 100 μl of PBS. Following FACS the cells were spun down after which all but 5 μl of PBS was removed from A1 well and the plate was than snap frozen on dry ice and stored at −80 °C until further processing.

### Shotgun proteomic analysis of isolated human microglia - PNNL

Sorted cell pellets were lysed and processed as previously described^54^. Briefly, samples were incubated with n-dodecyl β-d-maltoside-based lysis buffer at 70°C for 60 min followed by room temperature incubation for 10 min. Trypsin digestion was performed in two stages; initial digestion was performed with 25 ng of trypsin at 25°C for 2 hr, followed by addition of 50 ng trypsin and overnight incubation at 25°C. After quenching with acid, samples were desalted with Empore C18 extraction disks. Peptides were separated using a 120 min gradient on a 50 µm × 50 cm C18 (Phenomenex Jupiter) analytical column. The liquid chromatography system was coupled to a Q-Exactive Plus mass spectrometer (Thermo Fisher Scientific). Peptide identification and quantification was performed using the FragPipe (v21.1) computational platform using mostly default settings with MSFragger^55,56^ (v4.0), IonQuant^57^ (v1.10.12), and Philosopher^58^ (v5.1.0). Key parameters are as follows: fully tryptic search with proline rule, methionine oxidation and N-terminal acetylation as variable modifications, iodoacetamide alkylation of cysteine as a fixed modification, no matching between runs, false discovery rate (FDR) limited to 1% at protein level. Label-free quantitation (LFQ) was used to measure peptide abundance and protein inference was performed by ProteinProphet^59^. Protein intensity was converted to protein copy number using proteomic ruler^23^ by normalizing all protein intensities to the sum of all detected histone intensities.

### Computational analysis of proteomics results of freshly isolated human microglia

Protein abundance data were obtained and read into RStudio running R v4.3.1. Each protein’s UniProt ID was mapped to Ensembl Gene IDs for consistent gene representation using the biomaRt package v2.58.2.

The dataset was filtered to only include columns relevant to iBAQ data for each protein across multiple samples. To normalize the data, iBAQ values were log-transformed (log₂), with a constant of 1 added to each value to avoid undefined values resulting from zeros. Proteins with zero values across all samples were excluded.

To identify proteins with the highest variability across samples, standard deviation (SD) was calculated for each protein. The top 500 proteins with highest SD were selected for further analysis. A heatmap of the standardized protein abundances for these proteins was generated using the pheatmap package with row scaling (mean-centered and SD-scaled). Additionally, principal component analysis (PCA) was performed on this subset to visualize sample clustering patterns, with particular focus on disease status differences.

### Correlation analysis between sc-RNA Seq datasets with matching proteomics datasets from the same samples of freshly isolated human microglia

#### Data Processing and Normalization

The Seurat object of scRNA-seq data from microglial cells was processed using the Seurat package in Rstudio v2023.06.1+524. Pseudobulk aggregation of RNA-seq data was performed using the PseudobulkExpression function, grouping by the library names. Normalization was conducted using the LogNormalize method with a scaling factor of 10,000. This was done to facilitate correlation calculations against proteomics data downstream.

#### Proteomics Data Preprocessing

Proteomics data was loaded as a dataframe from an existing spreadsheet. Columns in this dataset were filtered to only include data from libraries in common with the scRNA-seq data. Row names were retained from the original data, and column names were updated based on mappings between individual IDs and scRNA-seq library names. In cases where duplicate column names existed, values were averaged across the duplicates using the rowMeans function.

Both dataframes were filtered to retain only the intersecting rows (proteins) and columns (libraries), ensuring consistency across both datasets.

#### Pearson Correlation Analysis

Pearson correlations between the proteomics and scRNA-seq datasets were computed using the cor function. For each common protein, the Pearson correlation coefficient was calculated between corresponding rows in each dataframe.

#### Visualization and Statistical Analysis

A histogram of the Pearson correlation coefficients with 20 bins was created. The average Pearson correlation coefficient across the 2,246 common proteins was calculated, yielding r=0.04. A one-sample t-test was performed to test the null hypothesis that the mean correlation was zero, resulting in a significant p-value (p<1.8×10^-16).

#### Shotgun proteomic analysis of HMC3 microglia

HMC3 cells were cultured and processed as previously described^60^. Briefly, HMC3 cells were lysed by incubating in a 8M urea-based lysis buffer. Disulfides were reduced by 5mM dithiothreitol for 1 hr at 37°C. Cysteine residues were alkylated with 10 mM iodoacetamide for 45 min at 25°C. After dilution to 2M urea, proteins were digested with LysC for 2 hr at 25°C followed by overnight trypsin digestion at 25°C. Digestion was quenched by acidification and 200-mg tC18 SepPak cartridges were used for desalting. Peptides were labeled with TMT-16 at 25°C for at least 1 hr before quenching. Samples were desalted using 200-mg tC18 SepPak cartriges prior to LC-MS/MS analysis. Peptides were separated using a 120 min gradient on an in-house prepared 75 µm x 25 cm C18 (Waters BEH) analytical column. The liquid chromatography system was coupled to a Q-Exactive Plus mass spectrometer (Thermo Fisher Scientific). Peptide identification and quantitation was performed using the FragPipe (v21.1) computational platform using mostly default TMT-16 settings with MSFragger^55,56^ (v4.0), IonQuant^57^ (v1.10.12), and Philosopher^58^ (v5.1.0). Key parameters are as follows: fully tryptic search with proline rule, methionine oxidation and N-terminal acetylation as variable modifications, iodoacetamide alkylation of cysteine as a fixed modification, false discovery rate (FDR) limited to 1% at protein level. Peptide abundance was measured using TMT-16 reporter ion intensities and protein inference was performed by ProteinProphet^59^. Protein intensity was converted to protein copy number using proteomic ruler^23^ by normalizing all protein intensities to the sum of all detected histone intensities.

### Comparative analysis of different human microglia preparations and model systems

As input datasets, we selected several published as well as unpublished datasets generated within the realm of this study, all summarized in **Table 4**: for HMC3 microglia, we selected published data from serum-free and serum-cultured (10% FCS) HMC3 cells^21^ and HMC3 microglia proteomics generated for this study. For iMG, we chose proteomic datasets generated by Llyod et al.^14^ and Chou et al.^17^ and a Columbia University internal iMG proteomics dataset generated for this study. For freshly isolated human microglia proteomics datasets, we used the dataset generated by Lloyd et al.^14^ as well as our generated dataset. As a control cell type, we included peripheral blood mononuclear cells (PBMCs) using three proteomic datasets derived from human PBMCs ^22,61,62^. Within the initial QC step of the analysis, Zeng et al.^61^ was excluded due to poor proteome coverage and no availability of histone data, while Ravenhill et al.^62^ was excluded as this dataset consisted only of cell surfaceome proteomics with also no histone expression data. Peptide identification for both data-dependent acquisition label-free quantitation (DDA-LFQ) and TMT-based analysis was performed using the FragPipe (v21.1) computational platform using mostly default settings with MSFragger^55,56^ (v4.0), IonQuant^59^ (v1.10.12), and Philosopher^58^ (v5.1.0). Key parameters are as follows: fully tryptic search with proline rule, methionine oxidation and N-terminal acetylation as variable modifications, iodoacetamide alkylation of cysteine as a fixed modification, no matching between runs, false discovery rate (FDR) limited to 1% at protein level. Datasets collected using data independent acquisition (DIA) were analyzed using DIA-NN^63^ (v1.8.2 beta 39) in library-free mode with default settings. Precursor ions were generated from the Swiss-Prot entries in the UniProt database (downloaded April 26, 2024) with precursor and fragment ion m/z range tailored to the bounds of the dataset. Samples within each dataset were averaged and resulting protein intensities were converted to copy numbers by normalizing to total histone intensity using proteomic ruler^23^. If raw data was unavailable, protein copy numbers provided by the authors were used directly (**Table 4**). Datasets were harmonized by applying a scaling factor, calculated as the median of protein intensity ratios (target dataset/reference dataset), to adjust the median counts to be equal to the median of the dataset with highest number of protein IDs (reference dataset). This corrects for variation in histone content across datasets. Missing values were imputed with the 2% quantile for each dataset. Finally, ratios between mean copy numbers of target model and the non-target datasets were calculated (Fold Difference score; **Figure 2B**) and only UniProt^24^-annotated membrane proteins were retained.

### Identification of common marker proteins across different model systems

Marker proteins which are common to multiple model systems were determined for the following combinations of models: 1. iMG and freshly isolated human microglia, 2. HMC3 and freshly isolated human microglia, 3. iMG, HMC3, and freshly isolated human microglia, 4. iMG and HMC3. Using non-imputed data, proteins were first required to be identified in all samples from the target model systems and absent from all remaining samples. A Fold Difference score was calculated for each system and the Top150 for each model were selected for further refinement. Finally, proteins were required to be most abundant in the target model and Top10/30/100 lists were selected by Fold Difference score.

### Identification of distinct marker proteins between different model systems

The Top150 proteins by Fold Difference score for each model system were selected for refinement to identify model system marker candidates. Proteins were subsequently required to be present in only one Top150 list among all model systems and most abundant in the target model. Top10/30/100 lists were selected from the remaining candidates using highest Fold Difference score.

### Identifying microglia-enriched GWAS risk genes for Alzheimer’s disease

As part of the design of our panel, we were interested in characterizing protein-level expression genes enriched in genetic risk for Alzheimer’s disease. However, given the limited panel size, we wanted to choose genes that were enriched in microglia so as to only include genes with high probability of expression on these cells. To do so, we used a list of GWAS risk genes that we derived from multiple different publications (**Table 1**). To identify microglia-enriched genes in this list, we referenced prior work that identified genes preferentially expressed by microglia in the aged human brain^64^. We thus only chose genes from the list of GWAS-derived AD risk genes that were identified as “microglia-enriched” by p-value and fold-change testing, resulting in 72 genes.

### In silico approach to identify human microglial specific cell surface and membrane proteins

To computationally identify cell surface and membrane proteins specific to microglia from the human microglia proteomics data generated, two distinct approaches employing different databases were employed. Initially, three search strategies using Uniprot^24^ (https://www.uniprot.org/; Uniprot version: Last modified February 2, 2021, analysis conducted in September 2021) were executed. Proteins under the Gene Ontology term “cell surface” [9986], with evidence - any manual assertion, and organism Homo sapiens (Human) were queried, resulting in 733 proteins. Subsequently, proteins under Subcellular location [CC], term Cell surface [SL-0310], with evidence - any manual assertion and organism Homo sapiens (Human) were searched, yielding 52 proteins. Additionally, proteins under Subcellular location [CC], note, term cell surface, with evidence - any manual assertion and organism Homo sapiens (Human) were queried, resulting in 104 proteins. The extracted proteins from these searches were consolidated into a comprehensive list. In the second approach, the surfaceome database (https://wlab.ethz.ch/surfaceome) was utilized^25^. The table_S3_surfaceome, containing details of the full human membraneome, was filtered for the “surface” label, identifying 2063 predicted cell surface proteins. The proteomics data from human microglia were then compared separately against curated Uniprot and surfaceome protein lists, leading to the identification of 57 proteins specific to microglia’s cell membrane or surface from both approaches. An additional six proteins were included from a previously curated list, resulting in a total of 63 candidates for a microglia-specific CITE-Seq panel. Furthermore, an unpublished list of microglia-specific AD genes from a Genome Wide Association Study (GWAS) was screened, adding 28 candidate proteins. We additionally assessed how many of these were ADSP reported risk/protective causal genes, by assessing their presence in the ADSP-reported lists^65,66^. After removing duplicates, a final list of 91 microglia-specific cell surface/membrane proteins was obtained. Expression levels of these candidates were assessed in proteomics datasets of HMC3 microglia (ATCC; Cat #: CRL- 3304), iPSC-derived human microglia, and RNA expression in single-nucleus and single-cell RNA-Seq datasets from human microglia. In the subsequent step, the availability of protein candidates in Total-Seq A format by BioLegend, antibodies in unconjugated or conjugated formats suitable for flow cytometry, supplier information, available clones, and isotypes were considered. Seventeen candidates were excluded due to the unavailability of antibodies, resulting in 74 remaining candidates. Of these, 37 candidates were already included in the TotalSeqA Human Universal cocktail V.1.0 (BioLegend, Cat#: 399907), leaving 37 candidates to be added as novel microglia-specific markers to the CITE-Seq panel. To enable consistent titrations across all antibodies, 32 antibodies from different suppliers were ordered in PE-conjugated format or, if unavailable, PE-conjugated secondary antibodies (see **Table 5**).

### Titration of purchased microglia-specific antibody candidates in HMC3 microglia

To determine the optimal concentration for each of the 32 antibodies to be added to the developing CITE-Seq/TS-A panel, titrations for all antibodies were initially performed on HMC3 microglia. In order to do so, HMC3 microglia (ATCC; Cat #: CRL-3304) were grown until confluency, harvested using Trypsin (Gen Clone; Cat #:25-510F), centrifuged at 1500rpm, 8min, 4°C, counted and 0.5×10^6^ HMC3 microglia/condition were transferred into a FACS tube (Corning; Cat#: 352235) for subsequent staining using different concentrations of each selected antibody. First, 0.5×10^6^ HMC3 were centrifuged at 1500rpm, 8min, 4°C and the pellet was resuspended in 100µl of *live dead aqua* (Thermo Fisher Scientific; Cat#: L34957) and incubated for 30min in the dark at room temperature. Subsequently 100µl PBS were added for washing, cells were centrifuged at 1500rpm, 8min, 4°C, supernatant was removed and the pellet resuspended in 95µl Cell Staining Buffer ( BioLegend, Cat#: 420201). Then, 5µl of Human TruStain FcX™ (BioLegend; Cat#: 422301) were added and the cell suspension was incubated for 5-10 minutes at room temperature. During that step, 100µl of 2x solutions with the recommended concentration, as well as 1:2, 1:4, 1:8, 1:16, 1:32 dilutions of the recommended concentration for each antibody were prepared. Following the incubation with blocking buffer, 100µl of each 2x dilution was then added to achieve 1x of recommended, 1:2, 1:4, 1:8, 1:16, 1:32 concentration of each antibody. An additional vial for incubation with each respective isotype (see Table 6A) matching the recommended concentration of each antibody was prepared, as well as unstained controls, live dead staining controls, as well as in the case of secondary PE-conjugated antibodies, respective control samples were prepared. Samples were incubated on ice for 15-20 minutes in the dark. Subsequently, samples were washed twice by adding 1-2ml of Cell Staining Buffer (BioLegend, Cat#: 420201) to each tube, followed by centrifugation at 350xg for 5 minutes, 4°C. In the case of PE-conjugated primary antibodies, stained cells were resuspended in 500µl Cell Staining Buffer (BioLegend, Cat#: 420201) and stored on ice until readout via flow cytometry. In the case of unconjugated primary antibodies, following the washing, cells were incubated in 100µl Cell Staining Buffer (BioLegend, Cat#: 420201) containing the respective concentration for each PE-secondary antibody as specified in the manufacturer’s instructions (Table 6B).

Cells were incubated on ice for 15-20mins, followed by two washes with at least 1-2ml of Cell Staining Buffer (BioLegend, Cat#: 420201) and centrifugation at 350xg for 5 minutes, 4°C. Stained cells were resuspended in 500µl Cell Staining Buffer (BioLegend, Cat#: 420201) and stored on ice until readout via flow cytometry.

Cells were subsequently recorded via flow cytometry using a 3L Aurora (Cytek Bio). Following unmixing for PE- and *live dead aqua* staining (Thermo Fisher Scientific; Cat#: L34957), cells were gated as follows: SSC-H vs. FSC-H to find the cell population, FSC-H vs. FSC-A to select single cells, FSC-H vs. Comp.-Zombie Aqua-A to select for live cells, Comp-PE-A vs. FSC-H to record PE-stained cells. 10.000 cells/sample and dilution were recorded and saved in the form of fsc. files for downstream analysis. Antibodies that were successfully titrated in HMC3s included: ADAM10 (BioLegend, Cat#: 352703), CD49 (BioLegend, Cat#: 328009), CD51 (BioLegend, Cat#: 327909), HLA-C (BioLegend, Cat#: 373302), ITGB5 (BioLegend, Cat#: 345203), PLXNB2 (Miltenyi, Cat#: 130-095-212), PTPRJ (BioLegend, Cat#: 328708).

In the case of non-detectable staining using any of the selected antibody candidates on HMC3 microglia, antibody titration was performed in PBMCs (*see below*).

### Titration of purchased microglia-specific antibody candidates in Peripheral Blood Mononuclear Cell (PBMCs)

Cryopreserved PBMC/blood collar samples (protocol AAAR4962) samples were thawed as follows: 1ml of prewarmed RPMI 1640 complete media (Thermo Fisher Scientific; Cat#: 61870010) containing 10% fetal calf serum (Fisher Scientific, Cat#: 10-438-026) and 1% Penicillin-/Streptomycin (Gibco, Cat#: 15140-122) was transferred into a 15ml falcon tube. Frozen PBMC/blood collar samples were thawed by gently shaking them in a 37°C water bath for 2mins. 1ml prewarmed RPMI 1640 complete media was transferred to the PBMC/blood collar vial and the solution was subsequently transferred to the 15ml falcon tube (*Sigma Aldrich, Cat#:* CLS430052) containing 1ml RPMI 1640 complete media. 8ml addition RPMI 1640 complete media were then added and the thawed PBMCs/blood collar was centrifuged for 12min, 1200rpm at room temperature. Supernatant was aspirated, cell pellet resuspended in 2ml RPMI 1640 complete media and the cells were counted and cell viability assessed using a Nexcelom cell counter. For titration experiments, 0.5×10^6^ cells/condition were transferred to each FACS tube (Corning; Cat#: 352235) for subsequent staining using different concentrations of each selected antibody.

First, 0.5×10^6^ PBMCs were centrifuged at 1200rpm, 8min, 4°C, washed with PBS (Corning; Cat#: 21-040-CV) once, centrifuged again and the pellet was resuspended in 100µl of *live dead aqua* (Thermo Fisher Scientific; Cat#: L34957) and incubated for 30min in the dark at room temperature. Subsequently 100µl PBS were added for washing, cells were centrifuged at 1500rpm, 8min, 4°C, supernatant was removed and the pellet resuspended in 95µl Cell Staining Buffer ( BioLegend, Cat#: 420201). Then, 5µl of Human TruStain FcX™ (BioLegend; Cat#: 422301) were added and the cell suspension was incubated for 5-10 minutes at room temperature. During that step, 100µl of 2x solutions with the recommended concentration, as well as 1:2, 1:4, 1:8, 1:16, 1:32 dilutions of the recommended concentration for each antibody were prepared. Following the incubation with blocking buffer, 100µl of each 2x dilution was then added to achieve 1x of recommended, 1:2, 1:4, 1:8, 1:16, 1:32 concentration of each antibody. An additional vial for incubation with each respective isotype (see Table 5) matching the recommended concentration of each antibody was prepared, as well as unstained controls, live dead staining controls, as well as in the case of secondary PE-conjugated antibodies, respective control samples were prepared. Samples were incubated on ice for 15-20 minutes in the dark. Subsequently, samples were washed twice by adding 1-2ml of Cell Staining Buffer (BioLegend, Cat#: 420201) to each tube, followed by centrifugation at 350xg for 5 minutes, 4°C. In the case of PE-conjugated primary antibodies, stained cells were resuspended in 500µl Cell Staining Buffer (BioLegend, Cat#: 420201) and stored on ice until readout via flow cytometry. In the case of unconjugated primary antibodies, following the washing, cells were incubated in 100µl Cell Staining Buffer (BioLegend, Cat#: 420201) containing the respective concentration for each PE-secondary antibody as specified in the manufacturer’s instructions (Table 6A). Cells were incubated on ice for 15-20mins, followed by two washed with at least 1-2ml of Cell Staining Buffer (BioLegend, Cat#: 420201) and centrifugation at 350xg for 5 minutes, 4°C. Stained cells were resuspended in 500µl Cell Staining Buffer (BioLegend, Cat#: 420201) and stored on ice until readout via flow cytometry.

Cells were subsequently recorded via flow cytometry using a 3L Aurora (Cytek Bio). Following unmixing for PE- and *live dead aqua* staining (Thermo Fisher Scientific; Cat#: L34957), cells were gated as follows: SSC-H vs. FSC-H to find the cell population, FSC-H vs. FSC-A to select single cells, FSC-H vs. Comp.-Zombie Aqua-A to select for live cells, Comp-PE-A vs. FSC-H to record PE-stained cells. 10.000 cells/sample and dilution were recorded and saved in the form of fsc. files for downstream analysis. Antibodies that were successfully titrated in PBMCs included: BST2 (BioLegend; Cat#: 348405), GLUT1 (R&D; Cat#: MAB1418-SP), SLC7A5 (R&D; Cat#: MAB10390-SP), TREM2 (BioLegend; Cat#: 824801; R&D; Cat#: MAB17291-SP), ABCG1 (Novus Biologicals; Cat#: NB400-132SS), CLU (BioLegend; Cat#: 848701), PILRA (BioLegend; Cat#: MAB64841-SP), TSPO (abcam; Cat#: ab109497), NLGN2 (Novus Biologicals; Cat#: NBP1-00220), ABCB11 (NSJ Bioreagents; Cat#: R31844), NRCAM (BioLegend; Cat#: 821602), PSEN1 (BioLegend; Cat#: 811101), ACE2 (BioLegend; Cat#: 375801), HLA-DPB1 (Novus Biologicals; Cat#: NBP3-07722), OPRL1 (Proteintech; Cat#: 12970-1-AP), PSEN2 (BioLegend; Cat#: 814203), SORT1 (BioLegend; Cat#: 852101), SLC1A4 (LS Bio; Cat#: LS-C179222).

### Analysis of titration and selection of final candidates for customized production of the microglia-specific CITE-Seq panel

Analysis of recorded samples was performed using FlowJo_v10.8.1_CL software. Analysis was conducted according to instructions of the manual FlowJo for Antibody Titrations provided by UWCCC Flow Cytometry Laboratory (http://www.uwhealth.org/flowlab). In short, using SSC-A vs. FSC-A populations were visualized, followed by gating for single cells using SSC-H vs. SSC-A and gating for live cells using FSC-H vs. Comp.-Zombie Aqua-A. Subsequently, the separation index (SI), an index evaluating the staining results was calculated using the following formula:

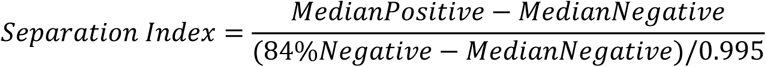

For visualization of the titration data for each antibody including all dilutions, controls and isotype control, populations of live cells were further gated into PE-negative and PE-positive and subsequently concatenated. Within the concatenated file, the axes were then changed to display the antibody channel (PE) on the y axis and the sample ID on the X axis to visualize changes in PE-positive cells across the different dilutions for each of the titrated antibody. Selection of the best suited concentration for each antibody to add to the CITE-Seq panel was then based on assessing the concatenated plots and selecting the concentration at which a plateau in the PE-positive population was reached. Selected concentrations were subsequently discussed with flow cytometry experts and the collaborating team at BioLegend. Following the selection, respective companies were contacted for quotes of 2mg of the selected antibodies as this comprised the amount of antibody needed for custom-conjugation in the TotalSeq-A format by BioLegend. Following negotiations, 17 microglia-specific antibodies were purchased for custom-conjugation (Table 8) with the aim to adding them to the TotalSeq A universal cocktail (BioLegend, Cat#: 399907; 147 targets), resulting in a microglia-marker enriched TS-A custom- cocktail consisting of 164 markers.

The final selection of microglia-specific antibodies contains:

## Validation experiments

### Reconstitution of customized microglia-specific CITE-Seq panel

Reconstitution of the CITE-Seq/Total-Seq A panel was performed according to the manufacturer’s instructions. In short, both lyophilized panels, the custom TotalSeq A (BioLegend, microglia-specific) and the TotalSeq A universal cocktail (BioLegend, Cat#: 399907) were equilibrated to room temperature for 5 minutes. Then, both panels were centrifuged at 10,000 x g for 30 seconds at room temperature and only the lyophilized custom panel (BioLegend, microglia-specific) was reconstituted in 27.5 μL of Cell Staining Buffer (BioLegend, Cat#: 420201), vortexed for 10s and incubated at room temperature for 5 minutes. After vortexing, the resuspended cocktail was centrifuged for at 10,000 x g for 30 seconds at room temperature, the entire volume (27.5 μL) of reconstituted custom cocktail was transferred to the TotalSeq A Universal Cocktail vial, vortexed and incubated at room temperature for 5mins. Following vortexing and centrifugation at 10,000 x g for 30 seconds at room temperature, the entire volume (27.5μL) of reconstituted combined cocktail was transferred to a low protein binding Eppendorf tube (Fisher, Cat#: 022431081) and centrifuged at 14,000 x g for 10 min at 4 °C. After this step, the cocktail was immediately added to the cell preparation following 5min incubation in Human TruStain FcX™ Fc blocking reagent (BioLegend, Cat#: 422301).

### Isolation, hashing and staining for CITE-Seq of freshly isolated human microglia

Microglia were isolated from human postmortem brain tissue samples as previously described by Mattei et al. ^67^. Briefly, postmortem tissue samples were stored on ice while determining their weight. For isolations, max. 2g of brain tissue was used and all steps were carried out at 4°C. pH of all buffers was adjusted to 7.3-7.4 and they were stored at 4°C until usage. A dounce homogenizer (Carl Roth, Cat#: CXE2.1) was placed on ice, filled with 12ml ice-cold Hibernate-A (Thermo Fisher scientific; Cat#: A1247501), while the tissue was minced in a pre-chilled petri-dish with 3-4 mL Hibernate-A (Thermo Fisher scientific; Cat#: A1247501), transferred to the dounce homogenizer on ice. The minced tissue was then dissociated using the loose pestle and subsequently the suspension was passed through a prewet 100µM strainer (Corning, Cat#: CLS352360) into a 50 ml falcon tube (fisher scientific, Cat#: 10203001). The single cell suspension was then centrifuged at 400 rcf, 4°C for 10 min. Meanwhile, isotonic percoll was prepared by adding 1ml 10x DPBS (pH is 7.2-7.4; Thermo Fisher Scientific; Cat#: 14200075) to 9ml pH-adjusted percoll solution (fisher scientific; Cat#: 10607095). Following centrifugation, the supernatant was aspirated and DPBS added to reach a final volume of 12ml. Then, 4ml of isotonic percoll was added, the solution was well mixed before overlaying the solution with 16ml of DPBS (Corning; Cat#:21-040-CV) by by slightly tilting the falcon tube and keeping the pipetboy tip against the wall of the tube to avoid mixing of the layers. The layered sample was then centrifuged at 3000 rcf, 10min, 4°C. Then, the supernatant including the myelin disk in-between the two phases was aspirated, 15ml of cold DBPS (Corning; Cat#:21-040-CV) added for washing, the falcon tube was tilted three times by 180°C and subsequently centrifuged for 10min at 400 rcf, 4°C.

For isolation of microglia from the myelin-depleted cell suspension, cells were first counted using a cell counter (Nexcelom) and subsequently isolated using Magnetic-Activated-Cell-Sorting (MACS) technology, according to the manufacturer’s instructions. In short, the cell suspension was centrifuged at 300g, 4°C for 10mins, then the supernatant was pipetted off completely. Cell pellet was resuspended in 80µl of MACS buffer (PBS pH 7.2, 0.5% BSA (bovine serum albumin) and 2 mM EDTA) per 10^7^ total cells. Per 10^7^ total cells, 20µl of CD11b (Microglia) MicroBeads (Miltenyi; Cat#: 130-093-636) were added, mixed well and incubated for 15mins at 4-8°C. Cells were then washed by adding 1-2ml of buffer per 10^7^ total cells and centrifuged at 300g, 4°C for 10mins. Supernatant was pipetted off completely and up to 10^8^ cells were resuspended in 500µl of MACS buffer (PBS (Corning; Cat#: 21-040-CV), 0.5% BSA (pluriSelect; Cat#: 60-00020-10BSA), 2mM EDTA (Thermo Fisher Scientific; Cat#: AM9260G)). A magnetic column (Miltenyi, LS coumns, Cat#: 130-042-401) was placed into the magnetic field of a suitable MACS separator (Miltenyi, Cat#: 130-042-303) and prepared by rinsing with 3ml of MACS buffer (PBS pH 7.2, 0.5% BSA (bovine serum albumin) and 2 mM EDTA). Subsequently, the cell suspension was applied onto the column, followed by three washing steps consisting of adding 3ml MACS buffer. The column was then removed from the separator, placed on a 15ml falcon tube (Corning, Cat#: CLS430790) for collection, 5ml of MACS buffer () were added and magnetically labelled cells were flushed out by applying the plunger firmly. Isolated CD11b-cells were centrifuged at 300g, 4°C for 10mins and resuspended in 1-2ml DPBS (Corning; Cat#:21-040-CV) for counting using a Nexcelom cell counter.

### Preparation of HMC3 microglia for CITE-Seq

1×10^6^ HMC3 microglia were seeded in a 10cm cell culture dish, cultivated for 3 days and on the third day harvested using Trypsin-EDTA(Gen Clone; Cat #:25-510F). Cells were counted, cell viability was assessed and subsequently 0.7×10^6^ cells were transferred to a 5ml low-protein binding tube (fisher scientific; Cat#: 13-864-407). 1ml cold PBS (Corning; Cat#:21-040-CV) was added, cells filtered through a blue lid filter (FACS tube) back into the 5ml low-protein binding tube (fisher scientific; Cat#: 13-864-407), spun down at 300g, 4°C for 10min with repetition of this washing step. Supernatant was then removed using a pipette tip. Cell pellet was then resuspended in 22.5µL of Cell Staining Buffer (BioLegend, Cat#: 420201), 2.5µL of Human FcBlock (BioLegend, Cat#: 422301) was added and cells were incubated for 10min on ice. For hashing of HMC3 samples together with THP-1 cells and freshly isolated microglia, hashing master mix was prepared as follows: 2µl of TS-A hashtag antibody (0.5 mg/ml, BioLegend, TotalSeq- A0251 anti human Hashtag1, Cat#: B364030; TotalSeq- A0252 anti human Hashtag2, Cat#: B369100; TotalSeq- A0253 anti human Hashtag3, Cat#: B378300; TotalSeq- A0254 anti human Hashtag4, Cat#: B381073) was added to 20µl of Cell Staining Buffer (BioLegend, Cat#: 420201), mixed and passed through a 0.1µm filter at 12000g for 2 min at 4°C. Then, 6.25µl of reconstituted combined CITE-Seq cocktail were added to 18.75µl of hashtag master mix and the combination was added to the cells incubating in FcBlock (BioLegend, Cat#: 422301) and Cell Staining Buffer (BioLegend, Cat#: 420201). After 30min incubation, 4x washing steps consisting of addition of 1ml Cell Staining Buffer (BioLegend, Cat#: 420201) followed by centrifugation at 300g for 10min at 4°C were performed, with the cells before the last wash being filtered through a blue lid filter cap back. Subsequently cells were counted and then processed further for cDNA library preparation.

### Preparation of iPSC-derived microglia for CITE-Seq (iMG I)

iPSC-derived microglia (iMG) were differentiated as previously shown ^18,68^. Briefly, when 70% confluent (day 0), iPSCs were harvested and passaged at a density of 20-40 colonies per well in a 6-well plate coated with 0.1 mg/ml Matrigel (Corning, NY, USA; Cat#: 354277) for the generation of hematopoietic stem cells (HPCs) with the STEMdiff Hematopoietic Kit (StemCell Technologies, Cat#: 05310). On day 1, Medium A was added to the culture and on day 4 it was switched to Medium B, until complete HPC differentiation on day 10-12. Fully differentiated HPCs, detached from the colonies and floating in the medium, were collected for further assays, like microglial differentiation. For iMG differentiation, HPC were cultured in iMG medium consisting of DMEM/F12, with 2X insulin-transferrin-selenite (ITS), 2X B27, 0.5X N2, 1X Glutamax, 1X non-essential amino acids, 400 mM Monothioglycerol, and 5 mg/ml human insulin, freshly supplemented with 100 ng/ml IL-34, 50 ng/ml TGFβ1, and 25 ng/mL M-CSF (PeproTech) for 25-28 days. Cells were subsequently collected by gentle pipetting up and down in PBS and subjected to the CITE-Seq staining protocol as described in *CITE-Seq staining of cell preparations*.

### Preparation of iPSC-derived microglia for CITE-Seq (iMG II)

For iMG differentiation, a two-step differentiation protocol was used^69^. The first step differentiation of 12 days was performed using STEMdiff Hematopoetic Kit (StemCell Technologies, Cat.#: 05310). Herewith, the first three days were supplemented with supplement A, followed by nine days of supplement B exposure for differentiation, with media change every other day. Subsequently, cells were transferred to Poly-L-lysine (PLL) (Sigma, Cat#: P4707-50ML)-coated plates and cultured in conditioned media supplemented with growth factors. Conditioned media was changed every other day and growth factors freshly added at the day of media change. After 8 days of culture in conditioned medium, microglia-like cells (iMG) were collected as floating cells every 2 to 3 days based on their confluency.Subsequently to harvesting, cells were washed 1x with PBS, centrifuged at 300g, 4°C, 5min, filtered through a blue lid filter (FACS tube), resuspended in 100uL PBS+0.04% BSA for counting using counting chambers (Bulldog Bio, Portsmouth, NH) and subsequently subjected to the CITE-Seq staining protocol as described in *CITE-Seq staining of cell preparations*.

### Preparation of iPSC-derived microglia for CITE-Seq (iMG III)

hiPSCs were maintained in StemFlex media (ThermoFisher) on reduced growth factor Cultrex BME (Biotechne, Cat.# 3434-010-02), and routinely split 1-2 times a week with ReLesSR™ (Stem Cell Technologies) without ROCKi as described previously^70^. hiPSCs were differentiated into iMGLs as described previously with minor adaptations^18,71^. In brief, FA000010 (RUCDR/BiologyX) iPSCs were differentiated into hematopoietic precursors cells (HPCs) using the STEMdiff Hematopoietic kit (STEMCELL Technologies) largely by manufacturer’s instructions. In brief, on day -1 iPSCs were detached with ReLeSR and passaged to achieve a density of 1–2 aggregates/cm^2^ of 100-150 cells. Multiple densities were plated in parallel. On day 0, colonies of appropriate density were switched to Medium A from the STEMdiff Hematopoietic Kit to initiate HPC differentiation. On day 3, cells were switched to Medium B with a full media change and fed again with a full media changed on day 5. Cells remained in Medium B for the rest of the HPC differentiation period with Medium B overlay feeds every other day. HPCs were collected 3 independent times by gently removing the floating population with a serological pipette at days 11, 13 and 15 (or days 12, 14 and 16). HPCs were either cryobanked in 45% Medium B, 45% knockout serum replacement (ThermoFisher) and 10% DMSO and stored in liquid nitrogen or directly plated for iMGL induction. HPCs were terminally differentiated at 28,000-35,000 cells/cm^2^ in microglia medium (DMEM/F12, 2X insulin-transferrin-selenite, 2X B27, 0.5X N2, 1X glutamax, 1X non-essential amino acids, 400 mM monothioglycerol, and 5 mg/mL human insulin (ThermoFisher)) freshly supplemented with 100 ng/mL IL-34, 25 ng/mL M-CSF (R&D System) and 50 ng/mL TGFβ1 (STEMCELL Technologies) for every other day until day 24. On day 25, 100 ng/mL CD200 (Bon Opus Biosciences) and 100 ng/mL CX3CL1 (R&D Systems) were added to Microglia medium to mimic a brain-like environment. Differentiated microglia were cultured in 500µl microglia medium (DMEM/F12, 2X insulin-transferrin-selenite, 2X B27, 0.5X N2, 1X glutamax, 1X non-essential amino acids, 400 mM monothioglycerol, and 5 mg/mL human insulin (ThermoFisher). For subsequent CITE-Seq staining, iMGs were harvested in low-protein binding tubes (fisher scientific; Cat#: 13-864-407) using media and PBS (Corning; Cat#:21-040-CV) to wash off iMGs from cell culture wells. Subsequently, cells were washed 1x with PBS, centrifuged at 300g, 4°C, 5min, filtered through a blue lid filter (FACS tube), resuspended in 100uL PBS+0.04% BSA for counting using counting chambers (Bulldog Bio, Portsmouth, NH) and subsequently subjected to the CITE-Seq staining protocol as described in *CITE-Seq staining of cell preparations*.

### Preparation of human PBMCs for CITE-Seq

Peripheral blood samples were drawn from four participants part of the Alzheimer Disease Research Center (ADRC) of Columbia University (IRB-AAAR2387). Briefly, up to 20mL of blood per participant was used to enrich for peripheral blood mononuclear cells (PBMCs) using density gradient centrifugation and series of washes, and followed by a cryopreservation as described in Touil et al.^72^. PBMC samples from individual participants were thawed using RPMI 1640 (Thermo Fisher Scientific; Cat#: 61870010) supplemented with 10% Fetal Bovine Serum (FBS) (Fisher Scientific, Cat#: 10-438-026) and 1% Penicillin-/Streptomycin (Gibco, Cat#: 15140-122) in 37C water bath. Cells washed twice for 12 minutes at a speed of 1200 rpm. Cell numbers from each study participant and viability were measured using the automated Nexcelom cell counter. To proceed with the CITE-Seq staining, we decided to pool PBMCs from four different participants (0.7×10^6^ cells per donor) into a 5ml low-protein binding tube (fisher scientific; Cat#: 13-864-407), to overcome the batch effect. For the washing steps, we centrifuged the samples at 300g, 4°C for 10min. Supernatant was then removed using a pipette tip. Cell pellet was then resuspended in 22.5µL of Cell Staining Buffer (BioLegend, Cat#: 420201), 2.5µL of Human FcBlock (BioLegend, Cat#: 422301) was added and cells were incubated for 10min on ice. Then, 6.25µl of reconstituted combined CITE-Seq cocktail were added to 18.75µl of cell staining buffer (BioLegend, Cat#: 420201) and the combination was added to the cells incubating in FcBlock (BioLegend, Cat#: 422301) and Cell Staining Buffer (BioLegend, Cat#: 420201). After 30min incubation, 4x washing steps consisting of addition of 1ml Cell Staining Buffer (BioLegend, Cat#: 420201) followed by centrifugation at 300g for 10min at 4°C were performed, with the cells before the last wash being filtered through a blue lid filter cap back. Subsequently cells were counted and then processed further for cDNA library preparation.

### Source of surgical central nervous system tumor specimens

Brain tumor samples were collected intraoperatively and given a 4-digit, de-identified code by the Honest Broker in the Bartoli Brain Tumor Laboratory in accordance with IRB-AAAA4666. De-identified samples were handed off only once a satisfactory histological diagnosis had been made. Samples were transported fresh in saline on ice to the De Jager laboratory where further analyses were conducted under the IRB protocol AAAR4962. The Bartoli Brain Tumor Laboratory is funded through Columbia University Medical Center’s Department of Neurological Surgery.

### CITE-Seq staining of cell preparations

Cells were counted, cell viability was assessed and subsequently 0.7×10^6^ cells were transferred to a 5ml low-protein binding tube (fisher scientific; Cat#: 13-864-407). 1ml cold PBS (Corning; Cat#:21-040-CV) was added, cells filtered through a blue lid filter (FACS tube) back into the 5ml low-protein binding tube (fisher scientific; Cat#: 13-864-407), centrifuged at 300g, 4°C for 10min with repetition of this washing step. Supernatant was then removed using a pipette tip. Cell pellet was then resuspended in 22.5µL of Cell Staining Buffer (BioLegend, Cat#: 420201), 2.5µL of Human FcBlock (BioLegend, Cat#: 422301) was added and cells were incubated for 10min on ice. For hashing, hashing master mix was prepared as follows: 2µl of TS-A hashtag antibody (0.5 mg/ml, BioLegend, TotalSeq- A0251 anti human Hashtag1, Cat#: B364030; TotalSeq- A0252 anti human Hashtag2, Cat#: B369100; TotalSeq- A0253 anti human Hashtag3, Cat#: B378300; TotalSeq- A0254 anti human Hashtag4, Cat#: B381073) was added to 20µl of Cell Staining Buffer (BioLegend, Cat#: 420201), mixed and passed through a 0.1µm filter at 12000g for 2 min at 4°C. Then, 6.25µl of reconstituted combined CITE-Seq cocktail were added to 18.75µl of hashtag master mix and the combination was added to

the cells incubating in FcBlock (BioLegend, Cat#: 422301) and Cell Staining Buffer (BioLegend, Cat#: 420201). If samples were not hashed, 6.25µl of reconstituted combined CITE-Seq cocktail were added to 18.75µl of cell staining buffer (BioLegend, Cat#: 420201) and the combination was added to the cells incubating in FcBlock (BioLegend, Cat#: 422301) and Cell Staining Buffer (BioLegend, Cat#: 420201). After 30min incubation, 4x washing steps consisting of addition of 1ml Cell Staining Buffer (BioLegend, Cat#: 420201) followed by centrifugation at 300g for 10min at 4°C were performed, with the cells before the last wash being filtered through a blue lid filter cap back. Subsequently cells were counted and then processed further for cDNA library preparation.

### Library preparation and CITE-Seq of different human microglia and PBMC preparations – without Cell hashing

Detailed single cell RNA sequencing procedures were performed following the Biolegend TotalSeq™-A Antibodies and Cell Hashing with 10x Single Cell 3’ Reagent Kit v3.1 (Dual Index) Protocol (https://www.biolegend.com/fr-ch/protocols/totalseq-a-dual-index-protocol) with minor modifications.

Briefly, for PV044, 045, 049, and 051-055, TotalSeq™-A antibody-labeled cells were counted using a Nexcelom Cellometer Vision 10x objective and AO/PI stain. 20µl of AO/PI were mixed with 20µl of cell suspension and 20µl were loeaded onto a standard thickness Cellometer cell counting chamber (Nexcelom, Cat#: CHT4-SD100-002) using a dilution factor set to 2.

Single cell libraries were constructed using 10x Chromium Next GEM Single Cell 3’ Reagent Kits v3.1 (Dual Index) with Feature Barcode technology for Cell Surface Protein (10x Genomics, Pleasanton, CA) according to the manufacturer’s protocol with modification. Briefly, a total of approximately 10,000 cells were loaded into one well of the Chip G kit for GEM generation using a 10x Genomics chromium controller single-cell instrument. Reverse transcription reagents, barcoded gel beads, and partitioning oil were mixed with the cells to generate single-cell gel beads in emulsions (GEM). Within one GEM, all generated cDNA shares a common 10x barcode. Libraries were generated and sequenced from cDNA, and 10x barcodes were used to associate individual reads back to individual partitions.

Incubation of the GEM then generated barcoded cDNA from polyadenylated mRNA and simultaneous generation of DNA from the same single cell within the GEM from cell surface protein Feature Barcode. ADT-added primers (0.2µM stock) were used to generate feature barcoded-DNA during complementary DNA (cDNA) amplification to enrich TotalSeq-A cell surface oligonucleotides. After incubation, GEMs were broken, and pooled fractions were recovered. Cell barcoded cDNA molecules were amplified by PCR and used to construct

libraries. The amplified cDNA was then separated by SPRI size selection into cDNA fractions containing mRNA derived cDNA (>400bp) and ADT-derived cDNAs (<180bp) and further purified by additional rounds of SPRI selection.

During sample index amplification, independent sequencing libraries were generated from the mRNA and ADT cDNA fractions using Dual-Index Kit TT, Set A (PN-1000215) and Dual Index ADT (DI_ADT*x*) i5/i7 Primer Pairs (10µM Stock) to enrich TotalSeq-A cell surface libraries. These libraries were analyzed and quantified using TapeStation D5000 screening tapes (Agilent, Santa Clara, CA) and Qubit HS DNA quantification kit (Thermo Fisher Scientific). The libraries were pooled and sequenced on a NovaSeq 6000 with S4 flow cell (Illumina, San Diego, CA) at a ratio of GEX:ADT = 2:1. Paired-end, dual-index sequencing was used to perform 28 cycles for read 1, 10 cycles for i7 index, 10 cycles for i5 index, and 90 cycles for read 2.

**cDNA Primers**

- ADT cDNA PCR additive primer: 5’CCTTGGCACCCGAGAATT*C*C

**Primers Used for Sequencing Library Construction**

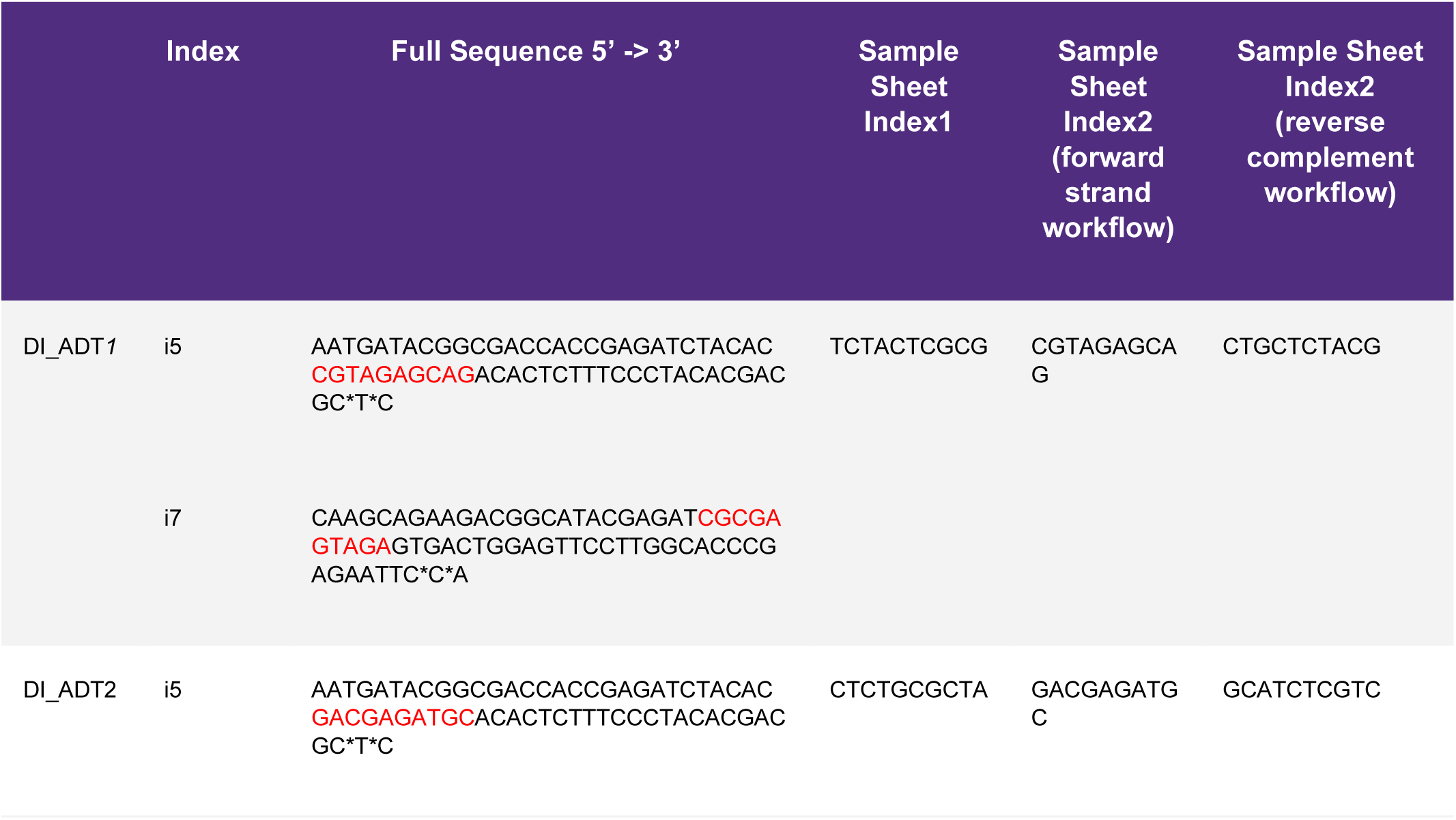
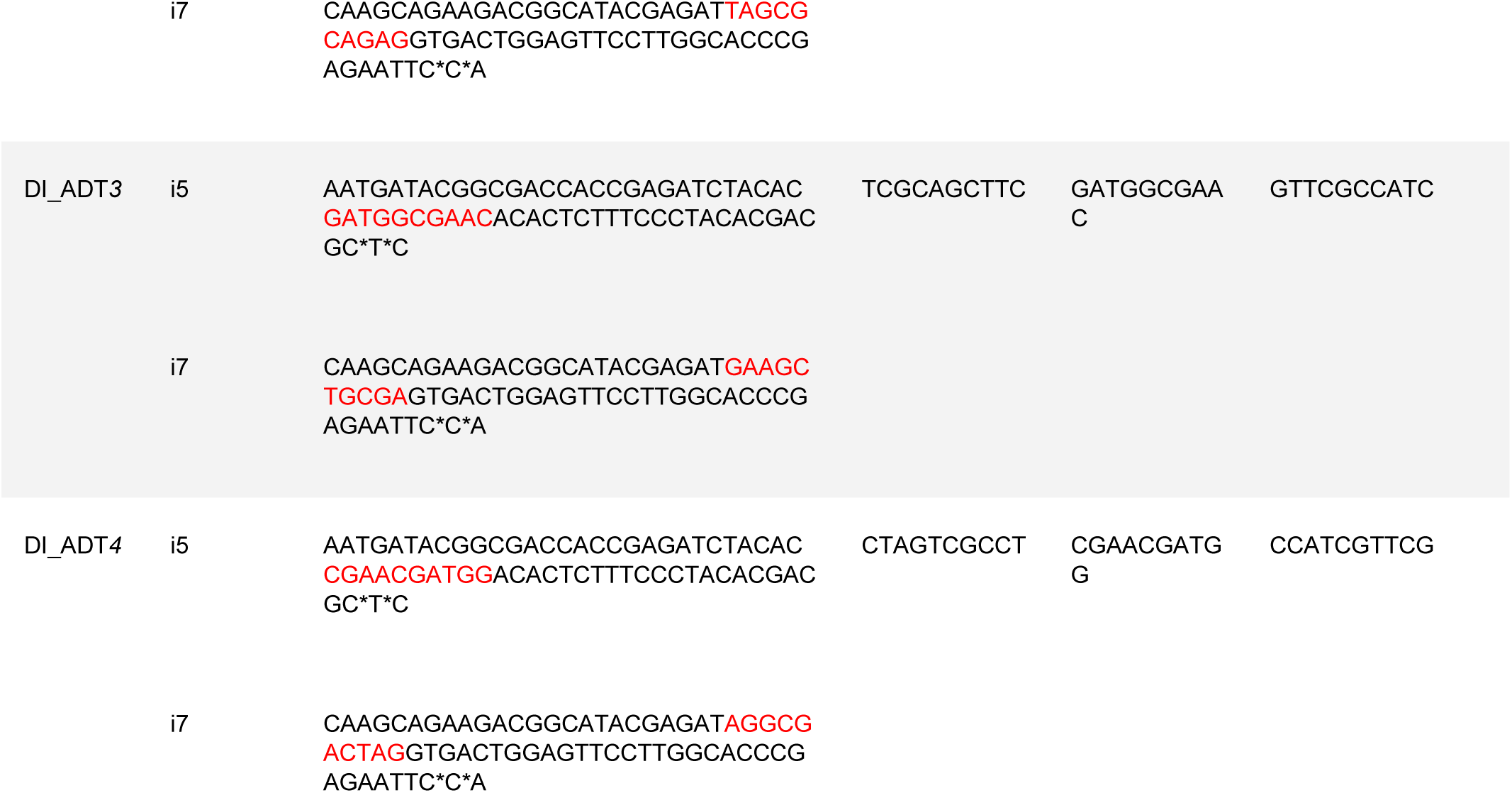

### Library preparation and CITE-Seq of different human microglia and PBMC preparations with Cell hashing

Detailed single cell RNA sequencing procedures were performed following the Biolegend TotalSeq™-A Antibodies and Cell Hashing with 10x Single Cell 3’ Reagent Kit v3.1 (Dual Index) Protocol (https://www.biolegend.com/fr-ch/protocols/totalseq-a-dual-index-protocol) with minor modifications.

Briefly, TotalSeq™-A antibody-labeled and hashed microglia cells from different brain region of the same donor were pooled, then counted using a Nexcelom Cellometer Vision 10x objective and AO/PI stain. 20µl of AO/PI were mixed with 20µl of cell suspension and 20µl were loeaded onto a standard thickness Cellometer cell counting chamber (Nexcelom, Cat#: CHT4-SD100-002) using a dilution factor set to 2.

Single cell library generation was constructed using 10x Chromium Next GEM Single Cell 3’ Reagent Kits v3.1 (Dual Index) with Feature Barcode technology for Cell Surface Protein (10x Genomics, Pleasanton, CA) according to the manufacturer’s protocol with modification. Briefly, a total of approximately 10,000 – 20,000 cells were loaded into one well of the Chip G kit for GEM generation using a 10x Genomics chromium controller single-cell instrument. Reverse transcription reagents, barcoded gel beads, and partitioning oil were mixed with the cells to generate single-cell gel beads in emulsions (GEM). Within one GEM, all generated cDNA shares a common 10x barcode. Libraries were generated and sequenced from cDNA, and 10x barcodes were used to associate individual reads back to individual partitions.

Incubation of the GEM then generated barcoded cDNA from polyadenylated mRNA and simultaneous generation of DNA from the same single cell within the GEM from cell surface protein Feature Barcode. ADT additive primer (0.2µM Stock) and HTO additive primer v2 (0.2µM Stock) were used to generate feature barcoded-DNA during complementary DNA (cDNA) amplification to enrich TotalSeq-A cell surface oligonucleotides. After incubation, GEMs were broken, and pooled fractions were recovered. Cell barcoded cDNA molecules were amplified by PCR and used to construct libraries. The amplified cDNA was then separated by SPRI size selection into cDNA fractions containing mRNA derived cDNA (>400bp) and ADT/HTO-derived cDNAs (<180bp) and further purified by additional rounds of SPRI selection.

During sample index amplification, independent sequencing libraries were generated from the mRNA, ADT and HTO cDNA fractions using Dual-Index Kit TT, Set A (PN-1000215), Dual Index ADT (DI_ADT*x*) i5/i7 Primer Pairs (10µM Stock) and Dual Index HTO (DI_HTO*x*) i5/i7 Primer Pairs (10µM Stock). These libraries were analyzed and quantified using TapeStation D5000 screening tapes (Agilent, Santa Clara, CA) and Qubit HS DNA quantification kit (Thermo Fisher Scientific). The libraries were pooled and sequenced on a NovaSeq 6000 with S4 flow cell (Illumina, San Diego, CA) at a ratio of GEX:ADT:HTO = 8:4:1. Paired-end, dual-index sequencing was used to perform 28 cycles for read 1, 10 cycles for i7 index, 10 cycles for i5 index, and 90 cycles for read 2.

**cDNA Primers**

- ADT cDNA PCR additive primer: 5’CCTTGGCACCCGAGAATT*C*C
- HTO cDNA PCR additive primer: v2 5’GTGACTGGAGTTCAGACGTGTGCTCTTCCGAT*C*T

**Primers Used for Sequencing Library Construction**

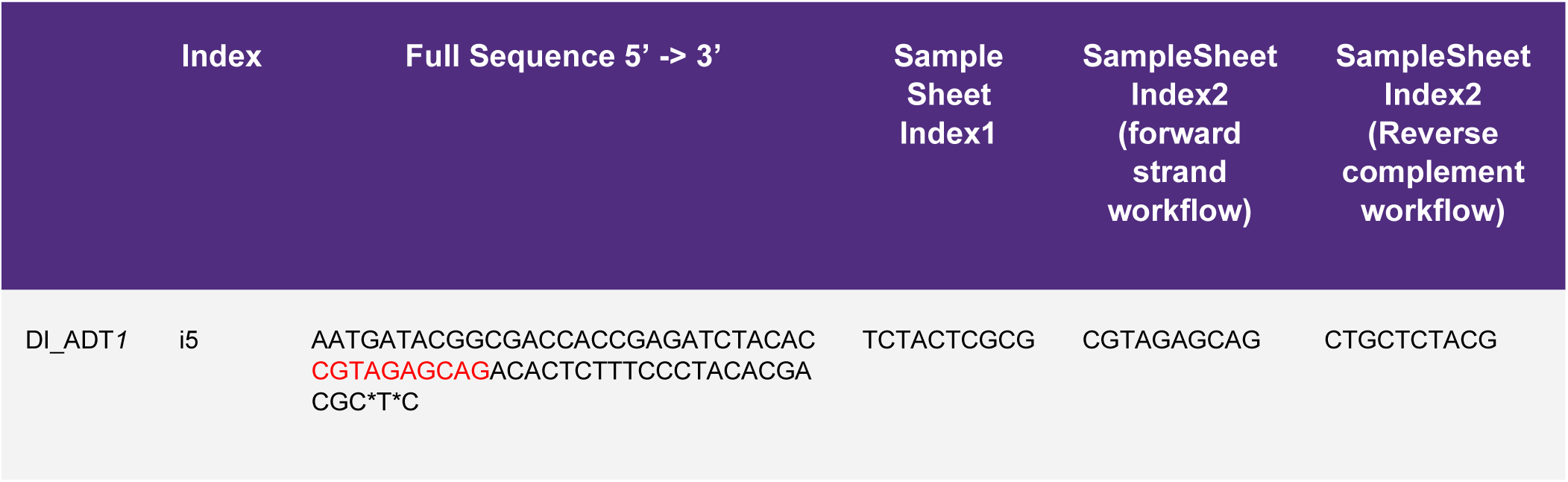
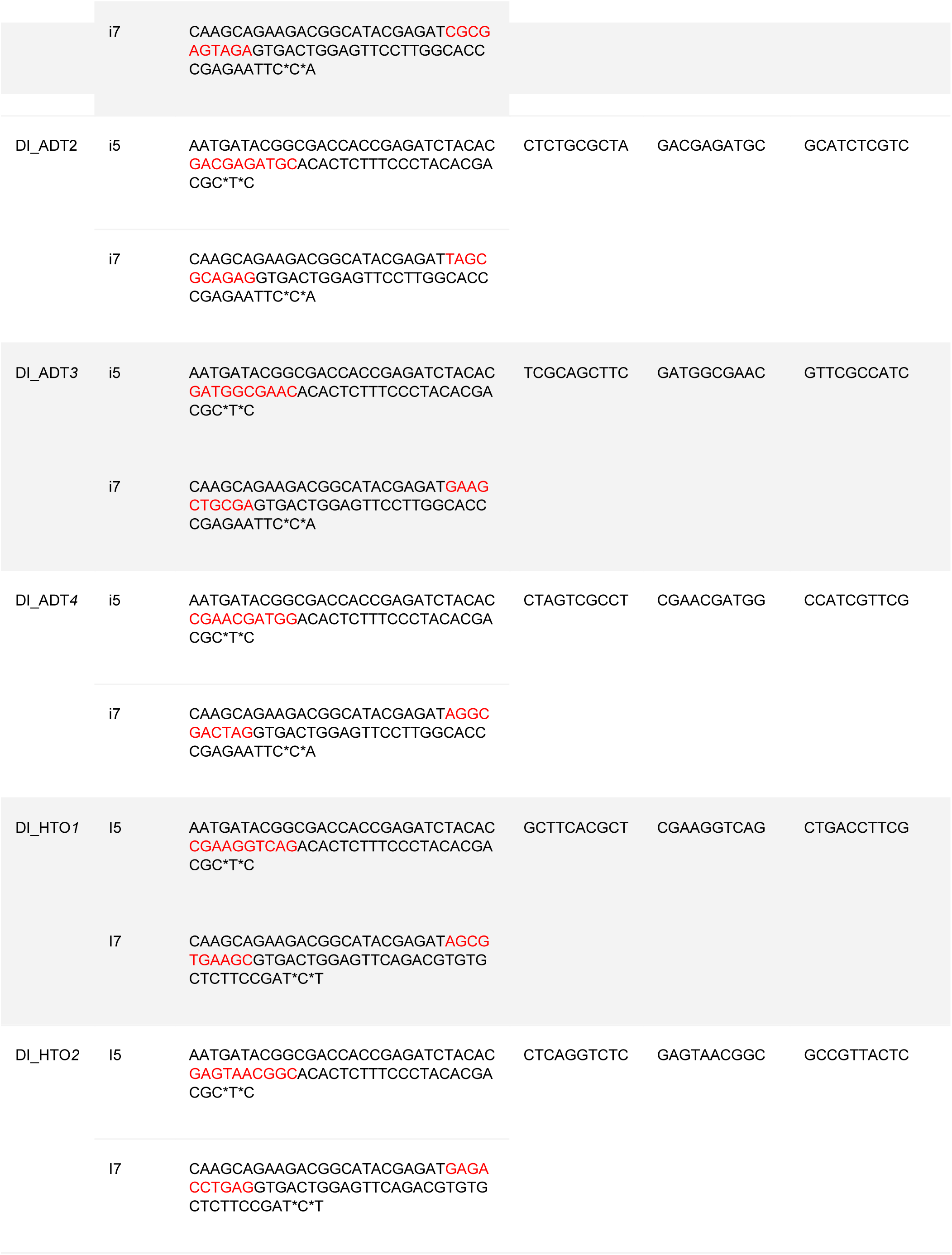
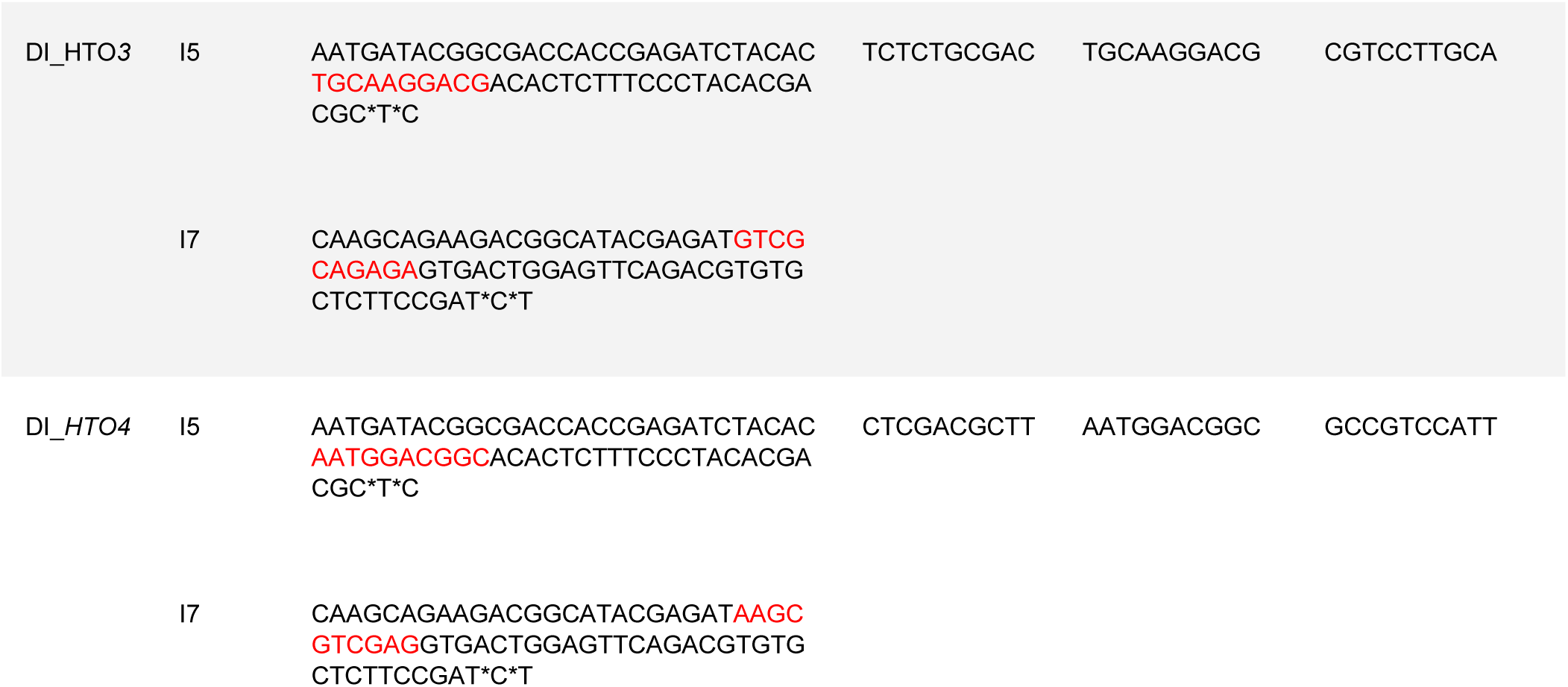

### Quality control and analysis of CITE-Seq data

#### CITE-Seq data preprocessing

The raw single-cell sequencing reads of gene expression libraries and antibody-derived tag libraries were processed using the Cell Ranger software from 10x Genomics (version 7.2.0). The computational analysis was conducted on a high-performance computing (HPC) cluster configured with 24 GB of virtual memory and four local cores. The GRCh38 human reference transcriptome (version GRCh38-2020-A) was employed for alignment, barcode processing, and UMI counting. A custom feature reference file was used for detecting unique molecular identifiers (UMIs) associated with antibody-derived tags.

#### Quality control of cells

The resulting filtered feature barcode matrix for each sample was loaded into Rstudio environment for preprocessing and quality control. The analysis utilized the Seurat and dplyr packages for the data analysis and the ggplot2 and viridis packages for the downstream visualizations. A Seurat object was created using the gene expression matrix as well as an assay object for the ADT data to allow separate analysis of antibody-derived tags. Sum and median of UMI counts per antibody were computed for to assess the distribution across cells. These counts were plotted using log-scale transformations. Hashtag ADT data underwent normalization using the Center Log Ratio (CLR) method.

By visual inspection of violin and UMAP plots focusing on mitochondrial percentage, RNA feature counts, and ADT counts, thresholds were established to exclude low quality/dying cells from further analysis. Cells with low RNA context formed clear clusters and partially overlapped with high-MT cells. These cells were excluded. Cells with low ADT counts also largely overlapped low RNA cells. Additionally, high ADT cells were also excluded. ADT counts were calculated on a log scale due to the presence of extreme outliers. Following this, the datasets underwent normalization. RNA data normalization was performed using Seurat’s default method, while ADT data underwent CLR normalization to prepare for dimensional reduction. Principal component analysis for dimensionality reduction, followed by UMAP visualization, was used to identify distinct cell populations.

#### Selection of usable antibodies

Our CITE-Seq antibodies contain the TotalSeq™-A Human Universal Cocktail, which is designed for PBMCs. They include antibodies against proteins that are not expressed in microglia. To exclude such antibodies from downstream analysis, we first chose a threshold of protein expression levels based on our PBMC CITE-Seq library PV049, and then applied the threshold to our microglia CITE-Seq libraries to select proteins expressed in microglia.

To choose a threshold of protein expression levels, we annotated cell types in PV049 based on the RNA expression using the R package Azimuth version 0.5.0^73^. For each of the level 2 cell types, we computed pseudobulk protein expression using NormalizeData() with the option normalization.method = “LogNormalize” and PseudobulkExpression() of the R package Seurat version 5.0.3. The pseudobulk protein levels of CD45, CD3, CD4, CD8, CD19, CD20, T-cell receptors, nine isotype controls, HLA class II, and human immunoglobulins were examined in the level 2 cell types. We determined that 40.0 is a reasonable threshold because binary thresholding of the aforementioned proteins was mostly consistent with known expression patterns in the cell types. Notable exceptions were IgE, IgL, and IgM as they were much higher than 40.0 in many non-B cell types.

To select proteins that are expressed in microglia, we computed pseudobulk protein levels per our microglia CITE-Seq libraries. We then tested each protein whether its pseudobulk was higher than the threshold 40.0 in at least one of the libraries. If true, the protein was considered as expressed in microglia and used in downstream analysis. As an exception, 17 proteins in our microglia-specific CITE-Seq panel were retained in the downstream analysis regardless of their pseudobulk level. Also, IgE, IgL, and IgM were excluded because of their anomalous expression pattern. Consequently, 101 proteins were selected for downstream analysis.

#### Protein-RNA correlations

For each protein-RNA pair, the Pearson correlation was computed by comparing the average RNA level and the average protein level across microglia libraries. Protein names were paired with RNA names based on Ensembl gene IDs. The RNA expression levels were computed by applying NormalizeData() to an RNA UMI count matrix with the option normalization.method = “LogNormalize”. For the protein expression levels, we applied four different normalization methods to an ADT UMI count matrix: (1) NormalizeData() with normalization.method = “LogNormalize”, (2) NormalizeData() with normalization.method = “CLR” and margin = 1, (3) NormalizeData() with normalization.method = “CLR” and margin = 2, and (4) DSBNormalizeProtein() of R package “dsb” (version 1.0.3)^74^. When we compared the four normalizations with respect to the median protein-RNA correlation, the “LogNormalize” approach had the highest median correlation. Therefore, we employed “LogNormalize” for normalizations of both RNA and ADT in all of the subsequent analyses.

#### UMAP embedding

Cells were embedded into the protein UMAP space and the RNA UMAP space separately using the standard procedure of Seurat with the top 10 principal components. The protein UMAP was computed using the above-defined 101 proteins expressed in microglia. The R package Harmony (version 1.2.0) was used to integrated cells from different libraries.

#### Clustering of microglia in glioblastoma data

Two glioblastoma libraries (GBM I and GBM II) were selected for further analysis because of their high cell viability and abundance of QC-passed cells. The two GBM libraries were integrated based on their RNA expression using Harmony. The cells were classified into three clusters by running FindClusters() in the Harmony space at resolution = 0.05. Differentially expressed RNAs between clusters were identified using FindMarkers(). Expression levels of marker genes for microglia, macrophage, and proliferating cells were examined to identify the cluster of microglia.

#### scHPF projection

After quality control, each of the query libraries was projected into a pre-existing consensus scHPF^75^ model. This model captures 23 non-orthogonal, latent factors representing microglial gene expression signatures defined in a dataset of 378,543 live human microglia from 127 donors and 17 unique brain tissue types, sequenced using 10x v3 chemistry. Broadly, the model represents key biological processes in microglia such as metabolism, phagocytosis, antigen presentation, disease-associated states.

For each query library, the raw UMI counts were extracted and downsampled to match the sequencing depth of the model using scHPF’s downsample_loom.py. All 8,478 genes in the scHPF model were present in the projected libraries. The downsampled counts were then projected into the model (scHPF’s prep-like, project, score). The resulting factor scores were projected into the reduced dimensional UMAP space of the scHPF model using umap_transform from the R package uwot^76^.

#### Label transfer to single-cell CITE-seq microglial datasets

We applied a label transfer approach as described in our prior publication, using that data as the reference^20^. In short, we concatenated the query and reference data in a single Seurat object and the unique differentially expressed genes from our pairwise differential expression testing (Identification of cluster-defining gene sets) were used for cross-batch merging with the fastmNN algorithm. For all these analyses, we used 40 components. The normalized “mnn.reconstructed” assay, which represents per-gene corrected log-expression values, was used for downstream analysis. We leveraged a combinatorial workflow leveraging two distinct models for different clusters showed the best accuracy: a set of pairwise support vector machine (SVM) classifiers using consensus voting to assign labels for the smaller clusters (8- 12), and a flat XGBoost^77^ (XGB) classifier to assign labels for the larger clusters (1-7) with higher transcriptional homology. Predictions from these two models were integrated to achieve higher predictive accuracy.

The overall workflow for both methods was similar: as a few our classes are transcriptionally similar, similar classes are condensed (clusters 1/6/7 and clusters 2/4), then a subset of the cells in our dataset are selected for training. Next, the differentially expressed genes from our pairwise differential expression testing (Identification of cluster-defining gene sets) were selected as the features for training and PCA was performed on the resulting subset of the data. For the SVM, the training subset was 0.2 for classes 1-9, and 0.5 for classes 10-12. A separate classifier was trained for each unique pair of clusters (i.e. a classifier to compare clusters 1/6/7 and 2/4, 1/6/7 and 3….1/6/7 and 12, then 2/4 and 3, 2/4 and 5….2/4 and 12) using only the genes found to be differentially expressed (both up and down) between that specific pair of clusters. Data classes were then rebalanced using combined over/under resampling to reduce class imbalance for smaller classes. Caret^78^ was used to perform PCA and hyperparameter optimization of a SVM model using a radial kernel and 10-fold cross-validation repeated three times. PCA was conducted independently during each fold. Conversely, for XGB, the training subset was 0.33, and the model trained only on cells from groupings 1/6/7, 2/4, 3, and 5. Similarly, PCA was performed upstream on the subset of scaled data consisting of all genes found to be differentially expressed between any clusters. Hyperparameter optimization with 5-fold validation was performed in a stepwise fashion: tree number was first optimized, then tree-specific parameters were tuned with a restrictive grid search, then regularization parameters were tuned with a restrictive grid search, then final optimization was conducted with grid search in a narrow range around prior optimal parameters.

To construct a validation subset, a subset of 50% of the dataset was sampled exclusively from cells not used for training of either the SVM or XGB models. The same scaling and sub-setting operations described above were applied to this data. Optimized SVM and XGB models were used to classify the data. For SVM models, final classifications were obtained with hard consensus voting, as the class with the majority of votes was chosen as the final class of the SVM voting ensemble. Similarly, for XGB, which outputs a probability for each class summing to 1 across all classes, the highest probability was used to choose the assigned label. However, the class probabilities for XGB also provided the opportunity to evaluate the confidence of the classifier and drop lower-confidence assignments. As such, cells were only retained for SVM classifications in classes 10-12 or for XGB classifications in classes 1-7 that had higher than 50% classification probability for the assigned probability. Final classifications were merged across datasets, and accuracy was evaluated by examining sensitivity, specificity, and congruence of marker gene expression patterns of cells assigned to each class with marker gene expression patterns seen in our original data. Identical procedures were performed for query datasets. Because feature scaling is applied in this workflow, and different data sources lead to dramatically different RNA expression patterns, different data sources were aggregated and processed separately in this workflow. The three data sources that were annotated independently were: human microglia, iMG, and HMC3. Prediction confidence was in line with prior label transfer experiments performed in our prior publication^20^.

#### Decision Tree and *in silico* sorting of validation datasets consisting of freshly isolated human microglia from GBM IV-VII

To assess the feasibility of the enrichment of high-scHPF cells for a given scHPF factor based on surface protein abundance, a systematic approach involving protein quantification, gating, and statistical modeling was employed. The datasets (GBMI-II, IV-VII), which included human microglia samples, were preprocessed by filtering for human microglia based on metadata annotations. The expression data matrix was then transformed into quantiles, ranking protein abundances across all samples within each library. Gating was performed to isolate cells based on protein abundance, with a custom filtering method used to select cells with protein abundance values either above or below a specified quantile threshold. These thresholds, defined using protein abundance quantiles, guided the gating process to retain either the top or bottom tail of the distribution based on the desired direction of enrichment, iteratively applying this for each of the top proteins identified by a decision tree model. For each of scHPFs, cells were classified into binary levels “scHPF high” and “scHPF low” depending on whether their scHPFs were higher than median + median absolute deviation (MAD).

The classification model for predicting the high or low status of scHPF based on protein expression was constructed using the decision tree algorithm in the R package “rpart” (version 4.1.24). The decision tree was trained on quantile-transformed discovery data (GBM I-II) and the binary classification of scHPF status with the maximum probability split from the tree used to define the gating thresholds. This was done by running the rpart() function with parameters maxdepth = 3 and cp = -1 for each scHPF. In our decision tree model, the leaf with the highest scHPF-high proportion in the discovery data is only the node that emits “scHPF-high” status, and all the other leaves predict “scHPF-low” status. After applying the gating thresholds for three proteins, the number of high-scHPF cells in both the discovery (GMB I-II) and validation datasets (GBM IV-VII) was quantified, and the proportion of high-scHPF cells before and after gating was calculated. Comparisons of these proportions were made between both datasets to evaluate the effect of gating. The results were summarized in a table that presented the total number of cells and the proportion of high-scHPF cells, both with and without gating, across all steps (**Table S3**).

#### Immunohistochemistry staining for CD51

For validation immunostaining, six μm formalin-fixed paraffin-embedded (FFPE) tissue sections from the frontal cortex of fifteen moderate and end-stage Alzheimer’s disease- diagnosed individuals were obtained from the New York Brain Bank at Columbia University. The tissues were stained with CD51 (1:100, 488 channel, Invitrogen, cat.# PA5-80745) and IBA1 (1:100, 647 channel, Wako, cat.# 01919741).

The FFPE tissue sections were deparaffinized using CitriSolv (d-limonene, Decon Laboratories, cat.# 1601H) as a clearing agent for 20 minutes. The sections were rehydrated and prepared for staining through a series of graded ethanol washes.

Heat-mediated antigen retrieval was performed with citrate buffer (pH=6, Sigma- Aldrich, cat.# C9999) using a microwave (800W, 30% power setting) for 25 minutes. Following this, the sections were blocked for 30 minutes at room temperature (RT) using a Bovine Serum Albumin-blocking medium (3%, Sigma-Aldrich, cat.# A7906) to minimize non-specific antibody binding. The sections were incubated overnight with the primary antibodies (anti-CD51 and anti-IBA1) at 4°C. After washing, the tissues were incubated for one hour with fluorophore-conjugated secondary antibodies (1:500, Alexa Flour 488 and 647, Invitrogen, cat.# A21206, A21202, A21447) to bind to the primary antibody for protein detection and signal enhancement. After incubation, the sections were washed and treated with True Black Lipofuscin Autofluorescence Quencher (Biotium, cat.# 23007) for 2 minutes at RT to minimize endogenous autofluorescence. An anti-fading DAPI mounting agent (Invitrogen, cat.# P36931) was used to coverslip.

#### Image acquisition and CellProfiler analysis for object intensity quantification

Imaging was performed using a Nikon NI Eclipse microscope at 20X magnification with the NIS-Elements Advanced Research software (v5.21.03). A Hamamatsu Orca-Fusion Digital Camera (C14440) was utilized to capture 40 images for each sample in a systematic zigzag pattern, ensuring coverage of all cortical layers for post-acquisition analysis. Analysis was performed using CellProfiler (v4.2.5)^75^ with a customized pipeline to segment DAPI+IBA1+ cells and measure the mean intensity of CD51.

First, nuclei (DAPI) were segmented using the “IdentifyPrimaryObjects” module with advanced settings. The diameter of the objects was set between 18 and 80 pixels, and the “Robust Background” method was used for thresholding. Next, IBA1 was segmented with the “EnhanceOrSuppressFeatures” module to augment the elongated morphology of ramified microglia, specifically using the ’Line structures’ method. IBA1+ cells were detected using the “IdentifyPrimaryObjects” module, with the diameter of objects set to range from 10 to 300 pixels. Importantly, the “SplitOrMergeObjects” module was used to group small structures, such as the processes of ramified microglia, and assign them to nearby objects. IBA1+DAPI- cells were filtered out using the “RelateObjects” module, with DAPI defined as the parent objects and IBA1 as the child objects. Subsequently, the size and shape of the merged IBA1- DAPI objects were quantified by calculating Zernike features, alongside measuring the object intensity of CD51 staining.

## Supporting information

Table S1_Tissue Samples

Table S2A_Human Microglia Proteomics_Raw Data

Table S2B_HMC3 Proteomics

Table S2C_iMG Proteomics

Table S3_Heatmap_new_no ET

Table S4A_Correlation ADT scHPF_discovery

Table S4C_scHPF_in silico gating

Table S4A_Correlation ADT scHPF_discovery

Table S5_ADT_RNA_DEG_Mic_Mac

Table S4B_Correlation ADT scHPF_validation

## Acknowledgements

Study data were generated from postmortem brain tissue provided by the Religious Orders Study and Rush Memory and Aging Project (ROSMAP) cohort at Rush Alzheimer’s Disease Center, Rush University Medical Center, Chicago, the New York Brain Bank (NYBB) and the University of Washington, Seattle, Brain Laboratory.

Study data were further generated from surgical tissue specimens provided by the Bartoli Brain Tumor Laboratory at Columbia University which is funded through Columbia University Medical Center’s Department of Neurological Surgery.

The part of the proteomics work was performed in the Environmental Molecular Sciences Laboratory, a national scientific user facility sponsored by the Department of Energy and located at Pacific Northwest National Laboratory, which is operated by Battelle Memorial Institute for the Department of Energy under Contract DE-AC05-76RL0 1830.

The work performed at CTCN, Columbia University was supported by the Chan-Zuckerberg Initiative’s Neurodegeneration Challenge Network grant CS-02018-191971. Some of the work also emerged from support from NIH/NIA grants R01 AG070438, U01 AG061356, RF1 AG057473, R01AG048015. Research reported in this publication was supported by the National Institute of General Medical Sciences of the National Institutes of Health under Award Number T32GM007367 and by the National Cancer Institute of the National Institutes of Health under Award Number F30CA261090. This work was further supported by a Medical Research Council Programme Grant (MR/Y014847/1) to B.D.S.

Research reported in this publication was partially performed in the Columbia Center for Translational Immunology and P&S Flow Cytometry Core, the Columbia Stem Cell Initiative vcb-Flow Cytometry Core at Columbia University and the Proteomics and Macromolecular Crystallography Core in the Columbia University Herbert Irving Comprehensive Cancer Center. AAS is supported by The Thompson Foundation (TAME-AD) and the Henry and Marilyn Taub Foundation. We further thank the New York Genome Center for a long-standing collaboration and efforts in RNA-Sequencing for the study.

We thank Berke Karaahmet for help with the visualization of Figure 5E using ggPlot2 and R Studio. All illustrations were created with BioRender.com

## Materials Availability statement

This study did not generate new unique reagents.

## Data and Code Availability statement

Sequencing and proteomics data reported in this paper were deposited in the AD knowledge portal and are accessible from the following website: *upload in progress*.

Code is shared in the GitHub repository, accessible through the following link: https://github.com/masashi-CU/Development-of-a-human-microglia-specific-CITE-seq-panel

## Supplementary Tables

**Table S1:** Tissue samples

**Table S2A:** human microglia proteomics dataset

**Table S2B:** HMC3 Proteomics

**Table S2C:** iMG Proteomics

**Table S3:** Heatmap

**Table S4A:** Correlation ADT scHPF Discovery Dataset

**Table S4B:** Correlation ADT scHPF Validation Dataset

**Table S4C:** scHPF *in silico* gating

**Table S5:** Differentially expressed genes/proteins (RNA/ADT) between microglia vs. macrophages isolated from human GBM

## Author contributions

Conceptualization, P.L.D., V.H.;

Methodology: P.L.D., V.H. and J.F.T.;

Investigation: V.H., A.R.B., R.C., T.L., J.P., A.B., J.L.F., I.H., R.P., R.P., G.T., H.T., Y.Z., L.C., P.L.D.;

Data Curation: J.F.T., Y.Z., M.F., P.L.D.;

Formal Analysis: V.H., J.F.T., V.M., N.A.P., S.S.K., M.F., P.B., L.Z., Y.L., P.L.D.;

Writing – Original Draft: V.H., P.L.D.;

Writing – Review & Editing: V.H., P.L.D;

Funding Acquisition: P.L.D;

Resources: J.N., L.D.W., J.S., A.T., R.A.S., N.A.S., T.L.Y-P., C.R., D.A.B., P.C., J.N.B., A.H., A.F.L., B.S., F.S., A.S., V.A.P., P.L.D;

Supervision: Z.Y., M.T., M.F., V.H., V.A.P., P.L.D.

