## Supplementary Figures for "A proteogenomic tool uncovers protein markers for human microglial states"

### Supplementary Figure 1

#### A Proteomics of freshly isolated human microglia

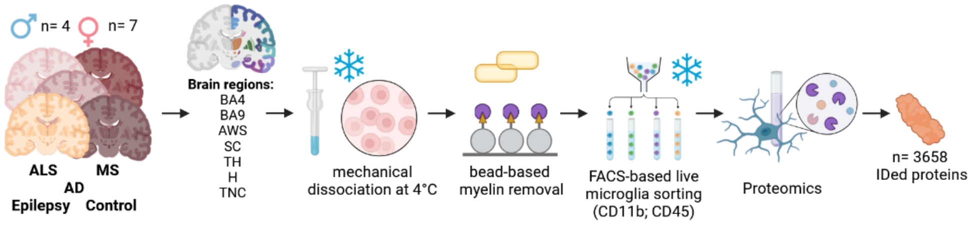


#### B PCA Plot

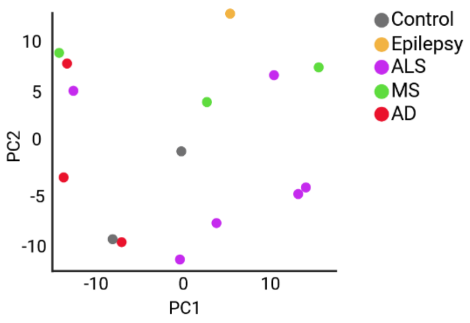


C

**HMC3**

CD44
ITGA5
CD276
TMX3
LGALS3
PTPRG
PLXNB2
PTPRJ
ABCB11
HLA-C
ITGB5
CD40
ABCB1 HLA-
A L1CAM
HLA-B
PSEN1 APP
SORT1
PICALM
ITGB1
ITGA6
SLC2A1
SIRPA
CD47 HLA-
E CDH2
ITGAV
ADAM10
FERMT2
SLC7A5
TSPO
SLC1A4
NPTN
ITGB3 ANO6

D

**iMG II**

| | |
|---|---|
| CD4 | PLXNB2 |
| ITGAM | HLA-B |
| FCGR3A | HLA-DRA |
| ITGB2 | HLA-DPB1 |
| ITGB1 | HLA-C |
| PECAM1 | BST2 |
| PTPRC | PTPRJ |
| CD86 | SORL1 |
| CD40 | SORT1 |
| HLA-A | APP |
| ITGA6 | ITGAV |
| CD47 | ITGA5 |
| CD276 | SLC2A1 |
| CR1 | ITGB5 |
| CD163 | SLC1A4 |
| ITGB3 | ANO6 |
| CD44 | TMX3 |
| HLA-E | NPTN |
| ITGAL | ABCA7 |
| CDH2 | PSEN1 |
| CD33 | PICALM |
| CD14 | ACE |
| ITGAX | PTPRG |
| SIRPA | HLA-DRB5 |
| P2RY12 | PILRA |
| ADAM10 | LGALS3 |
| | TSPO |

**iMG III**

| | |
|---|---|
| CD44 | HLA-DRB5 |
| ITGA5 | SORT1 |
| SORL1 | HLA-DRA |
| NCAM1 | PICALM |
| PILRA | ITGB1 |
| ITGB2 | FCGR3A |
| CR1 | ITGA6 |
| CD276 | PSEN2 |
| TREM2 | CD86 |
| TMX3 | SLC2A1 |
| LGALS3 | PTPRC |
| PTPRG | CD14 |
| PLXNB2 | SIRPA |
| ITGAX | HLA-DPB1 |
| PTPRJ | CD47 |
| HLA-C | CD33 |
| CX3CR1 | HLA-E |
| ITGB5 | ITGAV |
| ITGAL | ADAM10 |
| CD4 | P2RY12 |
| CD40 | SLC7A5 |
| MS4A6A | TSPO |
| HLA-A | SLC1A4 |
| ITGAM | ICAM1 |
| CD163 | SLC46A1 |
| PECAM1 | NPTN |
| BST2 | ITGB3 |
| HLA-B | ANO6 |
| ABCA7 | |
| PSEN1 | |
| APP | |

E

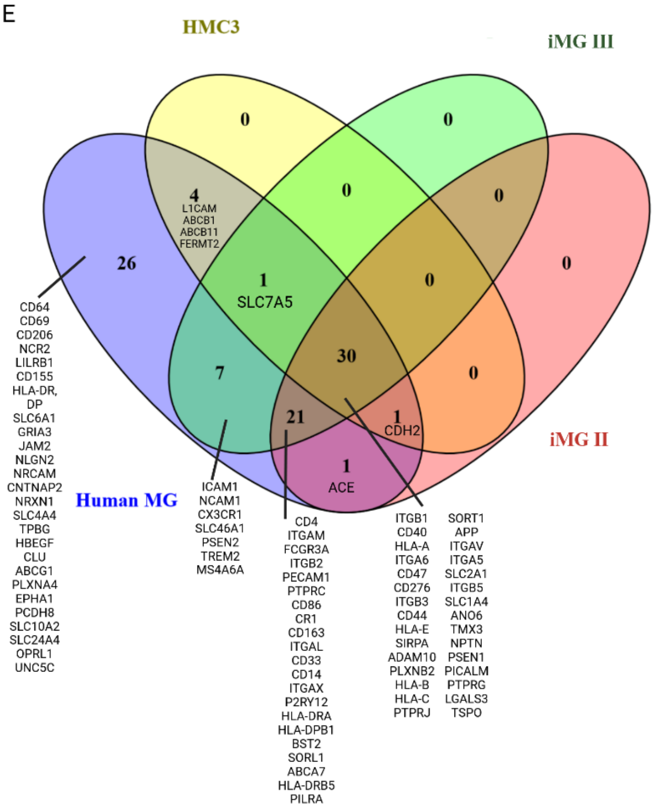


| | |
|---|---|
| **Candidates expressed by HMC3s** | 36 |
| **Candidates expressed by iMGs (Sher Lab)** | 53 |
| **Candidates expressed by iMGs (Sproul Lab)** | 59 |
| **Candidates expressed only by freshly isolated MG** | 26 |
| **Total candidates** | 91 |

F

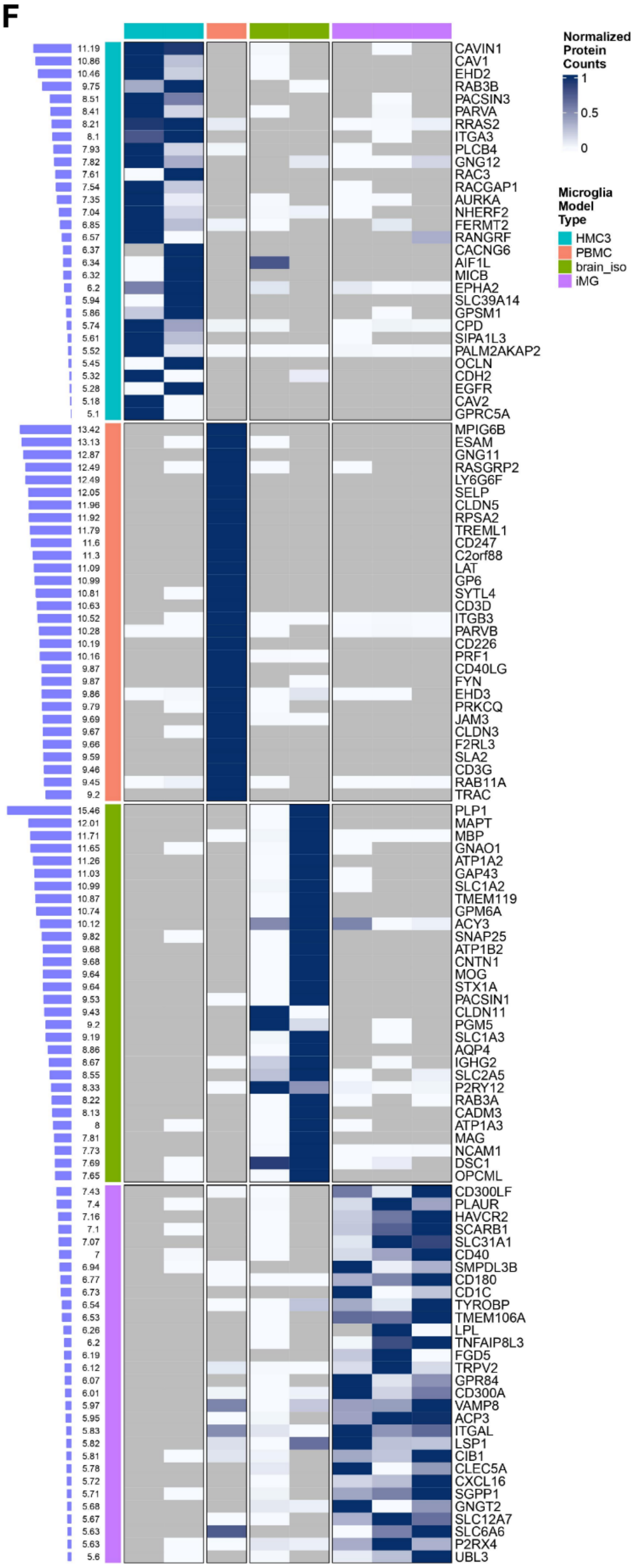


**Supplemental Figure 1. Generation of proteomics data, GWAS analysis and *in silico* analysis for identification of microglia-specific candidate proteins. A. Scheme of workflow for the generation of proteomics from freshly isolated human microglia samples.** Microglia from 38 individuals (women, n= 7; men, n= 4) and different brain regions

(BA4 and BA9 (cortex), AWS (Anterior Watershed), TH (Thalamus), SC (*superior colliculus*), H (Hippocampus), TNC (trigeminal nuclear complex)) were isolated using manual dissociation on ice, followed by magnetic-associated cell sorting (MACS) for CD11b, followed by flow-cytometry associated cell sorting (FASC) for CD11b+CD45+ cells and subsequently subjected to shotgun proteomics analysis. A total of 3658 proteins were identified. **B. PCA plot depicting the distribution of samples and their disease-status identity. Each dot represents a brain tissue sample derived from one donor. Color indicates the nature of the sample** (Control = grey; Epilepsy = orange; ALS (Amyotrophic Lateral Sclerosis) = purple; MS (Multiple Sclerosis) = green; AD (Alzheimer's disease) = red). **C. Expression of selected candidate proteins in HMC3 microglia.** To assess the expression of the selected candidate proteins for the development of a microglia-enriched CITE-Seq panel, the expression of the selected 91 candidate proteins was assessed in a previously generated proteomics dataset of HMC3 microglia. The list of proteins depict the candidates identified to be expressed in HMC3 microglia. **D. Expression of selected candidate proteins in human iPSC-derived microglia.** To assess the expression of the selected candidate proteins for the development of a microglia-enriched CITE-Seq panel, the expression of the selected 91 candidate proteins was assessed in two previously generated proteomics datasets of two independent human iPSC-derived microglia protocols (iMG II: Sher lab protocol[1]; iMG III: Sproul protocol[2]). The list of proteins depict the candidates identified to be expressed in the two independent iPSC-derived microglia preparations[1,2].**E. Venn diagram depicting the expression of assessed protein candidates across the different model systems of human microglia (HMC3, iMG II, iMG III) and freshly isolated human microglia.** Numbers indicate the number of shared proteins expressed/present across the different preparations/model systems of human microglia. **F. Heatmap indicating the expression of the Top 30 proteins expressed across different microglia model types including HMC3 microglia (turquoise), PBMCs (red), freshly isolated human microglia (green) and iMG (iPSC-derived human microglia, purple).** Each row represents a protein, each column a replicate of the different sample types: HMC3 microglia (turquoise), PBMCs (red), freshly isolated human microglia (green) and iMG (iPSC-derived human microglia, purple). Expression is depicted as normalized protein counts, ranging from high (blue) to low (white). Bars on the side including numbers, depict log2 fold difference derived from data imputed with the estimated limit of detection for each dataset.

### Supplementary Figure 2

A   Candidate expression in sc-RNA Seq data I [Cain et al. 2023]

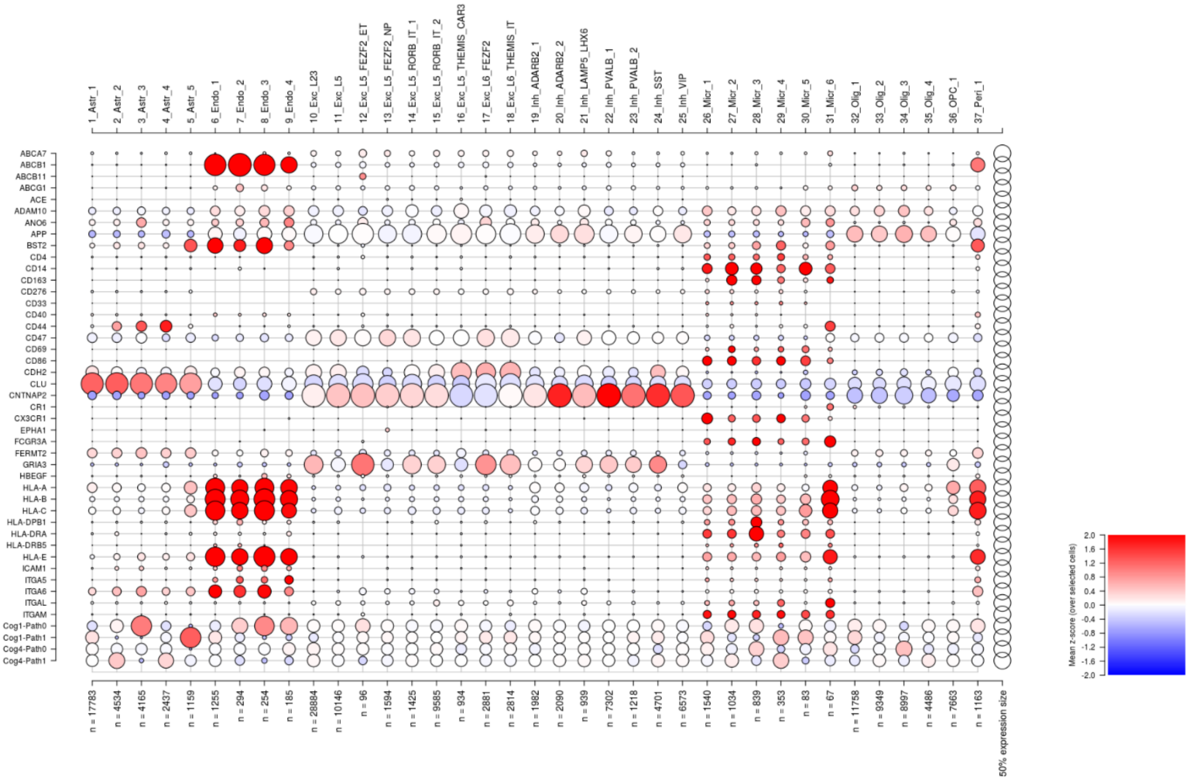


A   Candidate expression in sn-RNA Seq data II [Cain et al. 2023]

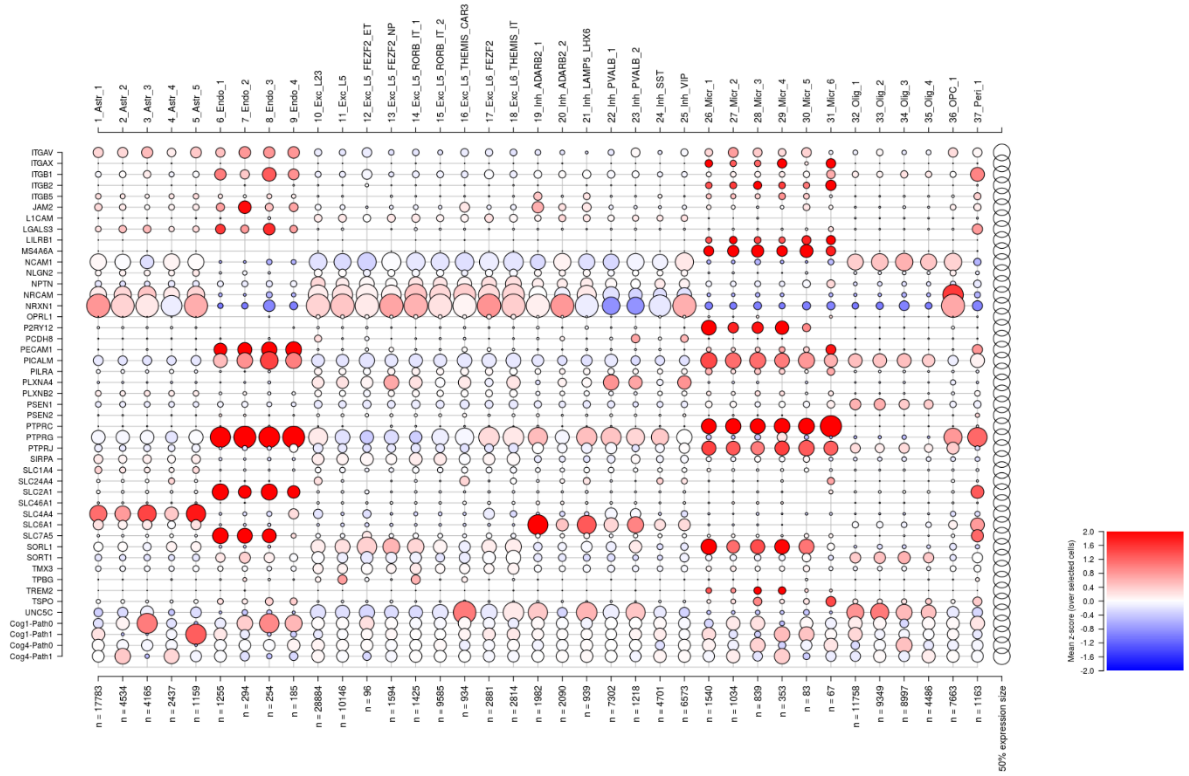

B Candidate expression in sc-RNA Seq data [Tuddenham et al. 2024]

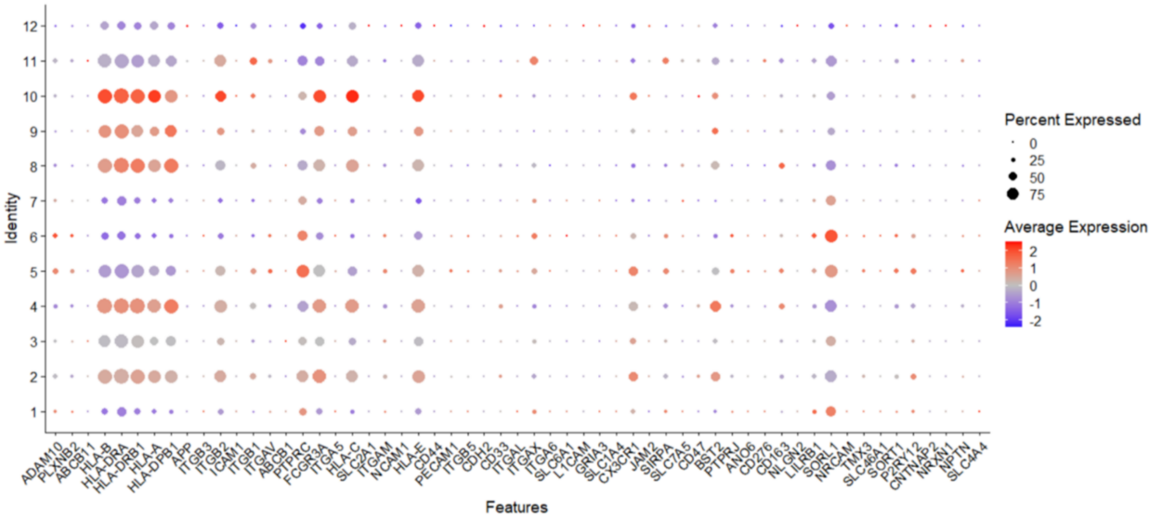


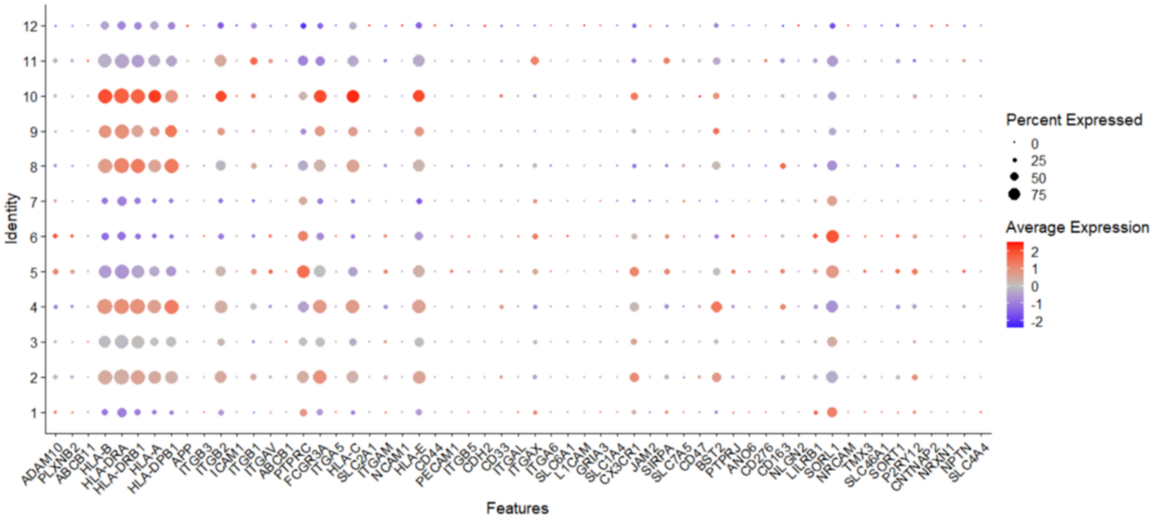

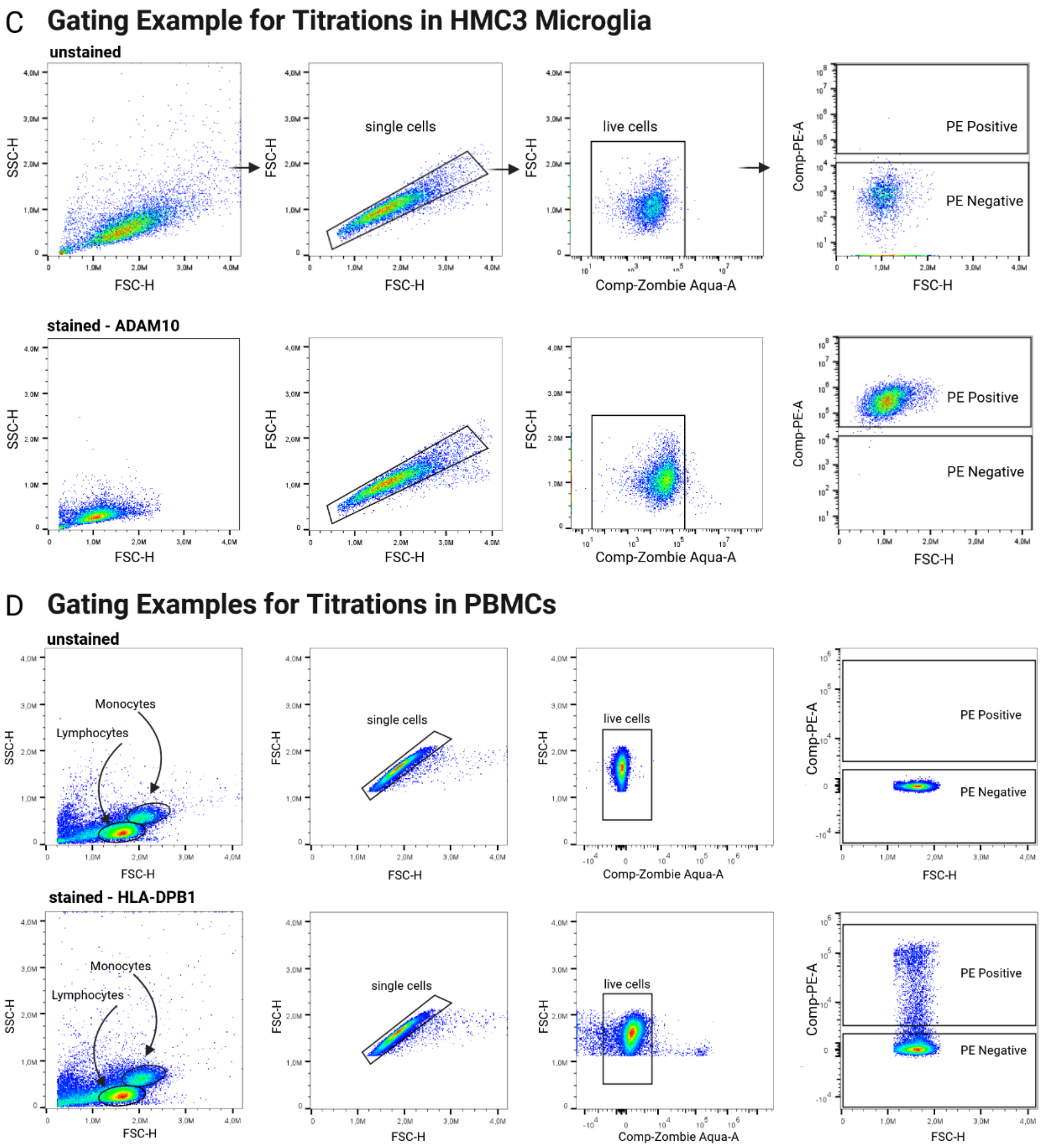

C Gating Example for Titrations in HMC3 Microglia
unstained
SSC-H
FSC-H
single cells
FSC-H
live cells
Comp-Zombie Aqua-A
Comp-PE-A
PE Positive
PE Negative
stained - ADAM10
D Gating Examples for Titrations in PBMCs
unstained
Monocytes
Lymphocytes
stained - HLA-DPB1

E Isotype Controls on HMC3 Microglia and PBMCs

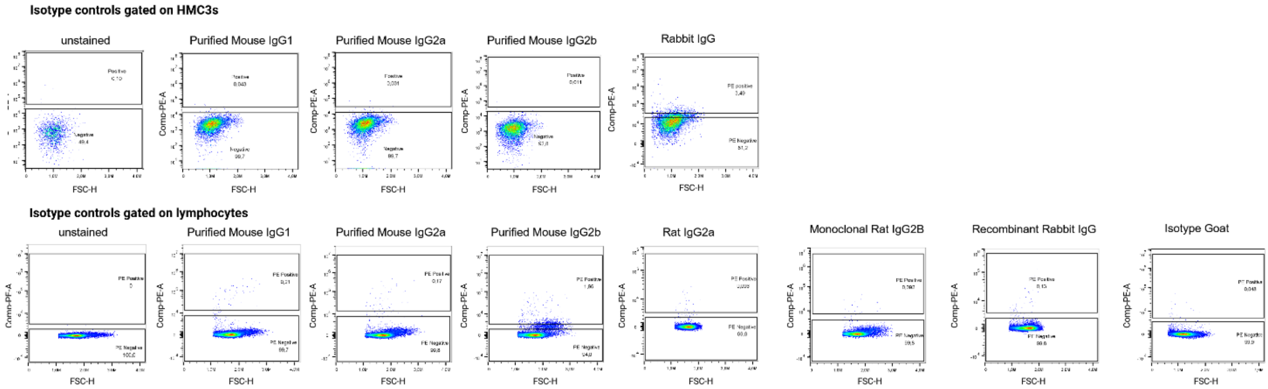


F Secondary Antibody Controls on HMC3 Microglia and PBMCs

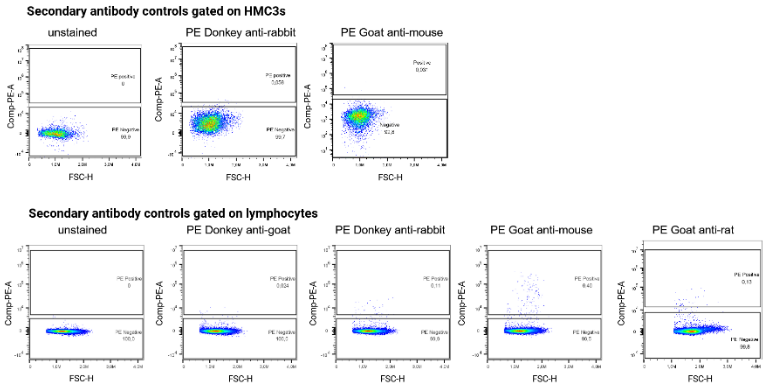


G Titrations in HMC3 microglia

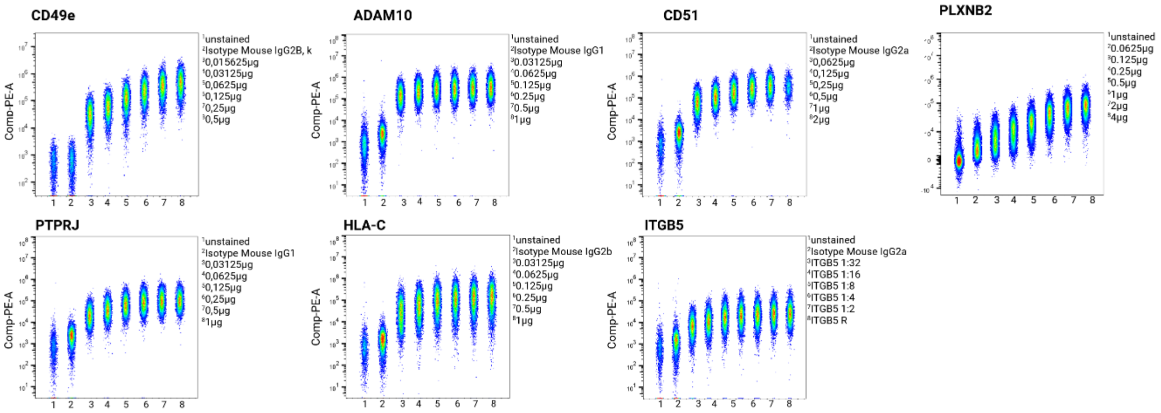

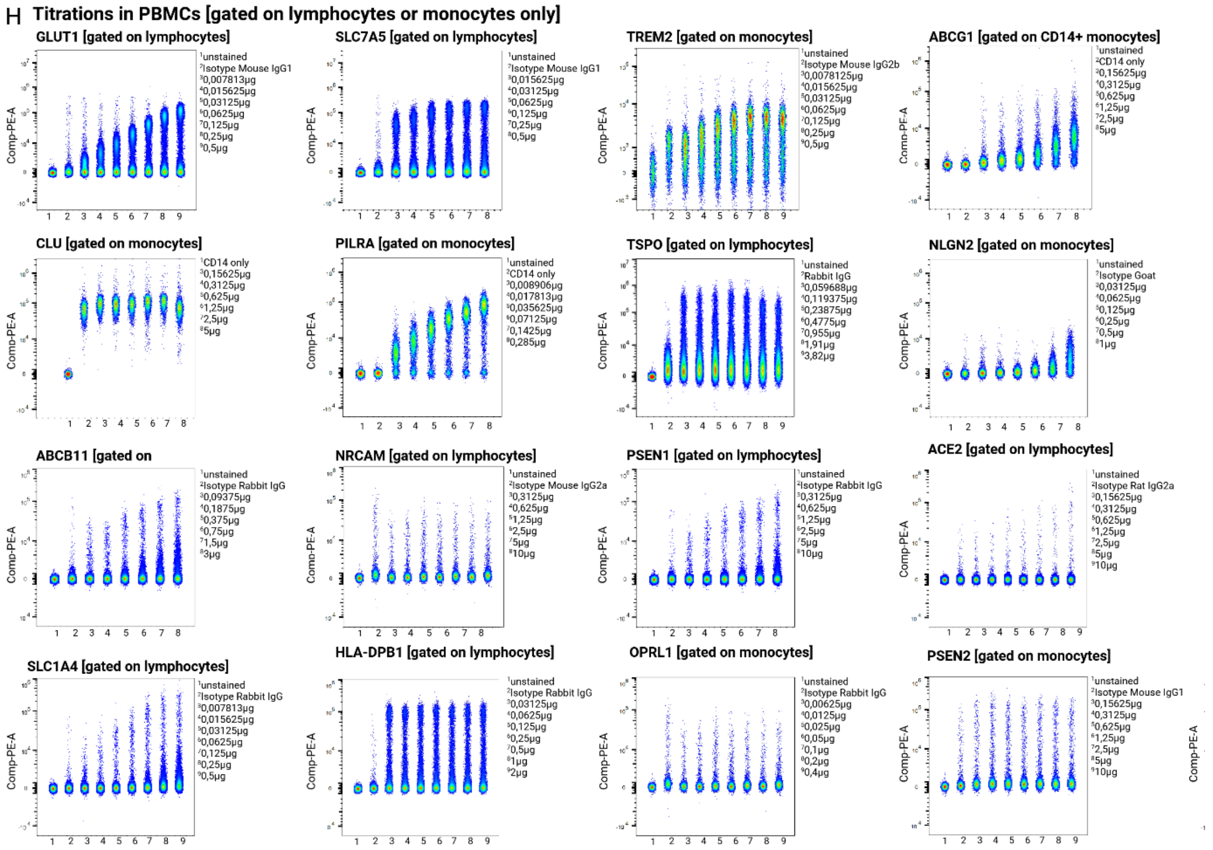


**Supplementary Figure 2. A. Expression of selected candidate proteins in different brain cell types in a selected single-nucleus RNA-Sequencing dataset of the dorsolateral prefrontal cortex (DLPFC) tissue from a structured subgroup of 24 well-characterized individuals[3].** Upper x-axis depicts cell subtype, lower x-axis depicts number of cells in each given cell subtype, y-axis depicts candidate protein in alphabetical order. Circle size depicts number of cells expressing any given protein in percent, colors depict levels of expression (red=high; blue=low). Legend depicts mean Z-score of protein expression over selected cells. **B. Expression of selected candidate proteins in previously defined human microglial subtypes 1-12 from a selected single-cell RNA-Sequencing dataset freshly isolated human microglia from 74 donors[4].** X-axis depicts protein candidates in alphabetical order, y-axis depicts human microglial subsets. Circle size depicts percent of expression, color depicts average expression ranging from high (=red) to low (=blue). **C. Gating example for titrations in HMC3 microglia.** Upper panel – gating strategy for unstained HMC3 microglia. Cells were detected by plotting FSC-H vs. SSC-H. Further gating was performed by plotting selected cell population in FSC-H vs. FSC-H in order to select single cells. Single cells were then plotted for live cells staining (Zombie Aqua-A) vs. FSC-H in order to select live cells (Zombie Aqua-A-negative cells). Live cells were then plotted using FSC-H and PE-A as all markers were purchased in a PE-conjugated format. PE-negative and PE-positive gates were subsequently determined. Lower panel shows example of HMC3 microglia stained for ADAM10, showing a 100% pos. population upon ADAM10 staining. **D. Gating example for titrations in PBMCs (Peripheral Blood Mononuclear Cells).** Upper panel – gating strategy for unstained PBMCs. Cells were detected by plotting FSC-H vs. SSC-H (indicated are monocyte and lymphocyte population). Further gating was performed by plotting selected cell population in FSC-H vs. FSC-H in order to select single cells. Single cells were then plotted for live cells staining (Zombie Aqua-A) vs. FSC-H in order to select live cells (Zombie Aqua-A-negative cells). Live cells were then plotted using FSC-H and PE-A as all markers were purchased in a PE-conjugated format. PE-negative and PE-positive gates were subsequently determined. Lower panel shows example of PBMCs stained for HLA-DPB1, showing a strongly positive, but also still negative population upon HLA-DPB1 staining. **E. Isotype controls on HMC3 microglia and PBMCs. Upper panel:** following gating of single, viable HMC3 microglia and setting the gate for unstained vs. stained HMC3 microglia as described in A., Isotype controls were tested for their expression on HMC3 microglia by plotting FSC-H against PE-A as all Isotypes were also conjugated to PE. Isotype controls tested included: purified mouse IgG1, purified mouse IgG2a, purified mouse IgG2b, Rabbit IgG. **Lower panel:** following gating of single, viable PBMCs, and setting the gate for unstained vs. stained PBMCs as described in B., Isotype controls were tested for their expression on PBMCs by plotting FSC-H against PE-A as all Isotypes were also conjugated to PE. Isotype controls tested included: purified mouse IgG1, purified mouse IgG2a, purified mouse IgG2b, Rat IgG2a, Rat IgG2b, Rabbit IgG and Goat Isotype. **F. Secondary antibody controls on HMC3 microglia and PBMCs. Upper panel:** following gating of single, viable HMC3 microglia and setting the gate for unstained vs. stained HMC3 microglia as described in A., secondary antibody controls were tested for their expression on HMC3 microglia by plotting FSC-H against PE-A as all secondary antibodies were also conjugated to PE. Secondary antibodies tested included: PE-donkey anti-rabbit and PE goat anti-mouse. **Lower panel:** following gating of single, viable PBMCs, and setting the gate for unstained vs. stained PBMCs as described in B., secondary antibody controls were tested for their expression on PBMCs by plotting FSC-H against PE-A as all secondary antibodies were also conjugated to PE. Secondary antibodies tested included: PE donkey anti-goat, PE donkey anti-rabbit, PE goat anti-mouse and PE goat anti-rabbit. **G. Overview of titrations in HMC3 microglia.** Titration results for HMC3 microglia. To evaluate all titration results for the antibody candidates titrated in HMC3 microglia, the PE-positive population of all samples (unstained[1], Isotype control[2], 1:32[3], 1:16[4], 1:8[5], 1:4[6], 1:2[7], recommended concentration[8] ) were concatenated and plotted based on their Comp-PE-A values. Titrations for all antibodies, for which the titrations could be completed in HMC3 microglia are shown, including: CD49e, ADAM10, CD51, PLXNB2, PTPRJ, HLA-C, ITGB5. **H.** Titration results for PBMCs. Examples for titration results for PBMCs. To evaluate all titration results for the antibody candidates titrated in PBMCs, the PE-positive population of all samples (unstained[1], Isotype control[2], 1:32[3], 1:16[4], 1:8[5], 1:4[6], 1:2[7], recommended concentration[8], doubled recommended concentration (2R)[9]) gated on lymphocytes and monocytes were concatenated and plotted based on their Comp-PE-A values. Titrations for all antibodies, for which the titrations could be completed in PBMCs are shown, including:GLUT1, SLC7A5, TREM2, ABCG1, CLU, PILRA, TSPO, NLGN2, ABCB11, NRCAM, PSEN1, ACE2, SLC1A4, HLA-DPB1, OPRL1, PSEN2, SORT1.

**Supplementary Figure 3**

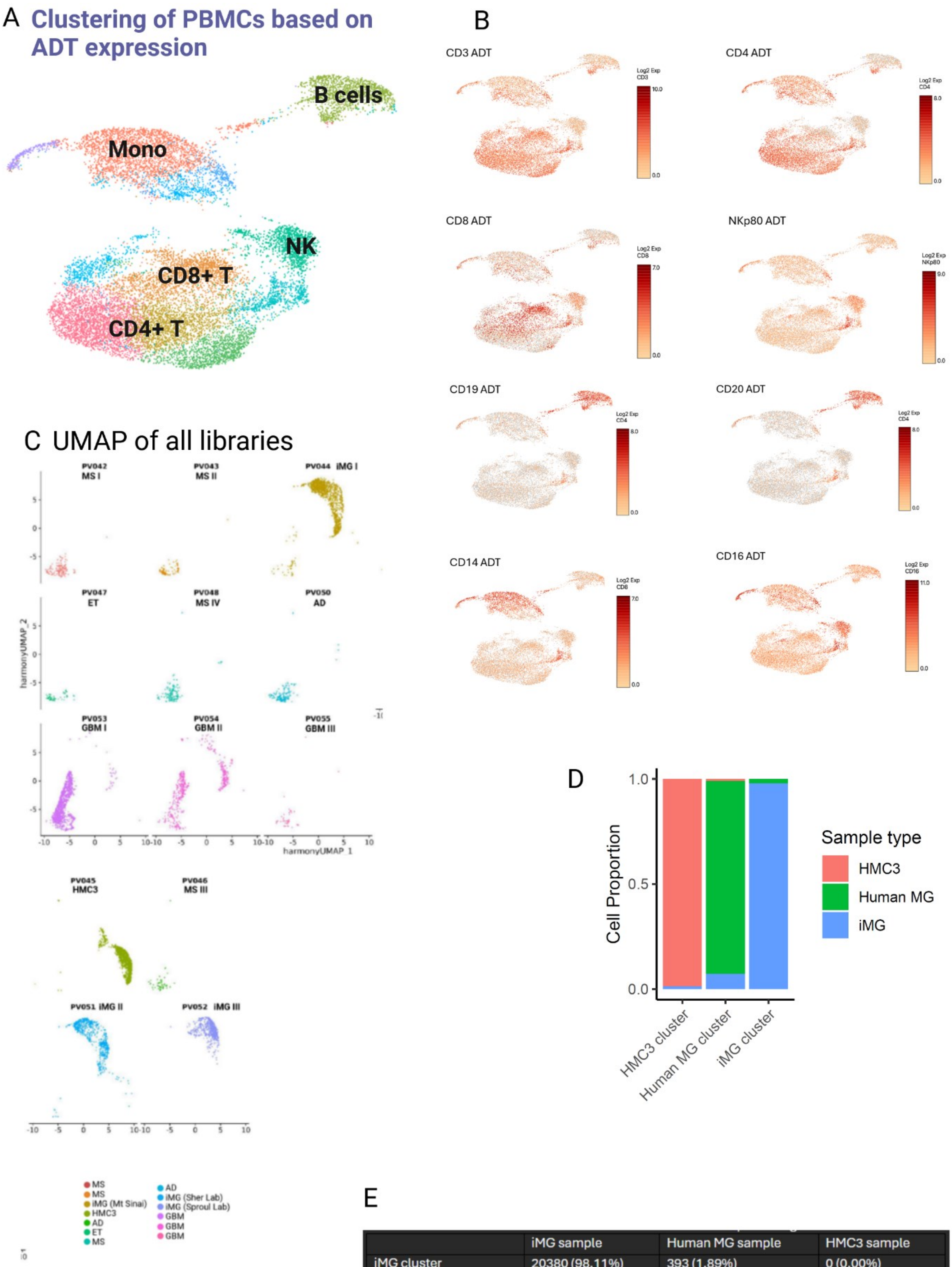


E

| | iMG sample | Human MG sample | HMC3 sample |
|---|---|---|---|
| iMG cluster | 20380 (98.11%) | 393 (1.89%) | 0 (0.00%) |
| Human MG cluster | 1158 (6.56%) | 16405 (92.87%) | 102 (0.58%) |
| HMC3 cluster | 48 (0.54%) | 0 (0.00%) | 8802 (99.46%) |

**Supplementary Figure 3. A. UMAP shows clustering of PBMC (Peripheral Blood Mononuclear Cell) samples based on ADT (Antibody-Derived Tag) expression.** Each cluster is labelled with its respectively identified cell type, based on the expression of protein markers present in the TS-A Universal cocktail for PBMCs which was mixed with the 17 microglia-specific antibodies. B cells are depicted in green, Monocytes (Mono) in orange, NK cells (NK) in turquoise, CD8+ T cells in brown and CD4+ T cells in red/olive/light green. **B.** Expression of main PBMC cell type markers across the PBMC-sample-derived clusters. Depicted are ADT expression patterns for CD3, CD4, CD8, NKp80, CD19, CD20, CD14 and CD16. **C.** UMAPs for all included CITE-Seq libraries from the discovery dataset (PV042-PV055) including their disease-status are depicted (MS = red, orange, turquoise; iMG = olive, blue, light purple; HMC3 = dark green; AD = green, dark turquoise; ET = light green; GBM =purple, pink, dark pink). **D. Bar chart representing cell proportions from different sample types** (HMC3 = red; Human MG = green; iMG = blue) across the different clusters – HMC3 cluster, Human MG cluster, iMG cluster. **E. Table representing cell numbers and percentage of cells for each cell type** (iMG, Human MG, HMC3) across the three different clusters – iMG cluster, HMC3 cluster, Human MG cluster).

**Supplementary Figure 4**

A

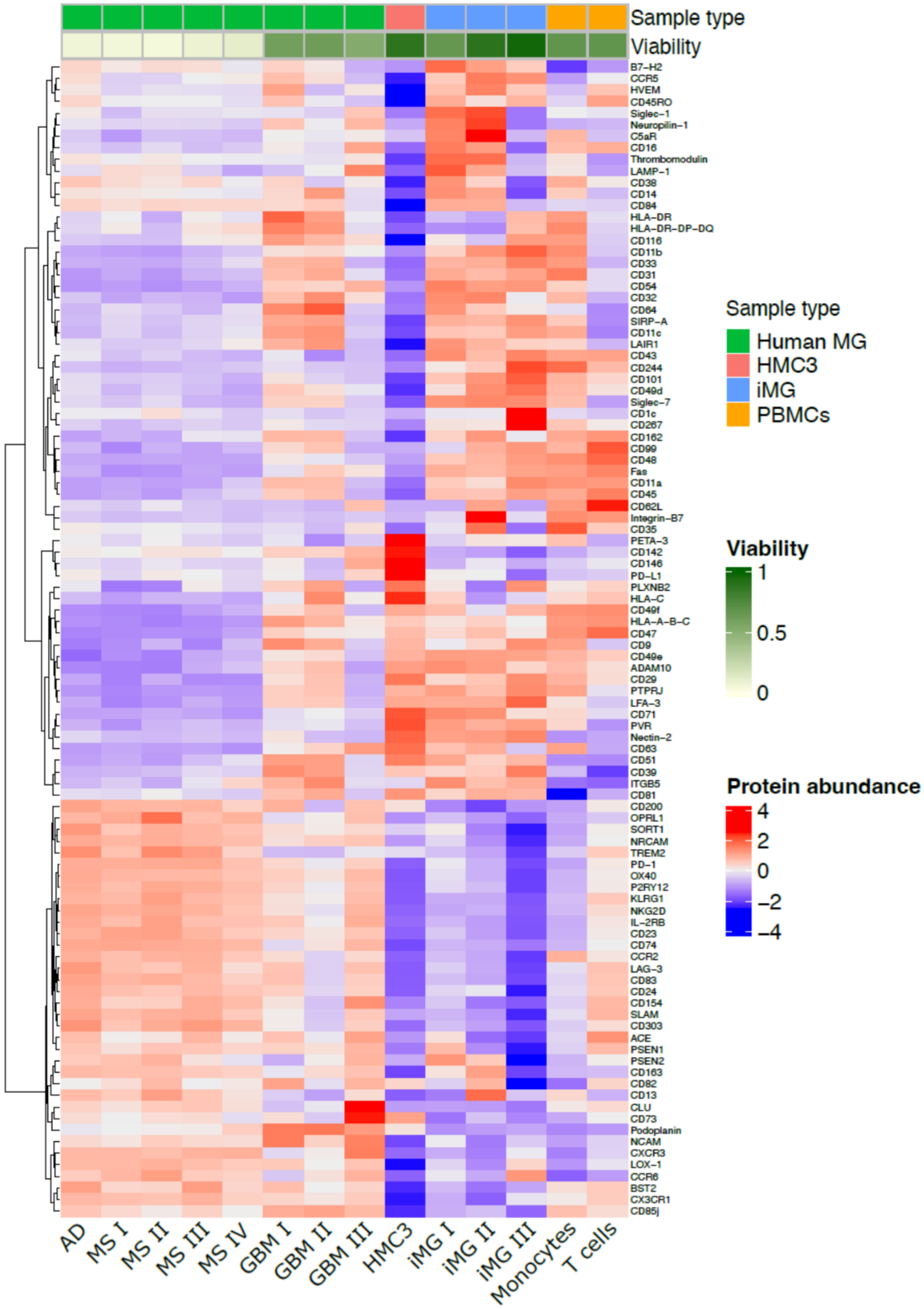

**Supplementary Figure 4. Expression of all CITE-Seq ADTs across different preparations of freshly isolated human microglia and microglia model systems. A. Heatmap depicting the expression of microglia-specific ADTs (Antibody-derived tags) across different samples of human microglia and model systems.** Rows depict the different antibody candidates while columns depict the different samples included in the analysis, including freshly isolated microglia from postmortem samples of an AD (Alzheimer's disease) patient, ET (Essential tremor) patient, MS patients I-IV (Multiple sclerosis), microglia isolated from surgical tissue derived from three GBM (Glioblastoma multiforme) patients (green), HMC3 microglia (red), three different preparations of iPSC-derived human microglia (iMG; blue) from different laboratories and PBMCs derived from 4 different individuals (here depicted are only data for monocytes and T cells; yellow). Cell viability in % ranging from 0% (white) to 100% (green) is indicated on top of each sample together with classification of the sample type (green: freshly isolated human microglia samples; red: HMC3 microglia; blue: iPSC-derived human microglia; yellow: PBMCs, extracted are data for monocytes and T cells). Protein abundance is depicted in colors ranging from red (high) to low (blue).

### Supplementary Figure 5

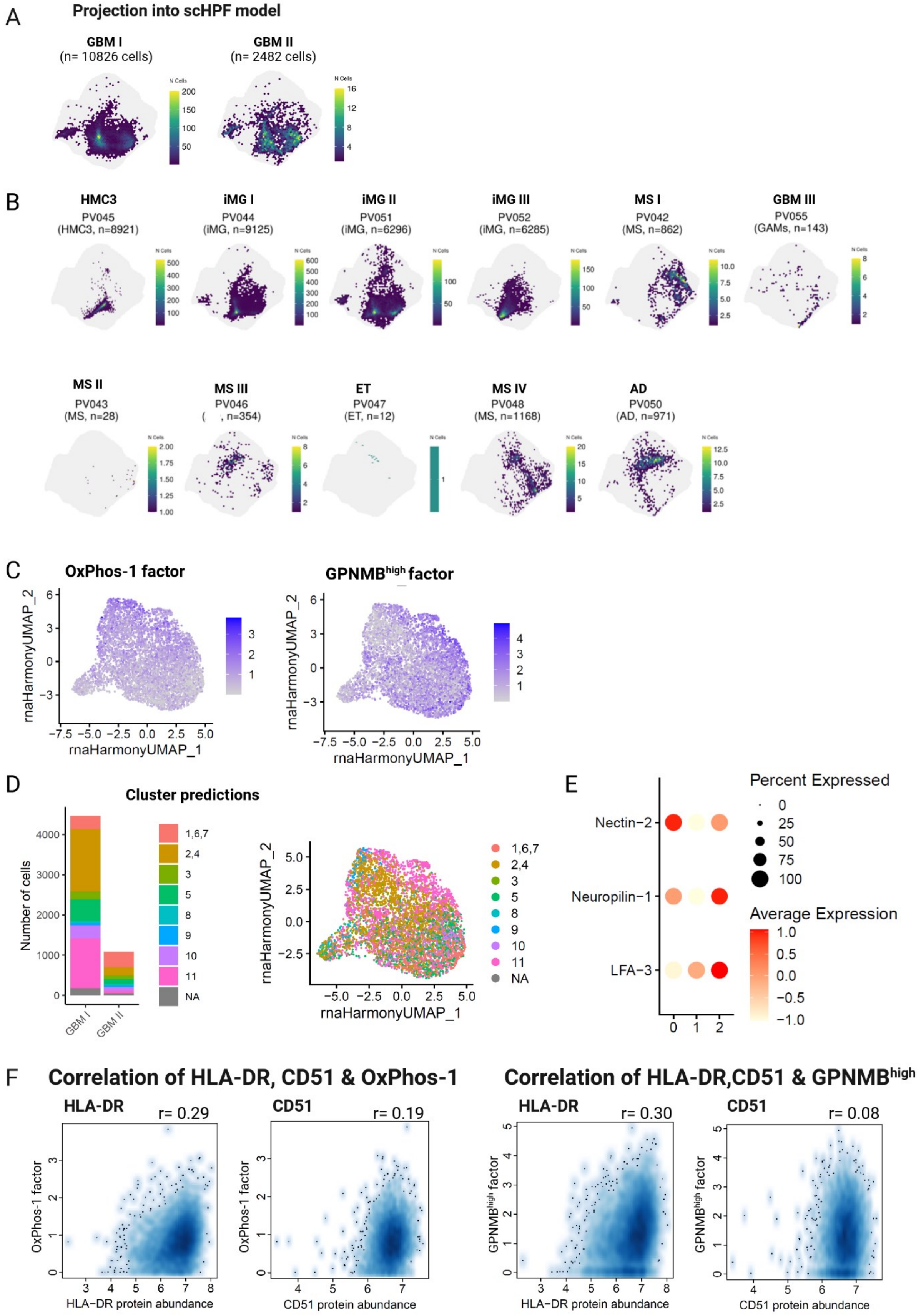

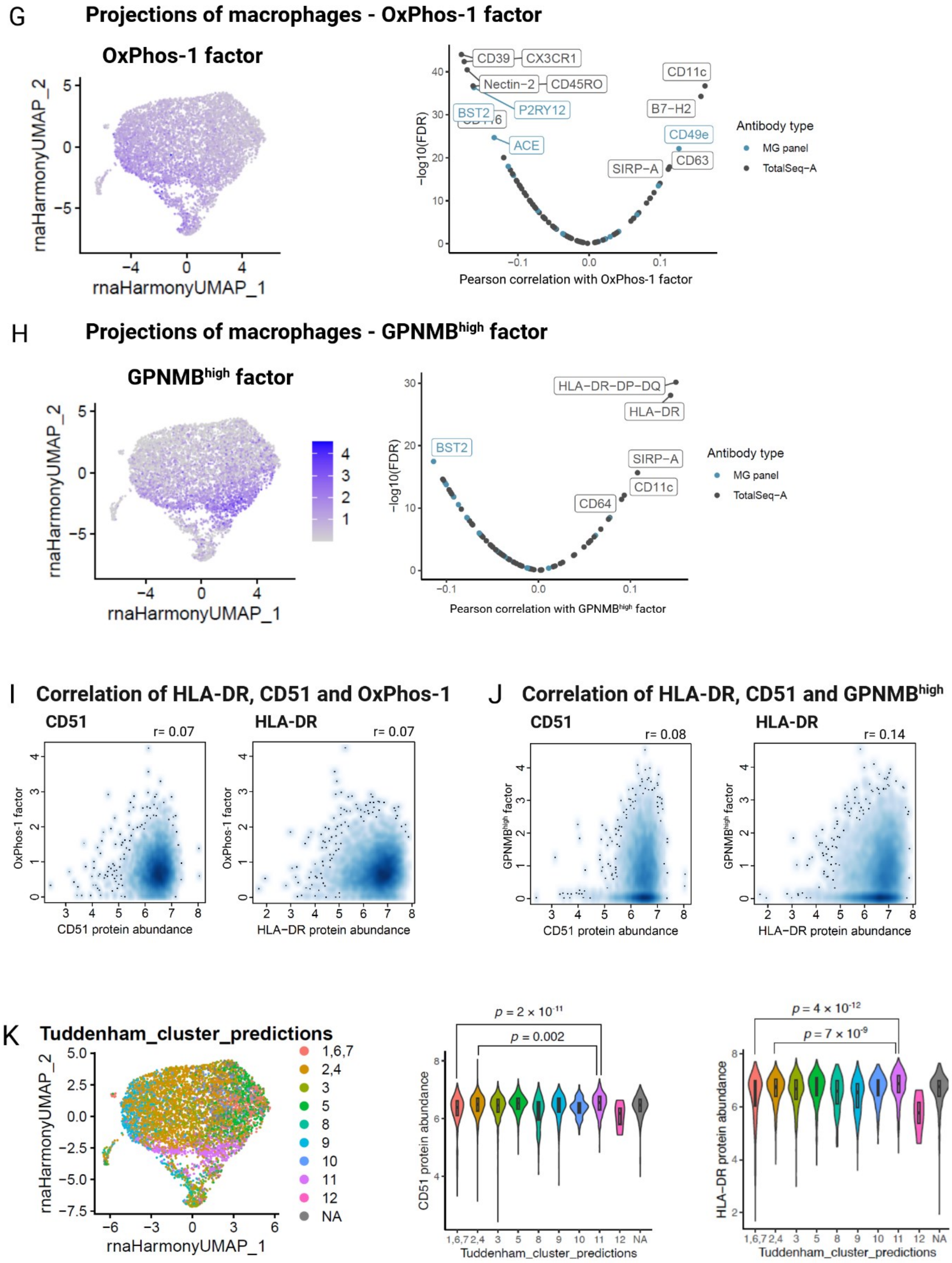

G
Projections of macrophages - OxPhos-1 factor
OxPhos-1 factor
rnaHarmonyUMAP_1
rnaHarmonyUMAP_2
CD39
CX3CR1
Nectin-2
CD45RO
BST2
P2RY12
ACE
CD11c
B7-H2
CD49e
CD63
SIRP-A
−log10(FDR)
Pearson correlation with OxPhos-1 factor
Antibody type
MG panel
TotalSeq-A
H
Projections of macrophages - GPNMB^high factor
GPNMB^high factor
HLA-DR-DP-DQ
HLA-DR
BST2
SIRP-A
CD11c
CD64
Pearson correlation with GPNMB^high factor
I
Correlation of HLA-DR, CD51 and OxPhos-1
CD51
r= 0.07
HLA-DR
r= 0.07
OxPhos-1 factor
CD51 protein abundance
HLA-DR protein abundance
J
Correlation of HLA-DR, CD51 and GPNMB^high
CD51
r= 0.08
HLA-DR
r= 0.14
GPNMB^high factor
K
Tuddenham_cluster_predictions
1,6,7
2,4
3
5
8
9
10
11
12
NA
p = 2 × 10^-11
p = 0.002
p = 4 × 10^-12
p = 7 × 10^-9
CD51 protein abundance
HLA-DR protein abundance
Tuddenham_cluster_predictions

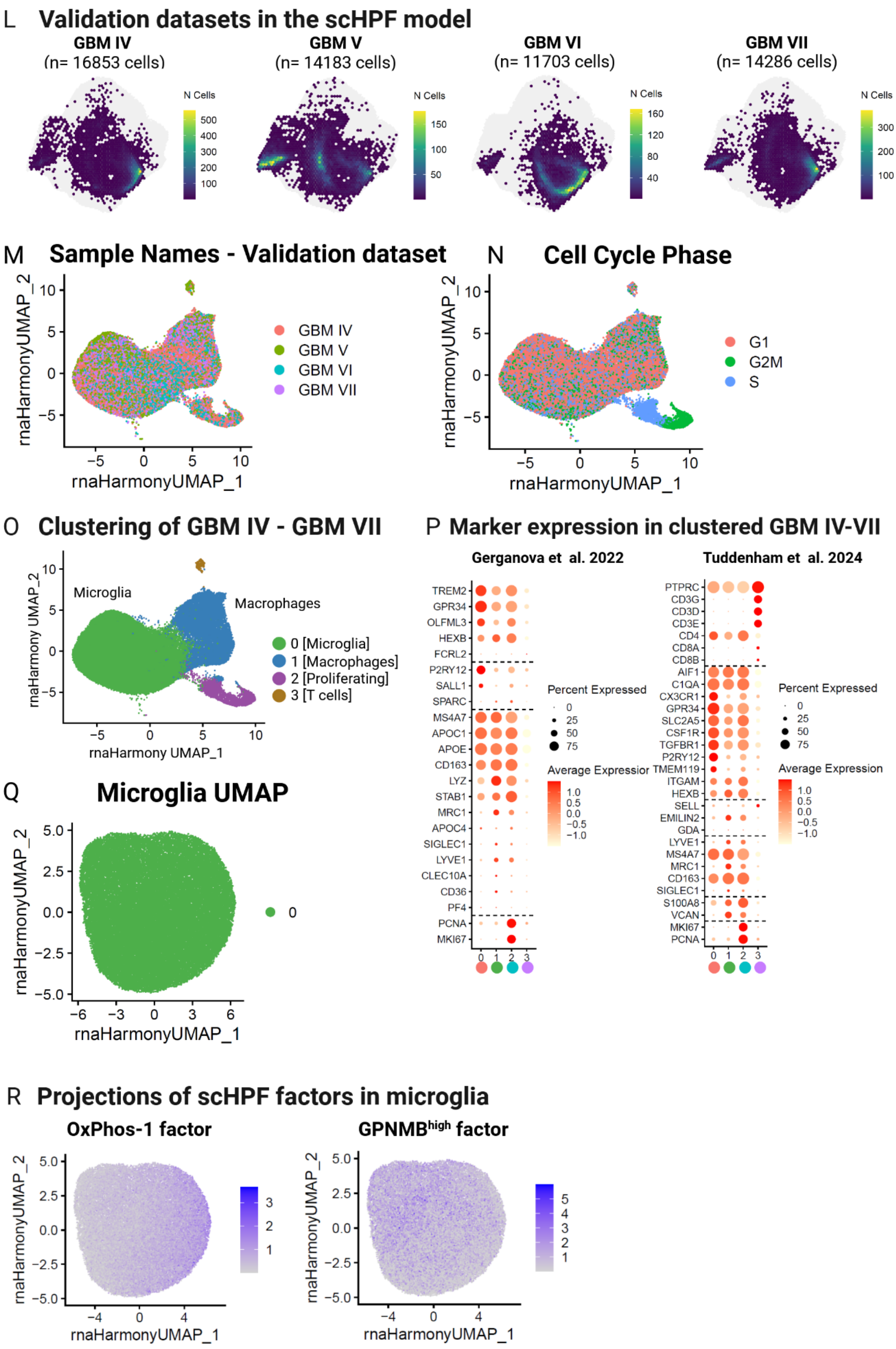
L Validation datasets in the scHPF model
GBM IV
(n= 16853 cells)
GBM V
(n= 14183 cells)
GBM VI
(n= 11703 cells)
GBM VII
(n= 14286 cells)
N Cells
M Sample Names - Validation dataset
GBM IV
GBM V
GBM VI
GBM VII
rnaHarmonyUMAP_1
rnaHarmonyUMAP_2
N Cell Cycle Phase
G1
G2M
S
O Clustering of GBM IV - GBM VII
Microglia
Macrophages
0 [Microglia]
1 [Macrophages]
2 [Proliferating]
3 [T cells]
rnaHarmony UMAP_1
rnaHarmony UMAP_2
P Marker expression in clustered GBM IV-VII
Gerganova et al. 2022
TREM2
GPR34
OLFML3
HEXB
FCRL2
P2RY12
SALL1
SPARC
MS4A7
APOC1
APOE
CD163
LYZ
STAB1
MRC1
APOC4
SIGLEC1
LYVE1
CLEC10A
CD36
PF4
PCNA
MKI67
Tuddenham et al. 2024
PTPRC
CD3G
CD3D
CD3E
CD4
CD8A
CD8B
AIF1
C1QA
CX3CR1
GPR34
SLC2A5
CSF1R
TGFBR1
P2RY12
TMEM119
ITGAM
HEXB
SELL
EMILIN2
GDA
LYVE1
MS4A7
MRC1
CD163
SIGLEC1
S100A8
VCAN
MKI67
PCNA
Percent Expressed
Average Expression
Q Microglia UMAP
rnaHarmonyUMAP_1
rnaHarmonyUMAP_2
R Projections of scHPF factors in microglia
OxPhos-1 factor
GPNMBhigh factor
rnaHarmonyUMAP_1
rnaHarmonyUMAP_2

S

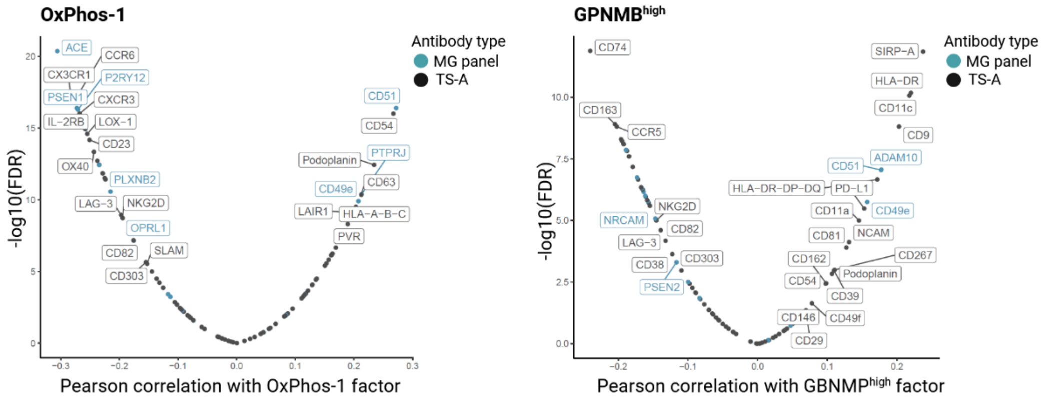


T **Projections of macrophages - OxPhos-1 factor**

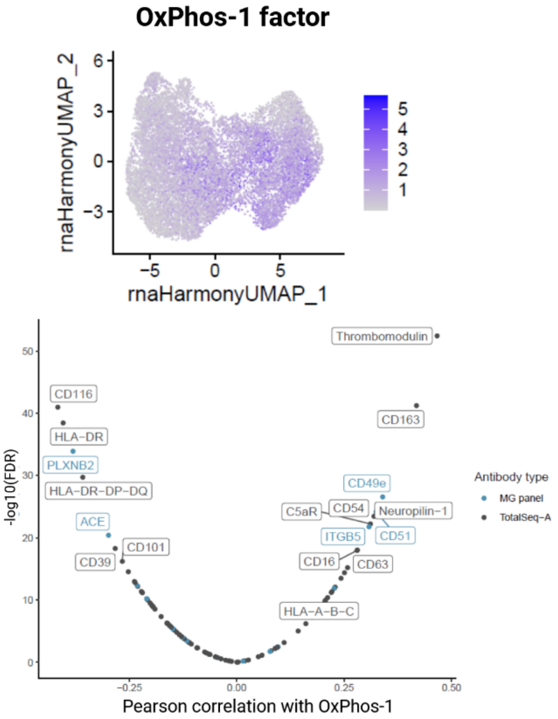


U **Projections of macrophages - GPNMB$^{high}$ factor**

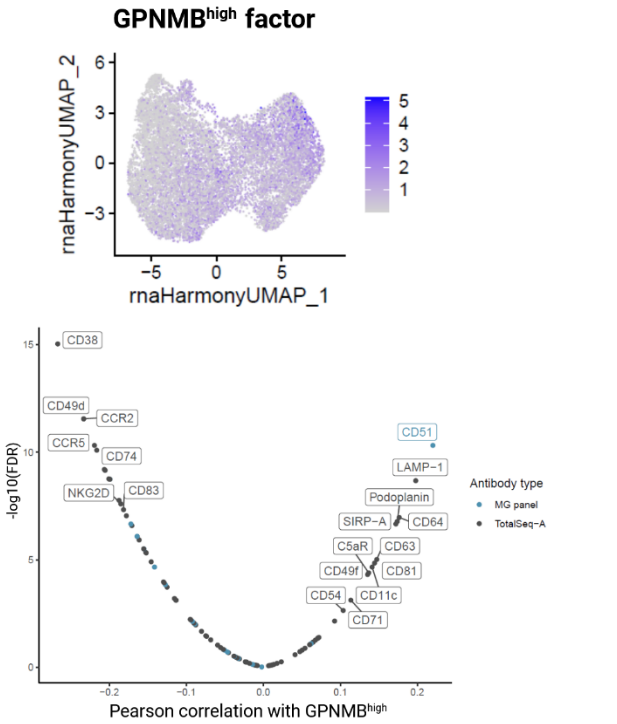

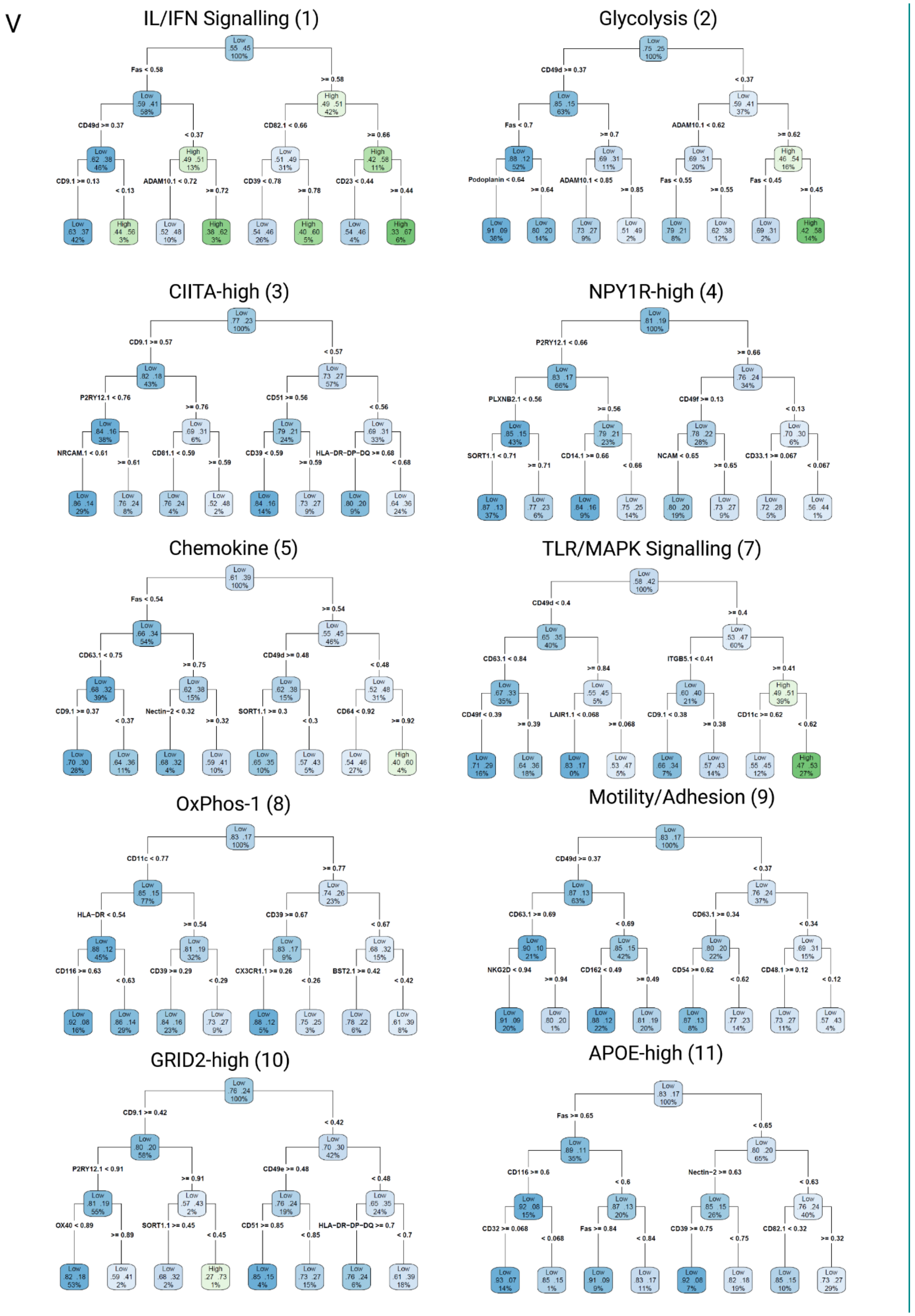

V
IL/IFN Signalling (1)
Fas < 0.58
>= 0.58
CD49d >= 0.37
< 0.37
CD82.1 < 0.66
>= 0.66
CD9.1 >= 0.13
< 0.13
ADAM10.1 < 0.72
>= 0.72
CD39 < 0.78
>= 0.78
CD23 < 0.44
>= 0.44
Glycolysis (2)
CD49d >= 0.37
< 0.37
Fas < 0.7
>= 0.7
ADAM10.1 < 0.62
>= 0.62
Podoplanin < 0.64
>= 0.64
ADAM10.1 < 0.85
>= 0.85
Fas < 0.55
>= 0.55
Fas < 0.45
>= 0.45
CIITA-high (3)
CD9.1 >= 0.57
< 0.57
P2RY12.1 < 0.76
>= 0.76
CD51 >= 0.56
< 0.56
NRCAM.1 < 0.61
>= 0.61
CD81.1 < 0.59
>= 0.59
CD39 < 0.59
>= 0.59
HLA-DR-DP-DQ >= 0.68
< 0.68
NPY1R-high (4)
P2RY12.1 < 0.66
>= 0.66
PLXNB2.1 < 0.56
>= 0.56
CD49f >= 0.13
< 0.13
SORT1.1 < 0.71
>= 0.71
CD14.1 >= 0.66
< 0.66
NCAM < 0.65
>= 0.65
CD33.1 >= 0.067
< 0.067
Chemokine (5)
Fas < 0.54
>= 0.54
CD63.1 < 0.75
>= 0.75
CD49d >= 0.48
< 0.48
CD9.1 >= 0.37
< 0.37
Nectin-2 < 0.32
>= 0.32
SORT1.1 >= 0.3
< 0.3
CD64 < 0.92
>= 0.92
TLR/MAPK Signalling (7)
CD49d < 0.4
>= 0.4
CD63.1 < 0.84
>= 0.84
ITGB5.1 < 0.41
>= 0.41
CD49f < 0.39
>= 0.39
LAIR1.1 < 0.068
>= 0.068
CD9.1 < 0.38
>= 0.38
CD11c >= 0.62
< 0.62
OxPhos-1 (8)
CD11c < 0.77
>= 0.77
HLA-DR < 0.54
>= 0.54
CD39 >= 0.67
< 0.67
CD116 >= 0.63
< 0.63
CD39 >= 0.29
< 0.29
CX3CR1.1 >= 0.26
< 0.26
BST2.1 >= 0.42
< 0.42
Motility/Adhesion (9)
CD49d >= 0.37
< 0.37
CD63.1 >= 0.69
< 0.69
CD63.1 >= 0.34
< 0.34
NKG2D < 0.94
>= 0.94
CD162 < 0.49
>= 0.49
CD54 >= 0.62
< 0.62
CD48.1 >= 0.12
< 0.12
GRID2-high (10)
CD9.1 >= 0.42
< 0.42
P2RY12.1 < 0.91
>= 0.91
CD49e >= 0.48
< 0.48
OX40 < 0.89
>= 0.89
SORT1.1 >= 0.45
< 0.45
CD51 >= 0.85
< 0.85
HLA-DR-DP-DQ >= 0.7
< 0.7
APOE-high (11)
Fas >= 0.65
< 0.65
CD116 >= 0.6
< 0.6
Nectin-2 >= 0.63
< 0.63
CD32 >= 0.068
< 0.068
Fas >= 0.84
< 0.84
CD39 >= 0.75
< 0.75
CD82.1 < 0.32
>= 0.32

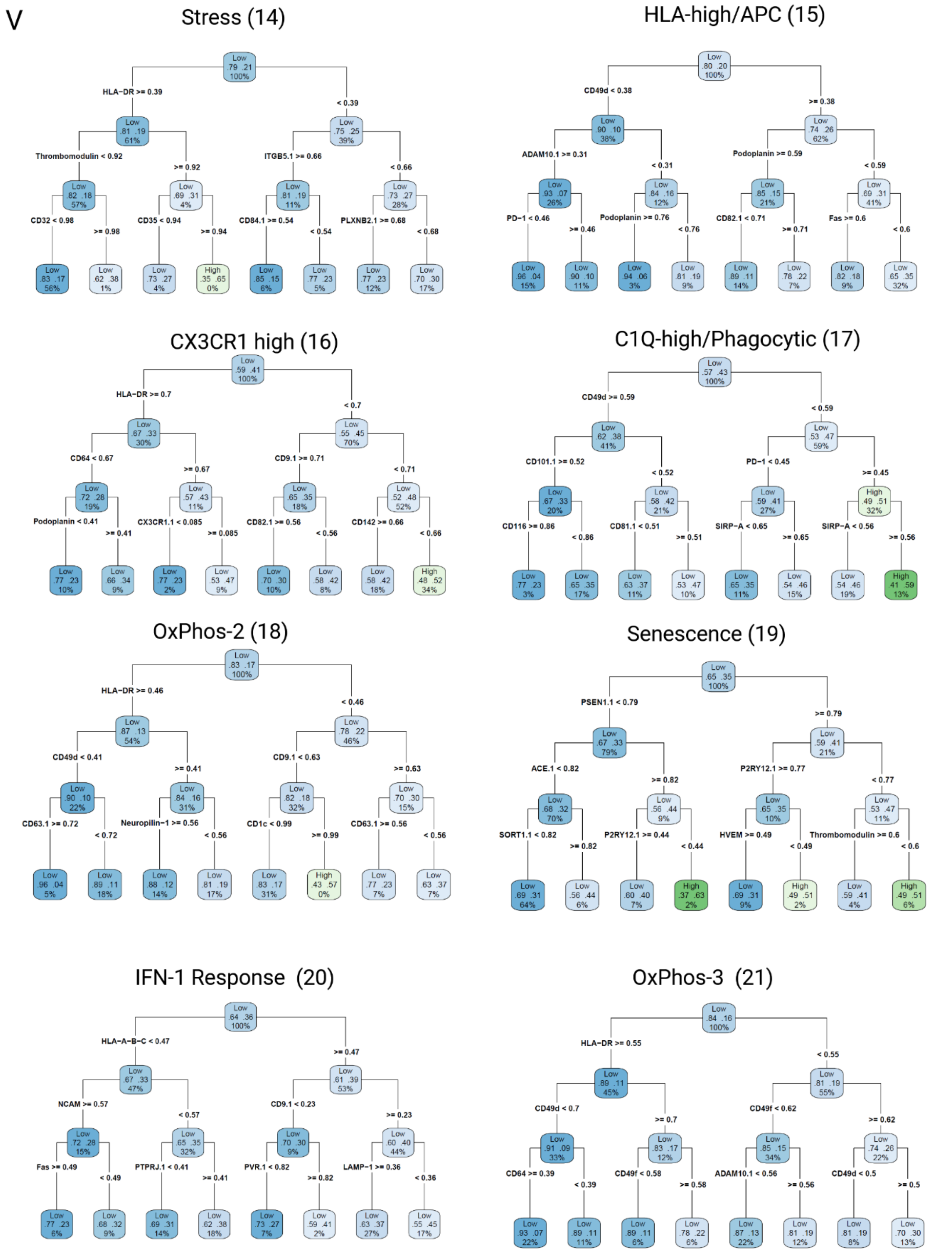
V
Stress (14)
HLA−DR >= 0.39
< 0.39
Thrombomodulin < 0.92
>= 0.92
ITGB5.1 >= 0.66
< 0.66
CD32 < 0.98
>= 0.98
CD35 < 0.94
>= 0.94
CD84.1 >= 0.54
< 0.54
PLXNB2.1 >= 0.68
< 0.68
HLA-high/APC (15)
CD49d < 0.38
>= 0.38
ADAM10.1 >= 0.31
< 0.31
Podoplanin >= 0.59
< 0.59
PD−1 < 0.46
>= 0.46
Podoplanin >= 0.76
< 0.76
CD82.1 < 0.71
>= 0.71
Fas >= 0.6
< 0.6
CX3CR1 high (16)
HLA−DR >= 0.7
< 0.7
CD64 < 0.67
>= 0.67
CD9.1 >= 0.71
< 0.71
Podoplanin < 0.41
>= 0.41
CX3CR1.1 < 0.085
>= 0.085
CD82.1 >= 0.56
< 0.56
CD142 >= 0.66
< 0.66
C1Q-high/Phagocytic (17)
CD49d >= 0.59
< 0.59
CD101.1 >= 0.52
< 0.52
PD−1 < 0.45
>= 0.45
CD116 >= 0.86
< 0.86
CD81.1 < 0.51
>= 0.51
SIRP−A < 0.65
>= 0.65
SIRP−A < 0.56
>= 0.56
OxPhos-2 (18)
HLA−DR >= 0.46
< 0.46
CD49d < 0.41
>= 0.41
CD9.1 < 0.63
>= 0.63
CD63.1 >= 0.72
< 0.72
Neuropilin−1 >= 0.56
< 0.56
CD1c < 0.99
>= 0.99
CD63.1 >= 0.56
< 0.56
Senescence (19)
PSEN1.1 < 0.79
>= 0.79
ACE.1 < 0.82
>= 0.82
P2RY12.1 >= 0.77
< 0.77
SORT1.1 < 0.82
>= 0.82
P2RY12.1 >= 0.44
< 0.44
HVEM >= 0.49
< 0.49
Thrombomodulin >= 0.6
< 0.6
IFN-1 Response (20)
HLA−A−B−C < 0.47
>= 0.47
NCAM >= 0.57
< 0.57
CD9.1 < 0.23
>= 0.23
Fas >= 0.49
< 0.49
PTPRJ.1 < 0.41
>= 0.41
PVR.1 < 0.82
>= 0.82
LAMP−1 >= 0.36
< 0.36
OxPhos-3 (21)
HLA−DR >= 0.55
< 0.55
CD49d < 0.7
>= 0.7
CD49f < 0.62
>= 0.62
CD64 >= 0.39
< 0.39
CD49f < 0.58
>= 0.58
ADAM10.1 < 0.56
>= 0.56
CD49d < 0.5
>= 0.5

V

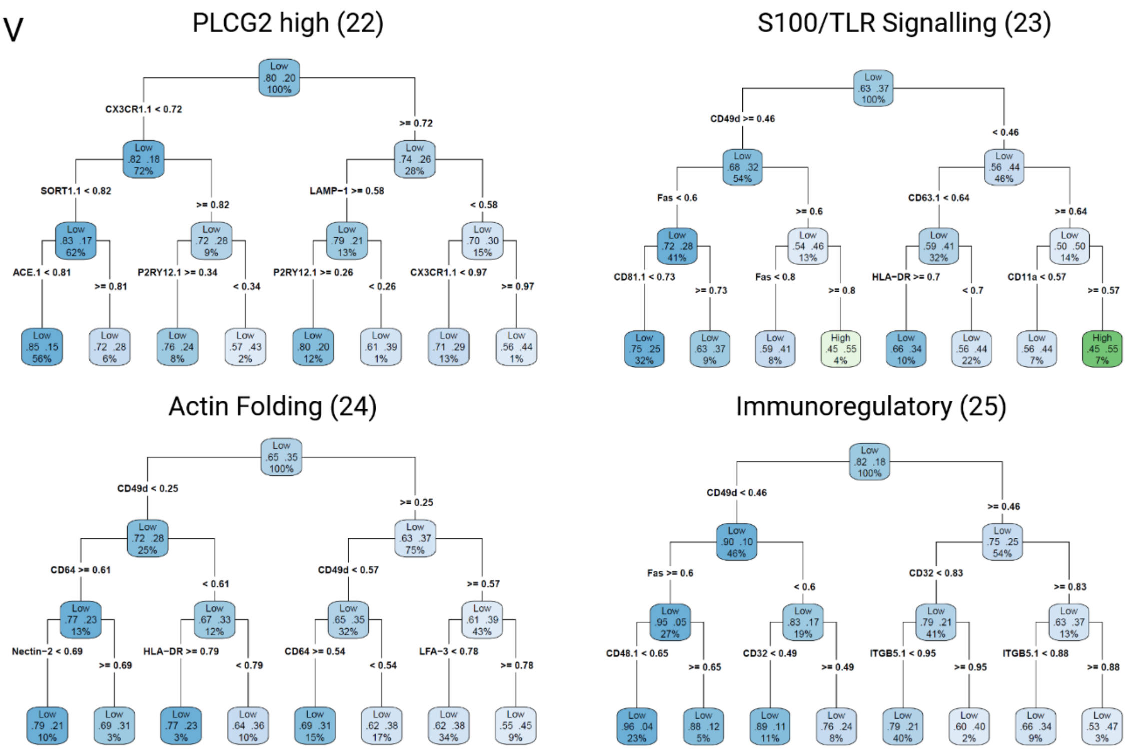
PLCG2 high (22)
CX3CR1.1 < 0.72
>= 0.72
SORT1.1 < 0.82
>= 0.82
LAMP−1 >= 0.58
< 0.58
ACE.1 < 0.81
>= 0.81
P2RY12.1 >= 0.34
< 0.34
P2RY12.1 >= 0.26
< 0.26
CX3CR1.1 < 0.97
>= 0.97
S100/TLR Signalling (23)
CD49d >= 0.46
< 0.46
Fas < 0.6
>= 0.6
CD63.1 < 0.64
>= 0.64
CD81.1 < 0.73
>= 0.73
Fas < 0.8
>= 0.8
HLA−DR >= 0.7
< 0.7
CD11a < 0.57
>= 0.57
Actin Folding (24)
CD49d < 0.25
>= 0.25
CD64 >= 0.61
< 0.61
CD49d < 0.57
>= 0.57
Nectin−2 < 0.69
>= 0.69
HLA−DR >= 0.79
< 0.79
CD64 >= 0.54
< 0.54
LFA−3 < 0.78
>= 0.78
Immunoregulatory (25)
CD49d < 0.46
>= 0.46
Fas >= 0.6
< 0.6
CD32 < 0.83
>= 0.83
CD48.1 < 0.65
>= 0.65
CD32 < 0.49
>= 0.49
ITGB5.1 < 0.95
>= 0.95
ITGB5.1 < 0.88
>= 0.88

W

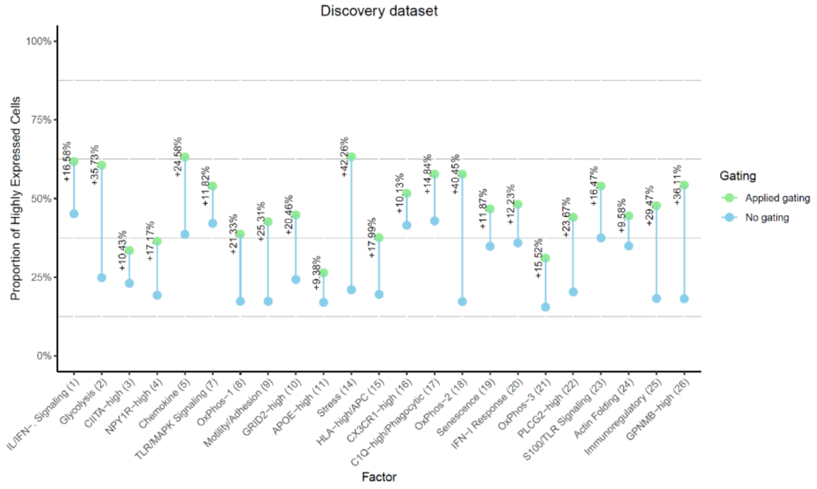
Discovery dataset
Proportion of Highly Expressed Cells
100%
75%
50%
25%
0%
Gating
Applied gating
No gating
IL/IFN-. Signaling (1)
Glycolysis (2)
CIITA-high (3)
NPY1R-high (4)
Chemokine (5)
TLR/MAPK Signaling (7)
OxPhos-1 (8)
Motility/Adhesion (9)
GRID2-high (10)
APOE-high (11)
Stress (14)
HLA-high/APC (15)
CX3CR1-high (16)
C1Q-high/Phagocytic (17)
OxPhos-2 (18)
Senescence (19)
IFN-I Response (20)
OxPhos-3 (21)
PLCG2-high (22)
S100/TLR Signaling (23)
Actin Folding (24)
Immunoregulatory (25)
GPNMB-high (26)
Factor

X

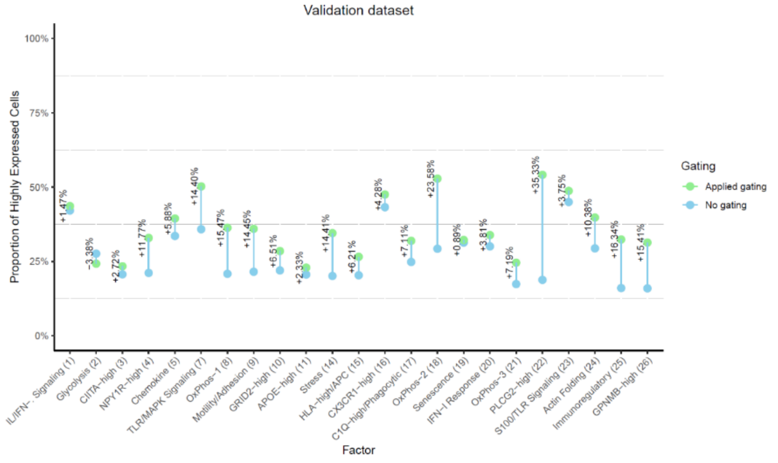
Validation dataset
Proportion of Highly Expressed Cells
100%
75%
50%
25%
0%
Gating
Applied gating
No gating
IL/IFN-. Signaling (1)
Glycolysis (2)
CIITA-high (3)
NPY1R-high (4)
Chemokine (5)
TLR/MAPK Signaling (7)
OxPhos-1 (8)
Motility/Adhesion (9)
GRID2-high (10)
APOE-high (11)
Stress (14)
HLA-high/APC (15)
CX3CR1-high (16)
C1Q-high/Phagocytic (17)
OxPhos-2 (18)
Senescence (19)
IFN-I Response (20)
OxPhos-3 (21)
PLCG2-high (22)
S100/TLR Signaling (23)
Actin Folding (24)
Immunoregulatory (25)
GPNMB-high (26)
Factor

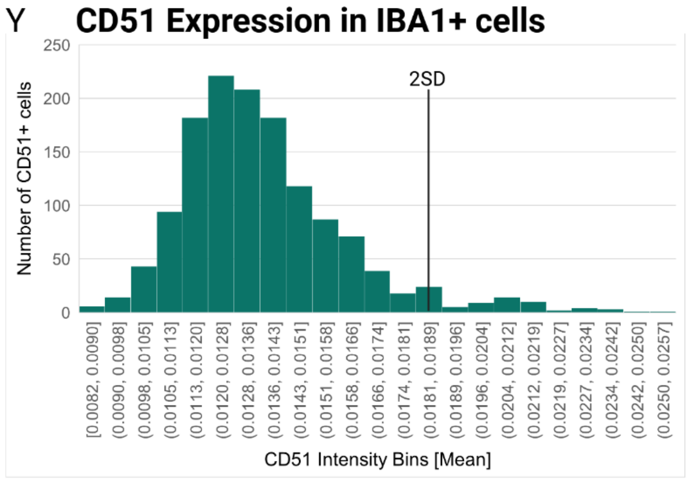


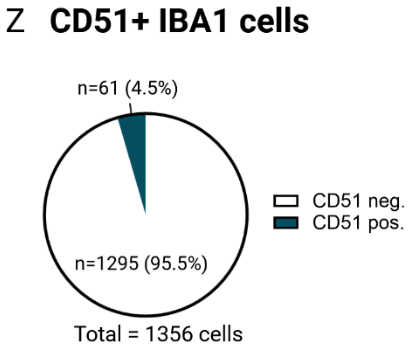


**Supplementary Figure 5. Projection into scHPF and sc-RNA Seq datasets of human microglia. A. Projection of GBM I and GBM II into the scHPF model[5]**. All cells for GBM I (n= 10826) and GBM II (n= 2482) were projected into the UMAP of 441,088 cells derived from 161 donors. The grey shape represents the UMAP of the ROSMAP data. Each cell is depicted as a single dot. Cell density is indicated by colors, ranging from low (purple) to high (yellow). **B. Projection of all samples into the scHPF model.** All cells for all samples – HMC3 microglia (n= 8921), iMG I (n= 9125), iMG II (n= 6296), iMG III (n= 6285), MS I (n= 862), MS II (n= 28), MS III (n= 354), MS IV (n= 1168), AD (n= 971), ET (n= 12), GBM III (n= 143) - were projected into the UMAP of microglia derived from 161 donors[5]. The grey shape represents the UMAP of the ROSMAP data. Each cell is depicted as a single dot. Cell density is indicated by colors, ranging from low (purple) to high (yellow). **C. Projections of scHPF factors into isolated microglia from GBM I and II.** Using Cluster 1 identified as microglia, previously identified scHPF factors[5] were projected into the microglia cluster to assess for their expression. Two factors, OxPhos-1 (left) and GBNMP$^{high}$ (right) were highly expressed in complementary parts of the cluster 1 UMAP. **D. Plot depicting the number of cells for GBM I and GBM II samples predicted per cluster (1-11) from the Tuddenham et al. study[4]. Left:** Each bar indicates the number of cells per sample (GBM I, GBM II) predicted to fall into 11 different clusters as defined by Tuddenham et al. following projection analysis. NA = not available. Red = clusters 1,6,7; brown: clusters 2,4; olive: cluster 3; green: cluster 5; turquoise: cluster 8; blue: cluster 9; violet: cluster 10; pink: cluster 11 – DAM-like. **Right:** Plot shows the projection of 11 microglia clusters/subsets defined from human microglial sc-RNA Seq data by Tuddenham et al. [4] into the cluster 1 defined as microglia from GBMI and GBMII samples. Each dot indicates a single cell, each color indicates a different microglial cluster as defined as clusters 1-11 by Tuddenham et al. [4](red: cluster group 1,6,7; brown: clusters 2,4; olive: cluster 3; green: cluster 5; turquoise: cluster 8; blue: cluster 9; violet: cluster 10; pink: cluster 11 – DAM-like). **E.** Expression levels of selected protein candidates Nectin-2, Neuropilin-1 and LFA-3 across the three identified clusters 0,1,2 in joint samples GBM I and GBM II. Each row represents a protein, each column represents the expression in each of the clusters 0,1,2 in the form of a dot. The size of each dot represents the percentage expression ranging from very small (0%) to large (100%). The color of each dot indicates the average expression level ranging from yellow (low) to red (high). **F. Correlation of HLA-DR & CD51 and OxPhos-1 (left).** Correlation between HLA-DR (left) and CD51 (right) protein abundance with the OxPhos-1 factor. Darker color represents higher cell density, and outlier cells were shown as individual dots. Pearson correlation coefficient is depicted at the top of the plot (HLA-DR - r=0.29; CD51 – r= 0.19).**. Correlation of HLA-DR & CD51 and GBNMP$^{high}$ (right).** Correlation between HLA-DR (left) and CD51 (right) protein abundance with the GBNMP$^{high}$ factor. Pearson correlation coefficient is depicted at the top of the plot (HLA-DR - r=0.30; CD51 – r= 0.08). **G-H. Projections of scHPF factors into isolated macrophages from GBM I and II.** Using Cluster 0 identified as macrophages, previously identified scHPF factors[5] were projected into the macrophage cluster to assess for their expression. Two factors, OxPhos-1 (**G**) and GBNMP$^{high}$ (**H**) were highly expressed in complementary parts of the cluster 0 UMAP. The second plot for each of the factors represented (OxPhos-1, GBNMP$^{high}$) shows the Pearson correlation for the CITE-Seq antibodies (microglia (MG) panel – turquoise; TS-A Human Universal Cocktail 1.0 - black) with each of the assessed factors (OxPhos-1 – **G**; GBNMP$^{high}$ - **H**). Correlation for all antibodies is shown in relation to -log10(FDR) values. Antibodies with lowest and highest correlation are labelled in different , colors depending on their origin - turquoise (microglia-specific panel) or black (TS-A Universal cocktail). **I. Correlation of HLA-DR & CD51 and OxPhos-1 in macrophages.** Correlation between CD51 (left) and HLA-DR (right) protein abundance with the OxPhos- 1 factor. Darker color represents higher cell density, and outlier cells were shown as individual dots. Pearson correlation coefficient is depicted at the top of the plot (CD51 – r= 0.07; HLA-DR - r=0.07). **J. Correlation of HLA-DR & CD51 and GBNMP$^{high}$ in macrophages.** Correlation between CD51 (left) and HLA-DR (right) protein abundance with the GBNMP$^{high}$ factor. Pearson correlation coefficient is depicted at the top of the plot (CD51 – r= 0.08; HLA-DR - r=0.14). **K. Projection into sc-RNA Seq dataset of human microglia (Tuddenham et al. 2024[4]) and expression of HLA-DR and CD51 across the sc-RNA Seq microglial clusters.** The plot on the left shows the projection of 11 microglia clusters/subsets defined from human microglial sc-RNA Seq data by Tuddenham et al. [4] into the cluster 0 defined as macrophages from GBMI and GBMII samples. Each dot indicates a single cell, each color indicates a different microglial cluster as defined as clusters 1-11 by Tuddenham et al. [4](red: cluster group 1,6,7; brown: clusters 2,4; olive: cluster 3; green: cluster 5; turquoise: cluster 8; blue: cluster 9; violet: cluster 10; pink: cluster 11 – DAM-like). Violin plots show the protein abundance of selected proteins –CD51 (left) and HLA-DR (right) across the 11 different clusters. Internal box plot indicates the first, second, and third quartiles of protein abundance. For statistical analysis, the Wilcoxon Rank Sum test was performed. **L. Projection of GBM IV-VII into the scHPF model[5]**. All cells for GBM IV (n=16853), GBM V (n=14183), GBM VI (n=11703), GBM VII (n= 14286) were projected into the UMAP of 441,088 cells derived from 161 donors. The grey shape represents the UMAP of the ROSMAP data. Each cell is depicted as a single dot. Cell density is indicated by colors, ranging from low (purple) to high (yellow). **M. UMAP of validation dataset consisting of microglia/macrophages freshly isolated from GBM-diagnosed donors.** UMAP consists of microglia isolated from four independent GBM-tumor samples (GBM IV-GBM VII), each represented in a different color (GBM IV= red; GBM V = green; GBMVI = turquoise; GBM VII = purple). Each dot represents a single cell. **N. UMAP of pooled GBM IV-VII samples indicating Cell Cycle Phase for each analyzed cell, represented by different colors (G1 = red; G2M = green; S = blue).** Each dot represents a single cell. **O. Hierarchical clustering of joint cells from GBM IV-VII.** Graph represents hierarchical clustering of joint cells derived from GBM IV-GBM VII samples. Clustering resulted in four clusters, termed 0 (green), 1 (blue), 2 (purple) and 3 (brown). **P. Expression of defined microglia- and macrophage markers across the identified clusters derived from GBM IV- GBM VII.** Marker expression of microglia- and macrophage markers as previously used by Gerganova et. al. [6](left) and Tuddenham et al. [4](right) across the clusters 0,1,2,3 derived from joint GBM IV-GBMVII samples. Plots show markers derived from both datasets and their expression across the different cluster 0,1,2,3. Average expression is indicated by color scheme with high expression in red and low expression in light yellow. Cluster 1 emerges as a microglia-specific cluster. **Q. UMAP representing microglia only following hierarchical clustering analysis and identification of each of the four identified clusters using marker genes previously published by Gerganova et al.[6] and Tuddenham et al.[4]** Each dot represents a single cell.

**R. Projections of scHPF factors into isolated microglia from GBM IV-GBM VII.** Using Cluster 0 identified as microglia, previously identified scHPF factors[5] were projected into the microglia cluster to assess for their expression. Two factors, OxPhos-1 (left) and $GBNMP^{high}$ (right) were highly expressed in complementary parts of the cluster 0 UMAP. **S.** The plots for each of the factors represented (left: OxPhos-1, right: $GBNMP^{high}$) shows the Pearson correlation for the CITE-Seq antibodies (microglia (MG) panel – turquoise; TS-A Human Universal Cocktail 1.0 - black) with each of the assessed factors (OxPhos-1; $GBNMP^{high}$). Correlation for all antibodies is shown in relation to -log10(FDR) values. Antibodies with lowest and highest correlation are labelled in different colors depending on their origin - turquoise (microglia-specific panel) or black (TS-A Universal cocktail). **T-U. Projections of scHPF factors into isolated macrophages from GBM IV- GBM VII.** Using Cluster 1 identified as macrophages, previously identified scHPF factors[5] were projected into the macrophage cluster to assess for their expression. Two factors, OxPhos-1 (**T**) and $GBNMP^{high}$ (**U**) were highly expressed in complementary parts of the cluster 0 UMAP. The second plot for each of the factors represented (OxPhos-1, $GBNMP^{high}$) shows the Pearson correlation for the CITE-Seq antibodies (microglia (MG) panel – turquoise; TS-A Human Universal Cocktail 1.0 - black) with each of the assessed factors (OxPhos-1 – **T**; $GBNMP^{high}$ - **U**). Correlation for all antibodies is shown in relation to -log10(FDR) values. Antibodies with lowest and highest correlation are labelled in different , colors depending on their origin - turquoise (microglia-specific panel) or black (TS-A Universal cocktail). **V. Decision tree diagram for protein-expression-based enrichment of the different scHPF factors[5] in microglia.** Diagram indicates selection of top3 markers for *in silico* sorting of scHPF factors 1-26-expressing microglia (factor number in brackets, factor names on top of each graph), resulting in three top candidate protein markers per scHPF factor. Numbers between 0-1 indicate percentage of cells with either higher or lower expression of each of the markers according to a previously defined threshold. Percentage on the bottom of each info box indicates percentage (%) of total input cells. **W. *In silico* sorting using the top 3 defined markers for each of the 26 defined scHPF factors[5]** using the discovery dataset GBM I-II. X-axis depicts each of the assessed 26 scHPF factors/expression programs, y-axis indicates proportion of highly expressed cells for the assessed factors in percent (%). Blue dots indicate proportion of highly expressing cells for each of the scHPF factors without prior gating, while green dots indicate percentage of highly expressing cells for each of the factors following gating based on the previously defined top 3 protein markers. Numbers between no gating and applied gating datapoints indicate enrichment (%) for any given scHPF factor in the proportion of highly expressed cells. **X. *In silico* sorting using the top 3 defined markers for each of the 26 defined scHPF factors[5]** using the validation dataset GBM IV-VII. X-axis depicts each of the assessed 26 scHPF factors/expression programs, y-axis indicates proportion of highly expressed cells for the assessed factors in percent (%). Blue dots indicate proportion of highly expressing cells for each of the scHPF factors without prior gating, while green dots indicate percentage of highly expressing cells for each of the factors following gating based on the previously defined top 3 protein markers. Numbers between no gating and applied gating datapoints indicate enrichment (%) for any given scHPF factor in the proportion of highly expressed cells. **Y.** Histogram depicting the distribution of CD51 mean intensity bins across the number of IBA1-positive cells. 2 SD (standard deviation) is shown as a cutoff for CD51+ cells. **Z.** Pie chart depicting the number and percentage of CD51-negative (CD51 mean intensity ≤ 2SD; n= 1295) and CD51-positive (≥ 2SD; n= 61) IBA1-positive microglia in stained brain tissue sections. Total number of analyzed cells following outlier removal – n= 1356.
